# On How, and Why, and When, We Grow Old

**DOI:** 10.1101/2022.05.29.493895

**Authors:** Luca Citti, Jessica Su, James S Michaelson

**Affiliations:** School of Computer Science and Electronic Engineering, University of Essex, UK; Department of Biomedical Engineering, Johns Hopkins University, USA; Department of Pathology, Massachusetts General Hospital, USA; Department of Surgery, Massachusetts General Hospital, USA; Department of Pathology, Harvard Medical School, USA

## Abstract

Growth and aging are fundamental features of animal life. The march from fertilization to oblivion comes in enormous variety: days and hundreds of cells for nematodes, decades and trillions of cells for humans.^1-4^ Since Verhulst (1838^5^) proposed the Logistic Equation - exponential growth with countervailing linear decline in rate – biologists have searched for ever better density dependent growth equations,^6-12^ none which accurately capture the relationship between size and time for real animals.^13-15^ Furthermore, while growth and aging run in parallel, whether the relationship is causal has been unknown. Here we show, by examining growth and lifespan in units of numbers of cells, *N*, (*Cellular Phylodynamics*^6^*), that both processes are linked to the same reduction in the fraction of cells dividing, occurring by a simple expression, the Universal Mitotic Fraction Equation*. Lifespan is correlated with an age when fewer than one-in-a-thousand cells are dividing, quantifying the long-appreciated mechanism of aging, the failure of cells to be rejuvenated by dilution with new materials made, and DNA repaired, at mitosis.^24-26^ These observations provide practical mathematical expressions for comprehending, and managing, the challenges of growth and aging, for such tasks as improving the effectiveness of COVID-19 vaccination in the elderly.

## INTRODUCTION

### Cellular Phylodynamics

As we grow older, we grow bigger, and we grow slower, and we grow frailer. We can gain insight into the deeper processes at work by considering growth in units of numbers of cells, ***N***. We have called this approach ***Cellular Phylodynamics***,^6^ building from Stadler, Pybus, and Stumpf,^16^ who have shown how methods developed for studying the populations of organisms in nature can also be put to use in studying the populations of cells within us. Indeed, as we have described previously, such insight can be gained for understanding the growth and creation of the body and its parts,^6^ for understanding the transformation, by somatic mutation, of normal growth into precancerous growth, and then into cancer,^17^ as well as for understanding the spread of cancer cells,^17,^18,19 and putting this knowledge to practical use.^20,21^ Here, we shall focus on how such a ***Cellular Phylodynamic*** approach can give us insight into to aging and its link to growth.

## RESULTS

### Growth

For more than two centuries, it has been appreciated that growth begins exponentially, but then declines in rate, and many equations to capture such density dependent growth have been considered,^6-12^ none of which accurately fit the growth of real animals.^13-15^ Rather than searching for another equation that fits growth, we carried out a ***Cellular Phylodynamic Analysis*** to examine what animal size data, in units of numbers of cells, ***N***, can tell us about the nature of the growth.^6^ To do so, we assembled values of numbers of cells, ***N***, from birth to maturity, for nematodes, frogs, chickens, cows, geese, quail, turkeys, mice, rats, fish, mollusks and humans (FIGURE 1, APPENDIX, and Reference 6). These data revealed, as expected, that at the beginning of life, almost all cells are dividing, displaying the ***Cell Cycle Time, c***. Soon however, the embryo steps back from such exponential growth, gradually at first, but with an accelerating decline in the fraction of cells that are dividing. We could make a rough measure of this mitotic decline by calculating the fraction of cells that would have to divide in order to account for the amount of growth that occurs at each moment in time, a value we call the ***Mitotic Fraction, m. m*** doesn’t take into account cell death, cell size, etc., but we shall see that it’s a good start for addressing the questions at hand. The practical execution of ***m***’s calculation can be a bit involved, but details may be found in the APPENDIX and Reference 6.

**FIGURE 1:**
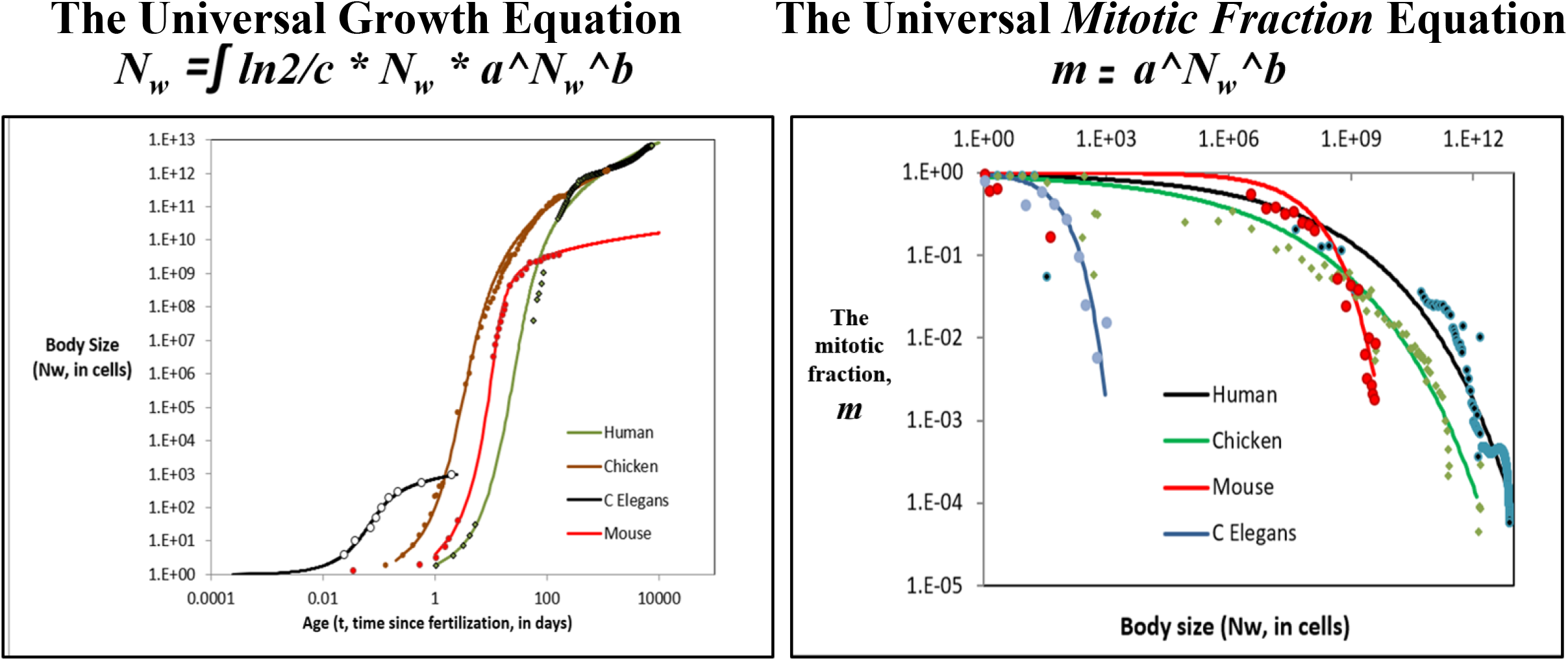
Growth data and the *Universal Growth Equation* and *Universal Mitotic Fraction Equation* for capturing growth from fertilization until maturity. For additional data, see APPENDIX and Reference 6 **Left**: **Growth, in units of numbers of cells, *Nw*, in the whole body, from fertilization until maturity, in units of days, *t***. Datapoints for animal size’s (***N***_***x***_) versus time (***t***), for humans, chickens, ***C elegans*** nematode worms, and mice, and their fit to the numerically integrated form of the ***Universal Growth Equation*** (#2).^6^ **Right**: **Estimate of the decline in the *Mitotic Fraction*, a measure of the fraction of cells dividing that occurs as animals increase in size, as estimated by the *Mitotic Fraction Method*, and captured by the** ***Universal Mitotic Fraction Equation***. Data points for animal size, ***N***_***w***_, in integer units of numbers of cells, vs. the ***Mitotic Fraction, m***, from fertilization, until maturity, for humans, chickens, mice, and ***C elegans*** nematode worms.^6^

Strikingly, when we graphed the ***Mitotic Fraction, m***, versus animal size, ***N***, all ten animals listed above displayed the same relationship:

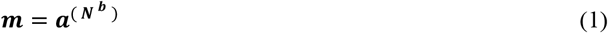

where the ***a*** and ***b*** parameters describe each animal’s specific growth qualities (FIGURE 1, APPENDIX, and Reference 6). We call Equation#1 the ***Universal Mitotic Fraction Equation***, as it describes the decline in the fraction of cells dividing, as measured by the ***Mitotic Fraction, m***, from ∼100% after conception, to very small numbers, as growth slows to adulthood.

Incorporating ***Cell Cycle Time, c***, yields the rate of growth:

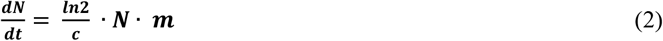

Integration yields the relationship between ***age, t***, and ***size, N***:

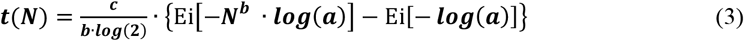

When tested against actual growth data, Equation #3 was found to closely capture the growth, from fertilization until maturity, for each of the species listed above (FIGURE 1, APPENDIX, and Reference 6). Tests of other widely used density dependent growth equations found none to capture actual growth data as well as Equation #3, and it also succeeds, as previous equations did not, in capturing growth over its full span, from the first fertilized cell to adult size (APPENDIX and Reference 6). For these reasons, we call Equation #3 the ***Universal Growth Equation***.

From a purely descriptive standpoint, the ***Universal Growth Equation***’s ***a*** and ***b*** parameters mold the “curviness” of the ***S-shaped*** growth curve, determining the sizes to which animals grow, while ***c*** expands or contracts the “***S*”** like an accordion, determining the speed at which they grow. We don’t know why growth occurs by this ***Universal Growth Equation***, but, curiously, for the idealized case of cells growing in a constant volume, such as an egg or a uterus, little more than the discrete allocation of inhibitory growth factor molecules binding to cells is enough to give rise to the ***Mitotic Fraction*** in the form of the ***Universal Mitotic Fraction Equation***. The mathematical demonstration of this finding can be seen in the APPENDIX and Reference 6, and the values of the ***a*** and ***b*** parameters reflect the thermodynamics of the growth factors at work. The appearance of the ***Cell Cycle Time, c***, in the ***Universal Growth Equation*** is also a pleasant surprise. The length of time that it takes a cell to divide has long been known to reflect genome size,^6,^22 and ***c***’s appearance in the ***Universal Growth Equation***, reveals that junk DNA may have an unanticipated use in providing the speed-ballast for animal growth.

The same ***Cellular Phylodynamic Analysis***, that is to say, the examination of growth and development in units of numbers of cells, ***N***, has made it possible to identify several additional features of the formation of the animal body, which will aid in our analysis of aging (APPENDIX and Reference 6). Such an examination of the growth of many tissues, organs, and anatomical structures, in units of numbers of cells, ***N***, revealed that parts of the body frequently arise and grow by an expression we call the ***Cellular Allometric Growth Equation***^6^:

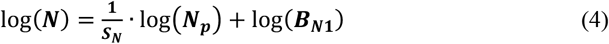

***where N***_***p***_ is the number of cells in a cell lineage, tissue, organ, or anatomical structure of the body (APPENDIX and Reference 6). We call the parameter ***B***_***N1***_ the ***Cellular Allometric Birth*** and ***S***_***N***_, the ***Cellular Allometric Slope***.

This ***Cellular Phylodynamic*** approach also makes it possible to reconstruct plausible ***cell-lineage-trees*** of animals and their parts, even in the absence of cell tracing microscopy, from the ***Universal Growth*** and ***Cellular Allometric Growth Equations*** and their parameters, by a method we call ***Cellular Population Tree Visualization Simulation*** (APPENDIX and Reference 6). Finally, the ***Cellular Phylodynamic*** examination of various parts of the body reveals that they often mold their final composition by making ***Clones Within Clones***, in such cases as the relative growth of the front and back parts of the drosophila wing, the creation and growth of the organs of nematodes, or the creation and maintenance of the crypts of the mammalian intestine (APPENDIX and Reference 6).

### Aging

The relationship between ***age, t***, and ***Mitotic Fraction, m***, can be calculated by combining Equations#1 and #3:

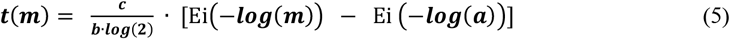

We call this expression, the ***Universal Mitotic Fraction in Time Equation***. With it, and the ***a, b***, and ***c*** values derived from the fit of growth data to the ***Universal Growth Equation*** (#3), we could generate curves for each of the 10 species listed above, ending at the age of the longest known individual in each species. In these curves, the ***Mitotic Fraction***, can be seen to mark mitosis in close to all of the cells of embryo soon after conception, while by the end of life, greater than 99.9% of the body’s cells are no longer engaged in mitosis (FIGURE 2). We call this boundary, marking where the ***Mitotic Fraction*** has declined from almost 100% at conception, to less than 0.1% at the end of life, the ***Death Zone***.

**FIGURE 2:**
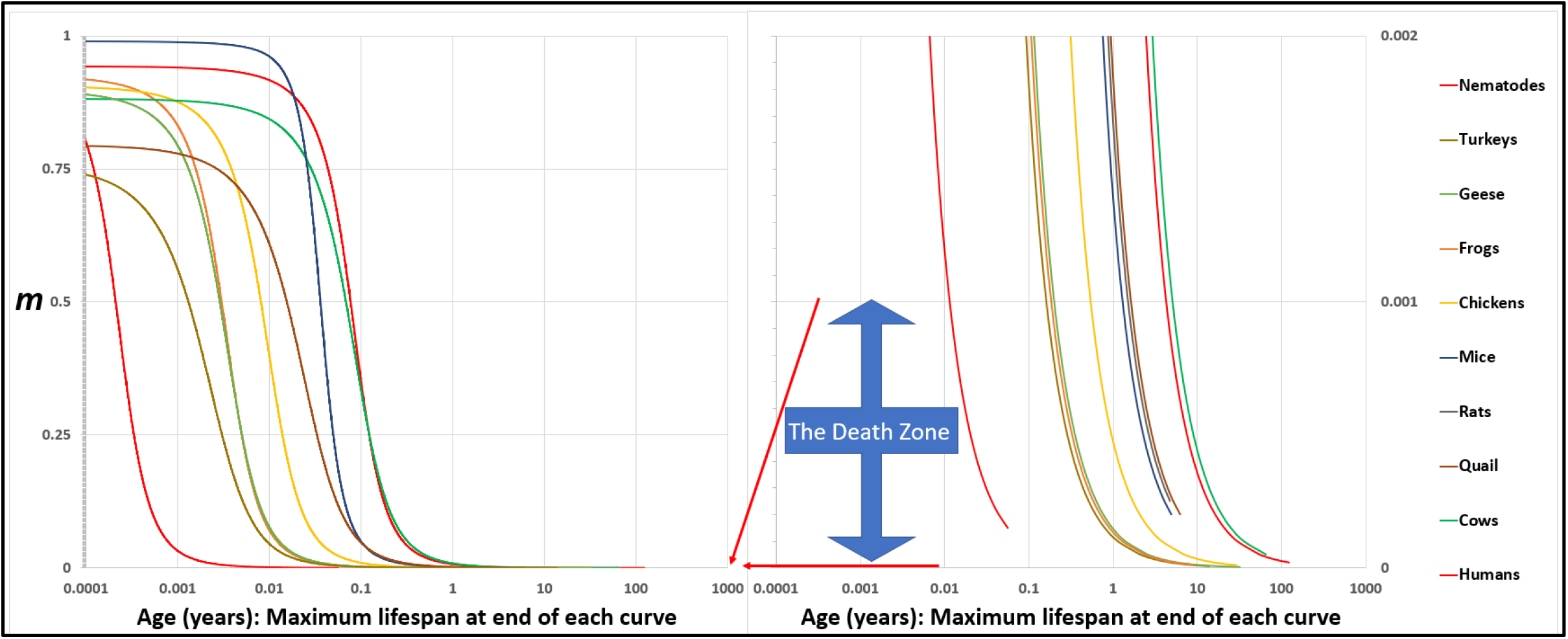
The *Universal Mitotic Fraction in Time Equation* (#5) captures change in *Mitotic Fraction, m*, with *Age, t*. 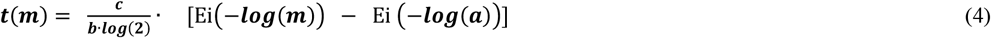

The graph on the right side shows the bottom 0.2% of the Y-axis of the graph on the left side: the ***Death Zone*** occurs when fewer the 0.1% of the cells of the body are dividing. Each curve ends at the age of the oldest known member of its species. Humans (♂) (Homo sapiens), Frogs (Rana pipiens), Nematodes (C. elegans), Chickens (Gallus gallus), Cows (Bos taurus), Geese (Anser anser), Mice (Mus musculus), Quail (Colinus virgianus), Rats (Rattus norvegicus), Turkeys (Meleagris gallopavo). See APPENDIX and Reference 6 for details.

### Lifespan

All three parameters of the ***Universal Growth Equation, a, b*** and ***c***, considered genetically, allow species to evolve to patterns of growth that give the greatest chances of survival, and to lifespans calculable with this expression:

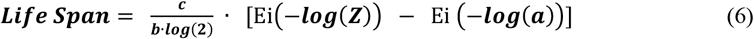

***Z*** is the ***Mitotic Fraction*** at the end of life. We call Equation#5 the ***Universal Lifespan Equation***, whose calculations, based on ***a, b***, and ***c*** values from growth data, distinguished between the longest lifespans (humans and cows), and the shortest (nematodes), lumping in between animals with 1-to-20-year lifespans (FIGURE 3). For the ten species we have examined, the average ***Z*** value of the average lifespan for each species, ***Z***_***A***_≈0.000037. The average ***Z*** value of the longest known lifespans, ***Z***_***L***_≈0.000024.

**FIGURE 3:**
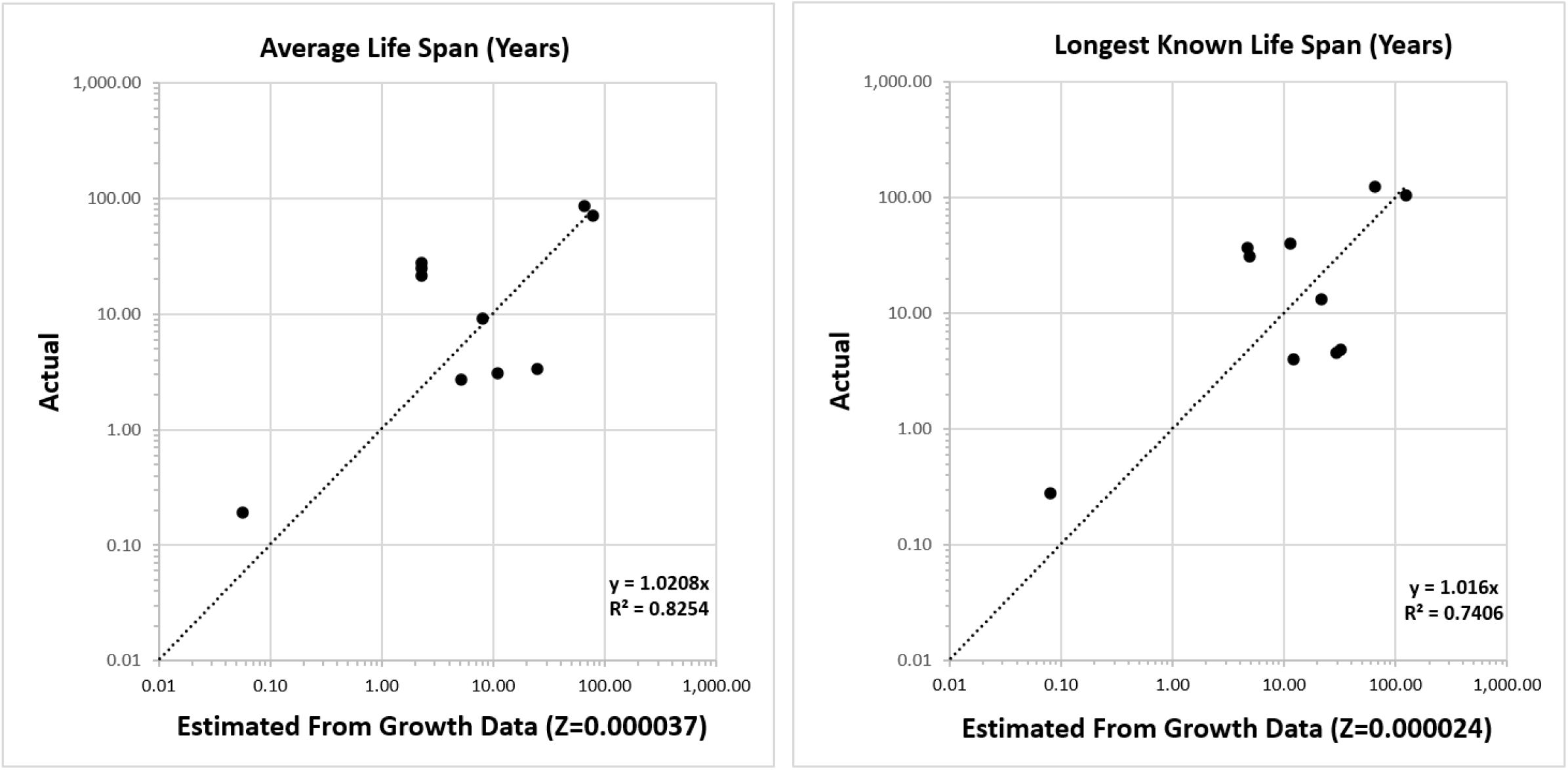
Life Span - Actual vs Predicted from Growth Data with the *Universal Lifespan Equation* (#6). Datapoints are for Humans, Frogs, Nematodes, Chickens, Cows, Geese, Mice, Quail, Rats, Turkeys. See APPENDIX and Reference 6.

## DISCUSSION

### Dilution or Destruction

Why would it be harmful for an animal’s ***Mitotic Fraction*** to dip to below one-in-a-thousand cells? Many irreversible chemical events, including protein denaturation, oxidation, and DNA mutation^23^, degrade cells, but most of this damage is flushed away by mitosis, when 50% or more of the cell’s components are made anew, and genome repair occurs.^24^ This concept of cell repair by dilution with new materials made at mitosis, thus driving “the cause of aging and control of lifespan”, was articulated by Gladyshev,^25^ expanding Sheldrake’s general suggestion,^26^ and Hirsch’s mathematical examination.^27^ As Ferrucci noted: “Aging is the ratio between damage accumulation and compensatory mechanisms. … we can replace these molecules … new DNA, new protein, new organelles, and so on … [thus] … aging starts at conception.”^28^ This process of aging decline, by infrequency of mitosis, leading to insufficiency of dilution repair, from conception till death, is captured quantitatively by the ***Universal Mitotic Fraction in Time*** and the ***Universal Lifespan Equations***.

### Why We Age

Some creatures, such as hydra, may avoid the ***Death Zone*** with a “constant and rapid cell turnover”^29^, with surplus cells pushed out into new buds of life. Other apparent immortals, such as mole rats, may grow so slowly that we lose interest in waiting for them to become old, while others never stop growing, i.e., marked indeterminate growth.^1^ However, most of us are fated by Darwinism to grow to sizes that maximize survival, achieved by ***Mitotic Fraction*** declining into the ***Death Zone***. The price of becoming the right size is decay, and ultimately, death.

### Aging is Reversible

The cellular damage that occurs when the ***Mitotic Fraction*** enters the ***Death Zone*** isn’t irreversible. Germ cells appear to rejuvenate by resetting the ***Mitotic Fraction*** to ∼100%, a region called ***Ground Zero***, adopting the terminology and insight of Gladyshev^30^, and the mathematics of the ***Universal Mitotic Fraction Equation*** (#1) when ***N≈*0**. A similar process may rejuvenate embryonic stem cells, and cells used to clone animals.^31^

### Growth and Aging Are Modifiable

Genetics, diet, pharmaceuticals, and other interventions have been found to change lifespan, often accompanied by altered growth, while large animals often live longer than small animals, and many genetic changes have been found to affect both growth and lifespan (reference 32, and others scattered throughout this report). These opportunities can now be examined quantitatively, with the ***Universal Growth*** and ***Universal Lifespan Equations*** offering both a mechanism, and a mathematics, for comprehending, and testing, aging’s link to growth.

### Varieties of Lethality

Death comes in great variety, from a myriad of places. Thus, the ***Mitotic Fraction*** in the whole body may be less important than in body parts such as stem cells^33^, heart,^34,35^ or immune system.^36,37^ Fortunately, the growth of such body parts, and thus the ***Mitotic Fraction*** in each of these places, is calculable with another expression of this ***Cellular Phylodynamic*** mathematics, the ***Cellular Allometric Growth Equation*** (#4, APPENDIX, and Reference 6).

### Aging and Health

For humans, ***Z*** provides a measure of the medical and public health system’s capacity to deal with vulnerability induced by aging’s reduction in ***Mitotic Fraction***. This frames useful questions: Do we search for actions and treatments that delay ***Mitotic Fraction’s*** approach to the ***Death Zone***, or do we look for ways to remain healthier after the ***Mitotic Fraction*** enters the ***Death Zone***, learning how to live longer with the burden which growing poses to aging?^38^

### Measuring Age and its Amelioration

Various laboratory measures of age have been developed,^39,40^ although the ultimate, and causal, metric might well be the ***Mitotic Fraction*** itself. By counting mitotic figures in histological sections, the ***Mitotic Fraction*** can be determined for single points in time, while measurements of the sizes of the clones in the liver marked by the activation of plasma protein genes, which occurs at a regular rate,^41^ can provide life-long values. Such measurements should improve the accuracy of the ***Universal Lifespan Equation***, for which the calculations described above have had to rely on ***a, b***, and ***c*** parameter values from growth studies, mostly early in life. These direct ***Mitotic Fraction*** measurements should also provide rapid screening and testing of pharmaceuticals for anti-aging activity.

### Anti-Aging Therapeutics: Life Extension

Many compounds have been found to extend life, in a variety of animals, most relevantly to us, in mice, a few of which are beginning to be tested in humans.^42,43,44^ So, why haven’t we found the pill that will make us live longer? A plausible difficulty has been the inability to predict, and test, the impact that a candidate agent might have on the extension in life, and the negative consequences associated with the pharmaceutical, so that the right agent, at the right dose, and the right schedule, could be identified and put to work to achieve a reasonable and predictable lifespan extension with minimal negative consequences. The ***Cellular Phylodynamic*** mathematics of growth and aging outlined here should help fill that gap.

For those agents found to change growth and aging together, identifying which parameter of the ***Universal Growth*** and ***Universal Lifespan Equations*** is changed, that is, ***a, b*** or ***c***, should provide actionable information, since each parameter can be expected to have different therapeutic effects, and different negative side effects. This can be measured in nematodes in days, and in mice in months, by fitting survival and growth data to the ***Universal Growth*** and ***Universal Lifespan Equations***. These equations can then be used to identify the optimal dose and schedule. They can also guide the search for new agents, and inform the synthesis of new analogues with the appropriate pharmacological qualities of affinity, half-life, and slow release.

Pharmaceuticals that are found to change growth by ***a*** or ***b*** may bump up the ***Mitotic Fraction***, and reverse aging, temporarily, returning the body to the vigor of a younger age, but also increasing body size, that is, making the body acquire more cells, which will probably be harmful; below we shall examine the possible utility of such agents in aiding vaccination, for COVID-19 especially. Pharmaceuticals that are found to slow growth by ***c*** will slow the decline in the ***Mitotic Fraction***, thereby stretching out ***Universal Lifespan Equation***, and slowing aging. They will not reverse the body to a younger state, but they will have the advantage of not having the potentially harmful negative effects of increasing body size. Agents such as rapamycin-related compounds might be expected to have this quality, a possibility that should be examined experimentally. Identifying the magnitude of the impact of such agents on the ***c*** parameter would make it possible to predict the dose/outcome relationship, by re-calculating the ***Universal Growth*** and ***Universal Lifespan Equations***. Such possibilities are obviously testable in mice.

### Anti-Aging Therapeutics:COVID & Immunity

Serious illness and loss of life in the current COVID-19 pandemic have occurred predominantly in the elderly, with the chance of COVID-19 death increasing exponentially with age.^45,46^ The efficiency of COVID-19 immunization has also been found to decrease with age, with too many excellent studies to list them all,^47,48,49^ in agreement with the long appreciated, if under-researched, dilemma of decreased efficacy of many vaccines in the elderly.^50^ This falls in line with the well-known decline in immune responsiveness that occurs with age.^36,37^

The data and math outlined here raise the possibility that aging of the immune system, as in the body as a whole, may be ascribable to the failure of mitotic dilution repair, traceable to the decline in the ***Mitotic Fraction*** into the ***Death Zone***. Such a consideration raises the possibility that vaccine effectiveness might be improved by increasing the ***Mitotic Fraction*** in the body, and in the immune system, at the time of immunization. This strategy would temporarily return the distribution of times since last mitosis for the cells of the body to the distribution of the times of an individual of a younger age, for whom the vaccine is known to be fully effective. Such a distribution of inter-mitotic times can be calculated with the ***Cellular Population Tree Visualization Simulation*** (APPENDIX and Reference 6).

Useful bumps in the ***Mitotic Fraction*** would be expected to occur for agents that increase the value of the ***a*** and ***b*** parameters of the ***Universal Growth*** and ***Universal Lifespan Equations***, but don’t change the value of ***c***. Whether this pans out can be tested in mice, as young mice are more easily immunized against COVID-19 than older mice, and this has been the basis for experimental studies of improving COVID-19 vaccination.^51^

A superficial look at the growth curves modified by growth hormone and other agents of the somatotropic pathways suggest that they both push up growth (i.e., ***a*** and ***b***) and stretch/contract growth (i.e., ***c***), but a serious look at which agents have which effects, in nematodes and mice, as described above, would replace speculation with numbers. Once promising agents are identified, their potential for achieving an improvement in vaccine effectiveness should be calculable with this ***Cellular Phylodynamic*** mathematics, and tested in mice, again as noted above.

Of course, agents that bump up the ***Mitotic Fraction*** in just the immune system, rather than in the body as a whole, would be ideal. Fortunately, the immune system, like any part of the body, is made of cells, which can be counted, and is thus amenable to ***Cellular Phylodynamic Analysis***. Since the discovery in the 1960’s that not all lymphocytes are the same, our understanding of the web of cellular activity in the immune system has increased greatly. However, until recently we have not been able to characterize numbers of immune cells, and their lineages, with precision, but this has recently be achieved by Plambeck et al.^52^ Note the ***cell-lineage-tree*** of a T cell clone in FIGURE 1I of that paper, which is of identical appearance to the ***cell-lineage-tree*** of a nematode, from which we extracted values, in units of numbers of cells, ***N***, which revealed that the whole body grows by the ***Universal Mitotic Fraction*** and ***Growth Equations***, while the clonal parts of the nematode body grow by the ***Cellular Allometric Growth Equation*** (#4, see APPENDIX and Reference 6). There is no reason why the clones of the immune system, and the populations which they comprise, could not also be mapped with data determined by the methodology of Plambeck et al^52^ to ***Cellular Phylodynamic*** equations, whose parameters could be traced to the ***Mitotic Fraction, m***, and ***Cell Cycle Time, c***, as well as measuring how the values of these parameters might change with age. Such an approach would allow us to understand the immune response, and its change with age, quantitatively, and design optimally effective strategies for improving immunization in the elderly, among other things. All that needs to be done is the hard work of counting the cells.

### Dilution, Mutation, Cancer

The features of cellular decay tell us that the mechanism of mitotic dilution isn’t a theory of aging, but a mechanism of all theories of aging, with the ***Universal Mitotic Fraction in Time Equation*** providing a measure for how fast all these processes go downhill. Mitotic dilution is also unlikely to be the only mechanism of aging. In a recent technical tour de force, Cagan and colleagues have found that the rate of somatic mutation, as measured in the intestine, scales with age.^23^ This effect, no doubt protective of cancer, is unlikely to be the result of the decline in the ***Mitotic Fraction***, as the cells in the intestine undergo a rate of mitosis vastly greater than the ***Mitotic Fraction*** in the body as a whole (although one point of view holds that the precursors of the crypt stem cells may have a more conventional level of mitotic activity^53,54^). As the authors concluded, “The most notable finding of this study is the inverse scaling of somatic mutation rates with lifespan. … Even if clear causal links between somatic mutations and ageing are established, ageing is likely to be multifactorial. Other forms of molecular damage involved in ageing could be expected to show similar anticorrelations with lifespan”. The authors also point to their own, superb, recent study^55^, which found that individuals with a genetic predisposition to DNA mutation display an increased risk of cancer, but not a pronounced reduction in life expectancy.

Cancer frequently occurs late in life, and thus has a surprisingly small contribution to life expectancy. Gandjou has calculated that even if we found the imaginary cure for cancer, life expectancy at birth would only increase by about 3.25 years.^56^ Of course, it’s obvious that animals with long lifespans would never reach their potential if they displayed rates of cancer incidence seen in animals with short lifespans, and Cagan and colleagues have provided a long-sought answer to this previously mysterious problem. The ***Cellular Phylodynamic*** mathematics outlined here should provide a starting point for sorting out the contributions of growth, and other factors such as mutation and cancer, to the overall survival.

### Life History, Lifespan, Evolution

Animals grow at different speeds, to different sizes, and different lifespans, and the ***Cellular Phylodynamic*** mathematics described above show us how this variety, and genetic variation leading to this variety, can be captured by the ***a, b***, and ***c*** parameters of the ***Universal Growth*** and ***Lifespan Equations***. These qualities go to the heart of evolution, and its molding by life history.^57^ For example, the time till maturity and reproduction is determined by the curviness of the “***S*”** in the ***S-shaped*** growth curve of the ***Universal Growth Equation*** (#3); the “***S*”** can stand tall, rapidly growing towards adulthood, or slouch languidly, slowly edging away from adolescence, all determined by the ***a, b***, and ***c*** parameters, together with lifespan, characterized by the ***Universal Lifespan Equation*** (#6).

The ***a, b***, and ***c*** parameters are also polymorphic pleiotropic genetic determinants, with positive impacts on growth early in life, and negative impacts on survival later, conforming to antagonistic pleiotropy theory of Williams^3^ and Hamilton^4^ and disposable soma theory of Kirkwood.^58^ A species may also delay entry into the ***Death Zone*** by steps such as enlisting apoptosis to boost the ***Mitotic Fraction*** or investing in enzymatic repair. This may account for some species-specific variability in the value of ***Z*** parameter below the one-in-a-thousand ***Death Zone*** border we have marked off. Medawar’s mutation accumulation theory^2^ provides a framework for comprehending the dilemma; is it worth the effort, as measured by Darwinian selection, to fight off the inevitable?

### The Pace and Shape of Lethality

While ***Z*** provides a single measure of lifespan, death occurs continuously, frequently increasing in magnitude, sometimes exponentially^1^, as noted by Gompertz in 1825^59^. The ***Universal Mitotic Fraction in Time Equation*** also displays an accelerating decline in ***Mitotic Fraction***, although of different form than Gompertz’s. Of course, mortality probably doesn’t directly reflect the decrease in the ***Mitotic Fraction***, but the increase in the times since last mitoses, which can be calculated with ***Cellular Population Tree Visualization Simulation*** (APPENDIX and Reference 6).

Species have lifespans that are short and long, whose risk of death may increase exponentially with age, as Gompertz showed for humans, or in other ways, or not at all, as Vaupel and colleagues made clear in the taxonomically wide survey of aging.^1^ This variety has been described as the pace and shape of lethality, appearing as the “two orthogonal axes of life history”.^1,^60 Note that for the 10 species we have examined, the curves of the ***Universal Mitotic Fraction in Time Equation*** (#5) cross over each other (FIGURE 2). Such interweaving is ascribable to the capacity of the parameters ***a*** and ***b*** to twist these curves and the capacity of the parameter ***c*** to stretch or contract them. This suggests a useful place to begin to consider the possible basis for the quantitative pathways of aging and their consequences.

### Growth and Aging

The ***Cellular Phylodynamic*** analysis of the data examined here suggests that animal growth and animal death are inescapably bound together, both arising from the same taming of mitosis. At the end of life, our now minuscule and plummeting ***Mitotic Fraction*** may no longer be perceptible in the mathematically negligible increase in size suggested by the ***Universal Growth Equation***, but it will be perceived by the increase in our frailty suggested by the ***Universal Lifespan Equation***. Thus, the size to which we grow, and the time at which we die, go hand in hand, both driven by the same reduction in the fraction of cells dividing. While the reward for this reduction in the ***Mitotic Fraction*** in space and time is the ability to grow to specific shapes and sizes^41^, the cost is the disability of cells starved of life-sustaining replacement materials made at mitosis. Our hope is that ***Cellular Phylodynamics*** might help us find ways to hold back that bitter hug of mortality.

Note: this study has not been peer reviewed and the findings could change.

