## APPENDIX for "On How, and Why, and When, We Grow Old"

### Analysis, Data, and Mathematics, in Full Detail


**James Michaelson PhD**

Department of Pathology, Massachusetts General Hospital
Department of Surgery, Massachusetts General Hospital, USA
Department of Pathology, Harvard Medical School, USA

15 DEER COVE ST UNIT # 2
LYNN, MA 01902-3119
United States
TEL (cell phone) 617 501 0590


#### TOC

***CELLULAR PHYLODYNAMICS***

The ***Cellular Phylodynamics*** of growth

***Cellular Phylodynamics*** is a mathematics of the discrete nature of cells and the cellular events that cause change in cells.

The ***Computational Animal***

Current concepts in growth and development

***UNI-GROWTH***

Data for the ***Cellular Phylodynamic Analysis*** of the growth of the whole body.

Basic features of growth and cell division: ***exponential growth, biotic potential***, the ***Mitotic Fraction***, and ***quiescence***

The ***Mitotic Fraction Method***, where the value of the ***Mitotic Fraction***, ***m***, is measured over the full range of growth from fertilization until maturity.

As animals grow in size, $N_w$, the ***Mitotic Fraction***, ***m***, declines rapidly.

As animals grow in size, $N_w$, the declines in the ***Mitotic Fraction***, ***m***, is well fit by the ***Universal Mitotic Fraction Equation***, $m = a^{(N_w^b)}$

The relationship between size, $N_w$, and age, ***t***, is well fit by the ***Universal Growth Equation***: $\int \left(\frac{\log(2)}{c}\right) N_w a^{(N_w^b)}$

Closed form solution to the integration of the ***Universal Growth Equation***

Density dependent growth equations

The relationship of the ***Mitotic Fraction*** (***m***) and size ($N_w$) to the ***logistic equation***.

The relationship of the ***Mitotic Fraction*** (***m***) and size ($N_w$) to the ***Gompertz equation***.

The relationship of the ***Mitotic Fraction*** (***m***) and size ($N_w$) to the ***West equation***.

Fit of the relationship of the ***Mitotic Fraction*** (***m***) and size ($N_w$), to the ***Gompertz***, ***logistic***, and ***West equations***.

Fit of the relationship between size ($N_w$) and Age (***t***) to the ***Gompertz***, ***logistic***, and ***West equations***.

Fine scale "***ripples***" identify a lower level of growth detail that occurs in the life of the organism.

The Meaning of ***UNI-GROWTH***

The ***Universal Growth Equation*** describes how growth occurs

The ***Universal Growth Equation*** is linked to the cell biology behind growth.

The ***Universal Growth Equation's*** description of how fast we grow is linked to the ***Cell Cycle Time***, ***c***, which is linked to genome size, which is linked to how much junk DNA we have.

The ***Universal Growth Equation's*** description of the fraction of cells dividing, the ***Mitotic Fraction***, ***m***, is linked to the action of cell signaling molecules that induce mitotic quiescence.

***UNI-GROWTH***: Summary Definition

***ALLO-GROWTH***

- ***The Cellular Phylodynamic Analysis*** of the growth and development of the parts of the body
  - The ***Allometric Growth Equation***
  - ***Cellular Allometric Growth Analysis***
    - Early development: ***Cell lineages***: ***Cellular Allometric Lineage Growth Analysis***:
    - Late development: ***Tissues***, ***Organs***, and ***Anatomical structures***: ***Cellular Allometric Batch Growth Analysis***:
- The Meaning of ***ALLO-GROWTH***, and its defining abstraction, the ***Cellular Allometric Growth Equation***
  - The ***Cellular Allometric Growth Equation***
  - The ***log-linearity*** of body part ***ALLO-GROWTH*** can be traced to whole body ***UNI-GROWTH***
  - Parameter $S_N$, the ***Cellular Allometric Slope*** of the ***Cellular Allometric Growth Equation***, tells us how the embryo uses the ***Cell Cycle Time*** to drive ***Cellular Selection*** to adjust the size of each part of the body:
    - Parameter $S_N$, the ***Cellular Allometric Slope***, at work in adjusting the size of each part of the body:
      - Parameter $S_N$, the ***Cellular Allometric Slope***, at work early in development
      - Parameter $S_N$, the ***Cellular Allometric Slope***, at work late in development
      - Parameter, $S_N$, the ***Cellular Allometric Slope***, $S_N$, and the ***Cell Cycle Time***, $c_p$,
      - The role of DNA methylation on the time and nature of change in the ***Cell Cycle Time***, $c_p$ within the body
  - Parameter, $B_{N1}$, the ***Cellular Allometric Birth*** of the ***Cellular Allometric Growth Equation***, tells us how embryos create body parts from single cells
    - Data on the number of ***Founder Cells*** that make various parts of the body.
    - When the parts of the body are born: The ***Cellular Allometric Birth in Time***, $B_T$
    - The time of the ***Cellular Allometric Birth*** of a body part affects its size.
    - Body parts can be made of one ***Founder Cell*** or more than one ***Founder Cell***.
  - The ***Cellular Phylodynamic Analysis*** of ***Drosophila Growth*** and ***Development***
    - Numbers of cells in the fly as a whole, $N_w$.
    - Numbers of cells in the body parts of the fly, $N_p$.
    - Log-Linearity of the increase in the numbers of cells in various body parts of the fly, $N_p$.
    - Mitotic Recombination
    - Gynandromorph Marking
    - Body Part Size to Body Part Size Relationships; ***Body Part Proportion Equations***
    - ***Clones Within Clones***
  - ***ALLO-GROWTH***: Summary Definition
  - The ***E Unim Pluribus*** Hypothesis: Perhaps Embryos Make Their Body Parts From Single ***Founder Cells***

***CELLULAR POPULATION TREE VISUALIZATION SIMULATION***

***AGING***

**PRESENT AND FUTURE OPPORTUNITIES FOR *CELLULAR PHYLODYNAMIC ANALYSIS***

4D Microscopy

Cell Death

Manual Execution of ***Cellular Allometric Lineage Growth Analysis***

Computational Execution of ***Cellular Allometric Lineage Growth Analysis***: The ***BinaryCellName*** Method

Comprehending Development in 4 Dimensions at the Microscopic Scale with the ***BinaryCellName*** Method

Comprehending Development in 4 Dimensions at the Macroscopic Scale with the $\boldsymbol{S_{N3}}$ Method

JAVA Code and Flow Charts for ***Cellular Population Tree Visualization Simulation***

Flow chart and code for ***Cellular Phylodynamic Lineage Simulation***

Flow chart and code for ***Cellular Phylodynamic Lineage Simulation*** of BODY PARTS

***CELLULAR PHYLODYNAMICS***

In the main text of this communication, we presented a condensed description of a new method, ***Cellular Phylodynamics***,[1] and its findings, focusing on aging, by which we mean the consideration of growth, development, and aging in terms of numbers of cells, $N$.[2] The equations derived from this approach for animals generally are summarized in **BOX 1.** Equations in the text are given numbers less than 100, while the additional equations introduced in this APPENDIX begin with 100. Related equations have a letter following the number (11b), followed by a number (11b1), etc.

In this APPENDIX we provide a more detailed and thorough report of this analysis. This includes derivations of all of the equations for the growth, development, and aging of animals and humans, and a more complete presentation of the growth data, which is exhaustive.

The ***Cellular Phylodynamics*** of growth

The ***Cellular Phylodynamics*** approach is based on counting the number of cells in the animal as a whole, $N_w$. and in its various anatomical parts, $N_p$. These values of $N_w$ and $N_p$ provide the raw material for mathematical analysis, from which equations could be discerned. Thus, the equations that emerge from a ***Cellular Phylodynamic Analysis*** of growth, development, and aging aren't mathematical inventions, describing imaginary humans and animals, but empirically based generalizations of actual growth and development, which occur in actual animals, including ourselves.

***Cellular Phylodynamics*** is a mathematics of the discrete nature of cells and the cellular events that cause change in cells.

We, and almost all animals, begin life as a single cell, which divides to become 2 cells, then 4 cells, then 8, growing to the trillion or so cells of a human[3] or the thousand or so cells of a nematode worm[20], comprised of ***cell lineages***, ***tissues***, ***organs***, and ***anatomical structures***, each of which grows to its recognizable size. How does this occur? At the microscopic scale, our cells are intrinsically discrete entities.[83] There is no such thing as 1/3rd of a cell or 1.33 cells. It follows that the number of cells in the body, or in any part of the body, can only change their integer numbers by binary events, such as 1 cell becoming 2 cells by mitosis, or 1 cell becoming 0 cells by cell death. We can, and will, avail ourselves of continuous mathematics when it can provide us with useful approximations, but the ***Cellular Phylodynamics*** of embryology is fundamentally discrete in nature.

The ***Computational Animal***

Since cells only come in integers, whose numbers only change by discrete events[83], multicellular animals remind us of computers, also comprised of integer units,[4] which have also been called "cells"[5], whose states also undergo discrete changes. As we shall see, ***Cellular Phylodynamics*** provides us with a way to understand ourselves as ***Computational Animals***, whose growth, development, and aging can be understood as the aggregate consequences of the many discrete changes that occur among our cells.

Current concepts in growth and development

In contrast to the discrete nature of animal life at the microscopic scale, from our macroscopic perspective the growth of the body and its parts appears in continuous qualities of length, volume, mass, and shape. These continuous features of growth have yielded to quantitative analysis, although not always in ways whose meanings are clear. For example, it has long been appreciated that ***as we grow older***, ***we grow bigger***, and ***we grow slower***, ***until growth becomes imperceptible***, and a number of ***density-dependent*** growth equations have been developed (the ***logistic***, ***Gompertz***, ***von Bertalanffy***, ***Richards***, ***West***, and other equations: see below), which capture this increase in size and decline in the speed, and form classic ***S-shaped growth curves***.[6-11] Unfortunately, none of these ***density-dependent*** growth equations accurately fit the growth of real animals.[12-14] It has also long been appreciated that as we grow, the sizes of the ***tissues***, ***organs***, and ***anatomical structures*** of the body, when compared with the size of the body as a whole, often form straight lines on log-log graphs, a phenomenon known as ***allometric growth***.[15-19] However, the reason behind the striking log-linearity of ***allometric growth*** has long been a mystery. Finally, many studies have used 4-dimensional microscopy to characterize the growth of ***cell lineages*** that arise from single ***Founder Cells*** at the beginning of development,[20,21] but the precise way in which the embryo uses the control of cell division to create these first body structures has been obscure.[22-24] ***Cellular-Selection***, that is, differential cellular proliferation and death, creates many anatomical structures by outgrowth from the undifferentiated body mass, and molds much of the fine scale detail of anatomy, although how cell division is harnessed to achieve this is obscure.[98,99] Cell lineages grow from individual ***Founder Cells***,[20] although how embryos use mitosis to create these parts, and how these ***Founder Cells*** undergo ***Cell-Heritable*** change that characterized each clone, remains unknown. The analysis of cellular diversification, by single cell mRNA expression,[25,150,151] cell marking,[26,150,151,150,151] and high resolution 4D microscopy,[22,23,102] has also wrestled with the problem of cell lineage formation. As we have seen in the main text of this communication briefly, and will present in greater detail below, when we re-examine these features of growth, development, and aging in units of numbers of cells, that is, by ***Cellular Phylodynamics***, the biological basis for many of these processes becomes clear.

***UNI-GROWTH***

Data for the ***Cellular Phylodynamic Analysis*** of the growth of the whole body.

To carry out the ***Cellular Phylodynamic Analysis*** of growth, we assembled data on the number of cells in the whole animal, $N_w$, by age, $t$, in days from fertilization until maturity, for 13 species of animals: zebrafish (***Danio rerio***),[27] European sea bass (***Dicentrarchus labrax***),[28,29] mice (***Mus musculus***),[30,31] rats (***Rattus norvegicus***),[32] cows (***Bos taurus***),[33,34] humans (***Homo sapiens***),[35,36,37] bobwhite quail (***Colinus virgianus***),[38,39] domestic chickens (***Gallus gallus domesticus***),[40,41] turkeys (***Meleagris gallopavo***),[42] geese (***Anser anser***),[38] nematode worms (***C. elegans***),[20] frogs (***Rana pipiens***),[43] and clams (***Merceneria mercenaria***).[44,45,46] Data on the numbers of cells in early embryos, from fertilization onward, were available for mice,[47,48] rats,[49] cows,[50] humans,[51] chickens,[52] turkeys,[53] nematode worms,[20] frogs,[54] clams,[45] and fish.[55,56] Data on the numbers of cells in later embryos were assembled from published values in units of weight or volume by taking advantage of the finding that there are about $10^8$ cells/gram-cc.[57] Excel files (TABLE A1) characterizing these basic growth data ("**Basic Data Files**"), and the calculations made from these data ("**Calculations Files**"), are available on request. Also available to interested readers are excel files with the data on the growth of the parts of the body parts that we examined (see below).

The graphs of these various datapoints for animal size's ($N_w$) versus time ($t$), are shown in FIGURES A1-A3. Graphing the number of cells in the body as a whole, $N_w$, vs. age, $t$, from fertilization until maturity, reveals, for each of these 13 species, that animal growth occurs by ***S-shaped*** growth curves (Fig A1). This is most easily viewed on log-log plots, because the greatest part of growth occurs rapidly at the beginning of life. For example, in terms of numbers of cells, $N_w$, humans grow about 11 orders of magnitude in the womb and about 2 orders of magnitude after birth. As we shall see below, the reason why all of these creatures have similar ***S-shaped*** growth curves is that they all grow by an equation of the same form, the ***Universal Growth Equation***.

Basic features of growth and cell division: ***exponential growth, biotic potential***, the ***Mitotic Fraction***, and ***quiescence***

Because cells increase in number by cell division, they are capable of exponential growth, in which the size of an organism, $N_w$ (in terms of cell number), with age, $t$, can be thought of in terms of its ***rate of growth***:

$$\frac{dN_w}{dt} = \frac{ln(2)}{c} \cdot N_w \qquad (100)$$

where $c$ is the average ***Cell Cycle Time*** among those cells that are dividing, which, for exponential growth, is all cells.

Integration of Equation #100, translates the integer nature of cells into the continuous appearance of growth:

$$N_w = e^{\left(\frac{ln(2)}{c}\right) \cdot t} \qquad (100b)$$

Thus, data on the relationship between cell number, $N_w$, and time, $t$, at the very beginning of development, when all cells are dividing, can give one a practical measure of the animal's ***Cell Cycle Time***, $c$.

Of course, exponential growth (Equation #100b) is ultimately unsustainable, a fact captured by the concept of the ***biotic potential***.[58] The departure from the ***biotic potential*** of our cells is a fundamental feature of animal growth, which can be captured by introducing the term $m$, the ***Mitotic Fraction***, giving us this expression for the ***rate of growth***:

$$\frac{dN_w}{dt} = \frac{ln(2)}{c} \cdot N_w \cdot m(N_w) \qquad (100c)$$

Equation #100c embraces all forms of growth, but without specificity, as the manner by which the ***Mitotic Fraction***, $m(N_w)$, changes with the animal size, $m(N_w)$, is completely unspecified. As we shall see below, this lack of specificity gives the Equation #100c the power to accomplish such things as leading to a way to calculate how the ***Mitotic Fraction***, $m$, changes as we grow (see the ***Mitotic Fraction Method*** below). Additionally, insertion of the ***Mitotic Fraction***, $m(N_w)$, into the exponential growth equation (#100b), converts it into Equation #100c, and gives us the general form of a ***density dependent growth*** equation, which balances the unlimited quality of exponential growth ($\frac{ln(2)}{c} \cdot N_w$) with the inevitable limit of resources ($m(N_w)$).

Although our treatment of the ***Mitotic Fraction***, $m$, relies on an obsessively rigorous adherence to this mathematical definition in Equation #100c, from time to time we will revert, for the purposes of clarity, to speaking of $m$ as a rough measure of the ***growth fraction***, $G$, the fraction of cells dividing, which is usually measured by the cytogenetic analysis of the uptake of DNA precursors.[59] The two abstractions of ***Mitotic Fraction***, $m$, and ***growth fraction***, $G$, are obviously closely, but not precisely, related, since other factors (cell death, cell size, cell density) play roles in growth that are likely to be secondary to cell division. Definitionally, the ***Mitotic Fraction*** is a measure of the amount of actual growth that occurs as a fraction of the growth that would have occurred had each cell divided, ignoring other effects, such as cell death and cell size, a plausible approximation, as mitosis is growth's principal driving force. Thus, the ***Mitotic Fraction*** $m$ can be thought of as the main component of the ***growth fraction***, $G$, the precise fraction of cells that are dividing. Operationally, this imprecision has little practical impact, as it will be cancelled out when we reintroduce $m$ into new equations of growth that we shall derive below.

When we think imprecisely of the ***Mitotic Fraction***, $m$, as the fraction of cells dividing:

$$m \approx N_{Wd}/(N_{Wd} + N_{Wq}) \qquad (100d)$$

where $N_{Wd}$ is the number of cells dividing and $N_{Wq}$ is the number of cells that have ceased from dividing, a process known as ***mitotic quiescence***. ***Quiescence*** is the complex biochemical mechanism that occurs within cells to prevent them from progressing to mitosis, which is set in motion by inhibitory and stimulators growth factor molecules, hormones, and other signals that cells use to communicate.[60-62]

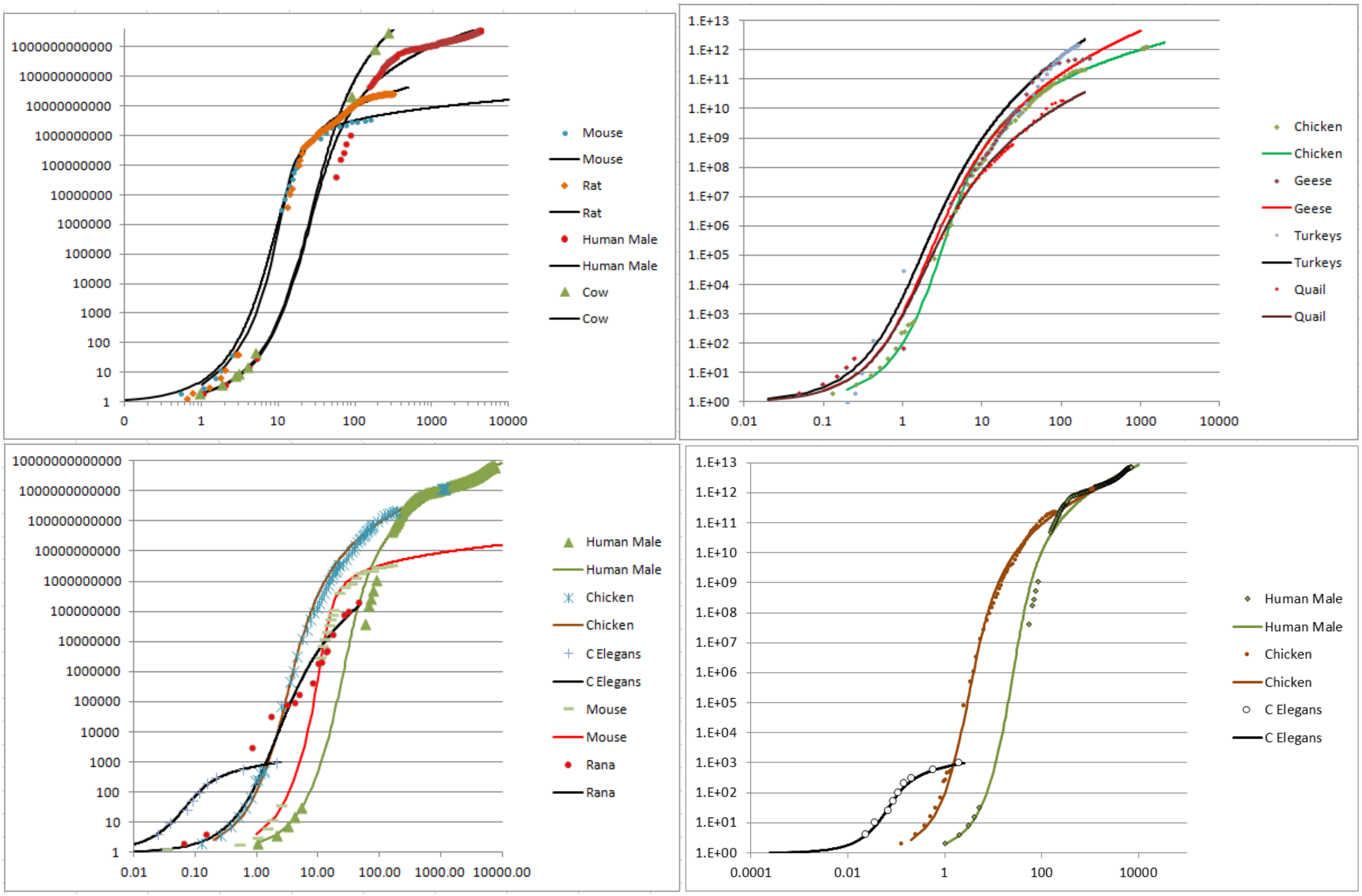


**FIGURE A1**: **Growth, in units of numbers of cells, $N_w$, from fertilization until maturity, in units of days, $t$.** Datapoints for animal size's ($N_x$) versus time ($t$), for ***C elegans***, chickens, mice, turkeys, quail, geese, frogs, and humans, and their fit to the numerically integrated form of the ***Universal Growth Equation*** (#5b)**.**

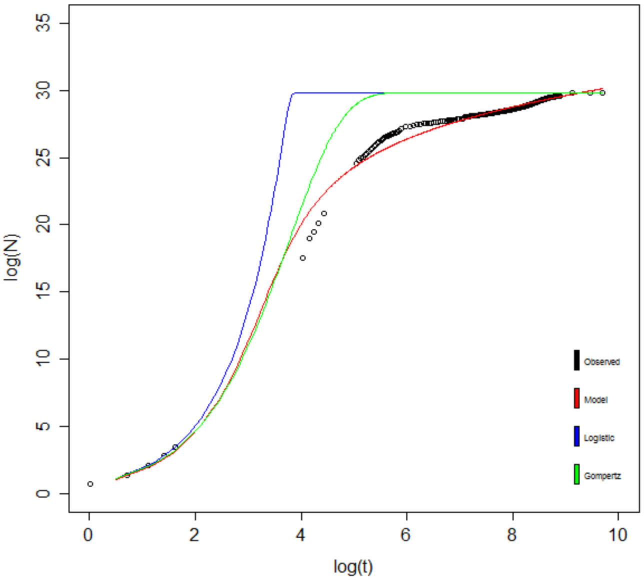


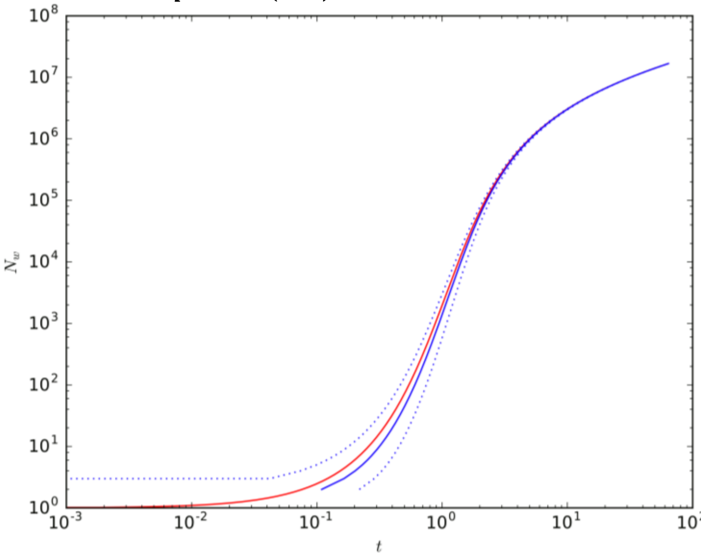


**FIGURE A2**: **Fit of human growth to the *Universal Growth Equation* (red), the *logistic equation* (blue), and the *Gompertz equation* (green).** Data values as black circles.

**FIGURE A3:**
**Analysis of the closed form integration of the *Universal Growth Equation* (#3).**

The ***Mitotic Fraction Method***, where the value of the ***Mitotic Fraction***, ***m***, is measured over the full range of growth from fertilization until maturity.

The value the ***Mitotic Fraction***, ***m***, for animals of various sizes, $N_w$, can be calculated by reversing ***Equation*** #100c:

$$m = \frac{dN_w}{dt} \,/\, [\frac{ln(2)}{c} \cdot N_w] \tag{3b}$$

We call the practical implementation of this equation the "***Mitotic Fraction Method***". To execute these ***Mitotic Fraction Method*** calculations, we use our assembled growth data to calculate the value of the ***Mitotic Fraction***, ***m***, (column M of the excel files listed in "**Calculations Files**" column of Table 1) by:

$$m_i = \frac{2N'(t_i)}{r(N_i + N_{i+1})} \tag{101}$$

The same iteration was also carried out based on an exponential calculation, rather than on the linear calculation show above, yielding essentially the same outcome (not shown).

Equation #101 occasionally resulted in a small number of data points with $m > 1$. Since this is both theoretically and numerically impossible (the biological meaning would be one cell dividing to give rise to more than two cells, which is nonsensical), these values of ***m*** were manually overwritten with a value of $m = .999$ noted by orange highlights in the excel spreadsheets.

As animals grow in size, $N_w$, the ***Mitotic Fraction***, ***m***, declines rapidly.

Our ***Mitotic Fraction Method*** calculations, based on the animal growth data noted above (Table 1), characterized how the value of the ***Mitotic Fraction***, ***m***, declines as growth occurs, that is, as $N_w$ increases, from fertilization until maturity, for nematodes, chickens, cows, geese, quail, turkeys, fish, clams, mice, rats, humans, and many other animals (FIGURES A4-A6). Graphs of the values of the ***Mitotic Fraction***, ***m***, calculated by the ***Mitotic Fraction Method***, for each of the diverse species of animals noted above, reveal that growth is close to exponential during the first few cell divisions ($m \approx 1$), but soon begins its "slippery slope" descent, resulting in a rapid and accelerating decline in the value of ***m***, reaching values of $10^{-2}$ to $10^{-6}$ by the time adult size is reached (FIGURES A4-A6).

As animals grow in size, $N_w$, the declines in the ***Mitotic Fraction***, ***m***, is well fit by the ***Universal Mitotic Fraction Equation***, $m = a^{(N_w^b)}$

The basis for the relationship between the ***Mitotic Fraction***, ***m***, and $N_w$ becomes evident by graphing the $\log(N_w)$ vs$-\log(-\log(m))$, revealing roughly straight rows of dots (FIGURE A4d). This suggests that the relationship between the ***Mitotic Fraction***, ***m***, and $N_w$ can be captured with the expression:

$$m = a^{(N_w^b)} \tag{1}$$

The best fit values for ***a*** and ***b*** for each species were calculated as regression inputs of $\log N_i$ and $-\ln(-\ln g_i)$ using Excel's LINEST function. Values for ***a*** and ***b*** were calculated from the LINEST output (in columns P-Q of the excel files listed in Table 1) together with the $r^2$ value. This relationship is well borne out, as indicated by the high $r^2$ values from non-linear regressions for each of the eleven animals listed in the Table 2.

Because Equation #1 so closely captures the relationship between the ***Mitotic Fraction***, ***m***, and size, $N_w$, from fertilization until maturity, for all of the animals we have examined, we call it the "***Universal Mitotic Fraction Equation***". ("Universal"; from the Latin "universalis"; Etymology: "uni (Latin, "one"), Versum (Latin, "turned"); thus something "turned into one"). Note that the ***Universal Mitotic Fraction Equation***, and the ***Universal Growth Equation***, that is derived from it, and which we shall describe below, holds for all of the animals that we have examined, and for all of the sizes we have examined, from the first fertilized cell onward.

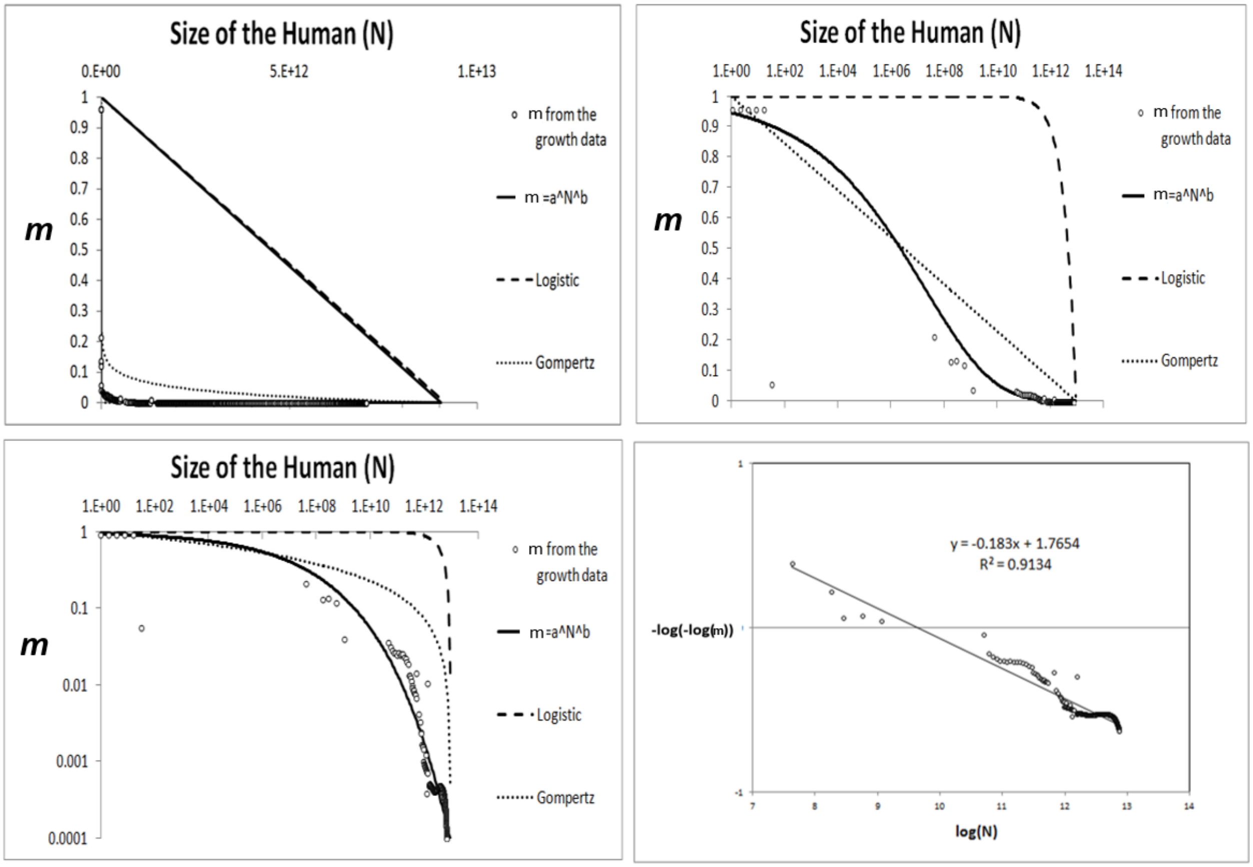


**FIGURE A4**: **Decline in fraction of cells dividing, the *Mitotic Fraction*, *m*, calculated by the *Mitotic Fraction Method*, that occurs as animals increase in size, as seen by the *Universal Mitotic Fraction Equation*.** Data points for size, ***N***$_w$, in integer units of numbers of cells, vs. the ***Mitotic Fraction***, ***m***, from fertilization, until maturity, for humans.

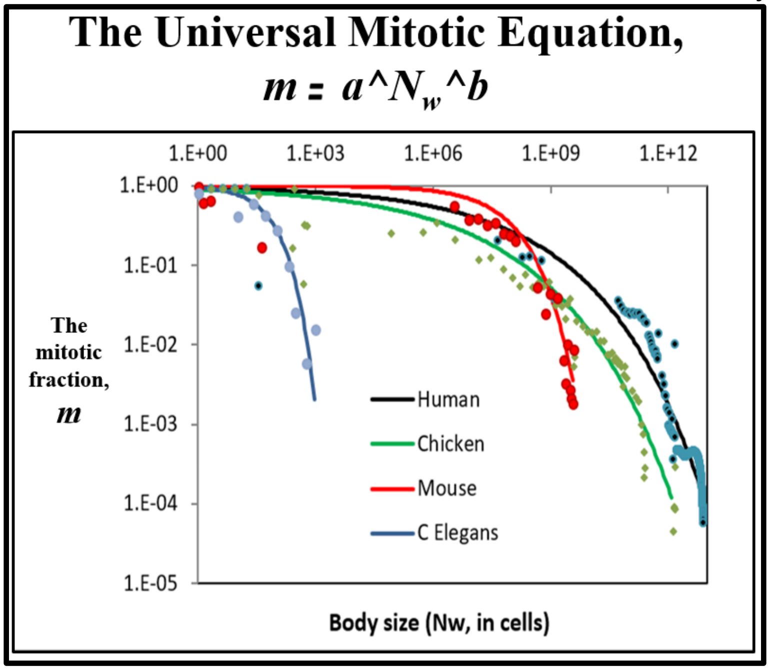


**FIGURE A5**: **Values of the *Mitotic Fraction*, *m*, calculated by the *Mitotic Fraction Method*, as a function of animal size, *N*$_w$, as shown at various scales.** Data points for animal size, ***N***$_w$, in integer units of numbers of cells, vs. the ***Mitotic Fraction***, ***m***, from fertilization, until maturity, for humans, chickens, ***C elegans*** nematode worms, and mice.

The relationship between size, $\boldsymbol{N_w}$, and age, ***t***, is well fit by the ***Universal Growth Equation***: $\int \left(\frac{\log(2)}{c}\right) \boldsymbol{N_w a^{(N_w^b)}}$

Combining Equation #3 and #4 leads to this expression for describing the rate of animal growth:

$$\frac{dN_w}{dt} = \frac{ln(2)}{c} \cdot N_w \cdot a^{(N_w^b)} \tag{5a}$$

The relationship between size, $\boldsymbol{N_w}$, and age, ***t***, is captured by numerical integration of Equation #5, where the age, ***t***, at fertilization is ∼0, and the age at death is ***d***:

$$N_w = \int_{t=0}^{t=d} \frac{ln(2)}{c} \cdot N_w \cdot a^{(N_w^b)} \tag{5b}$$

Numerical integration of Equation #5, to derive Equation #5b, was carried out $4^{th}$ order Runge-Kutta method, employed manually in Excel to carry out such growth curve reconstructions, so as to examine relationship between Size ($\boldsymbol{N_w}$) and Age (***t***). The step size ***h*** (cell b11 of the excel files listed in "**Calculations Files**" column of the excel files enumerated in Table 1) was set such that 10,000 steps was used for each organism. At each $\boldsymbol{t_i}$ (column W), the model value of $\boldsymbol{N_i}$ was calculated recursively in column X as

$$N_i = N_{i-1} + \frac{h(k_1+2k_2+2k_3+k_4)}{6} \tag{200}$$

where:

$$\text{if: } f(N) = N' = rNg \tag{201}$$

$$k_1 = f(N_i) \tag{202}$$

$$k_2 = f\left(N_{i-1} + \frac{k_1}{2}h\right) \tag{203}$$

$$k_3 = f\left(N_{i-1} + \frac{k_2}{2}h\right) \tag{204}$$

$$k_4 = f(N_{i-1} + k_3 h) \tag{205}$$

The values of each $\boldsymbol{k_i}$ are calculated in columns Z-AC of the excel files listed in "**Calculations Files**" column of the table above. Excel's VLOOKUP and CORREL functions were then used to calculate correlations of observed and model **ln *g*** and **ln *N*** (columns AM-AU). These correlations, as well as the regression correlation coefficient, were used as quality-of-fit metrics for the model and recorded in the summary columns A-B in the excel files.

The fit of Equation #5b, the numerically integrated form of Equation #5, to the various datapoints for animal size's ($\boldsymbol{N_w}$) versus time (***t***), are shown in FIGURE A1. The strength of this relationship was confirmed by the high $\boldsymbol{r^2}$ values for each of the eleven animals listed in the Table 2: humans, frogs, nematodes, chickens, cows, geese, mice, quail, rats, turkeys, fish, and clams. Because Equation #5, in its numerically integrated form (Equation (#5b)) closely captures the relationship between size, $\boldsymbol{N_w}$, and age, ***t***, from fertilization until maturity, for all of the animals we have examined, we call it the "***Universal Growth Equation***". We call the biological process which the ***Universal Mitotic Fraction*** and ***Universal Growth Equations*** capture "***UNI-GROWTH***".

Closed form solution to the integration of the ***Universal Growth Equation***

A closed form solution to the integration of the ***Universal Growth Equation*** (#5), can also be seen by considering:

$$\frac{dN}{dt} = r\,N\,a^{(N^b)} \tag{301}$$

where $r = \frac{\log(2)}{c}$. Rearranging, one obtains:

$$N^{-1}\,a^{(-N^b)}\,dN = r\,dt \tag{302}$$

which can be integrated as:

$$\int_1^{N_w} N^{-1}\,a^{(-N^b)}\,dN = \int_0^t r\,d\tau \tag{303}$$

Leading to the growth equation:

$$t = \left[\mathrm{Ei}\left(-N_w^b \log(a)\right) - \mathrm{Ei}(-\log(a))\right]\frac{c}{b\,\log(2)} \tag{3}$$

where $t = 0$ when $N_w = 0$, and **Ei** is the exponential integral function[63].

Equation #3 can be considered as a continuous approximation of the expected time required by a Poisson model with individual cell growth $r a^{(N^b)}$.

We show this by considering the random variable $t_N$ representing the time required to reach the number of cells $N$, starting from a single cell. This variable has expected value:

$$E[t_N] = \sum_{n=2}^{N} E[t_n - t_{n-1}] \tag{305}$$

We assume that each one of the $n$ cells that make up the organism at a given time behaves according to a Poisson process with rate $ra^{(n^b)}$ and that they are all independent. As a result, the event represented by the growth of a new cell out of any of the $n$ current cells behaves according to a Poisson process with rate $n\,r\,a^{(n^b)}$ (this can be easily seen by considering the cumulative distribution function of the exponential distribution). Therefore:

$$\overline{t_N} = E[t_N] = \sum_{n=2}^{N} \frac{1}{nra^{(n^b)}} \tag{306}$$

$$\sigma_N = \sqrt{\mathrm{var}(t_N)} = \sum_{n=2}^{N} \frac{1}{\left(nra^{(n^b)}\right)^2} \tag{307}$$

where $\overline{t_N}$ and $\sigma_N$ are, respectively, the mean and standard deviation of the time required to reach the number of cells $N$. The continuous version in #6 and the discrete (point-process) version in #306 provide very similar results. For example, using $c = 0.070$, $a = 0.925$, $b = 0.269$, we get the behavior shown in FIGURE A3 for the continuous version in red, and the discrete version in blue (mean ± std) (FIGURE A3).

Density dependent growth equations

A number of ***density-dependent*** growth equations have been developed to capture either the growth of the number of organisms in a population,[64,65] or the growth of the size of individual organisms,[7-11] none of which accurately fit the growth of real animals.[12-14] The simplest of these ***density-dependent*** growth equations, the logistic, occurs by $\boldsymbol{m}$ declining linearly as the size of the organism, $\boldsymbol{N_w}$, increases.[7] In the ***Gompertz equation***[8], $\boldsymbol{m}$ declines as the log of $\boldsymbol{N_w}$. Similarly, for many other equations, including the ***von Bertalanffy***[9], the ***Richards***[10], and ***West***[11] equations, the value of $\boldsymbol{m}$ declines monotonically as size, $\boldsymbol{N_w}$, increases.

The relationship of the ***Mitotic Fraction*** (***m***) and size (***N_w***) to the ***logistic equation***.

***Density dependent*** growth is captured by the ***logistic equation***[7] in the form:

$$\boldsymbol{m = 1 - \frac{N_w}{K}} \tag{401}$$

The relationship of the ***Mitotic Fraction*** (***m***) and size (***N_w***) to the ***Gompertz equation***.

***Density dependent*** growth is captured by the ***Gompertz equation***,[8] in the form:

$$\boldsymbol{m = \left(\frac{1}{\log K}\right) \log\left(\frac{N_w}{K}\right)} \tag{402}$$

The relationship of the ***Mitotic Fraction*** (***m***) and size (***N_w***) to the ***West equation***.

***Density dependent*** growth is captured by the ***West equation***,[11] for total mass of the organism (***M***), in the form

$$\boldsymbol{\frac{dM}{dt} = aM^{\frac{3}{4}}\left(1 - \left(\frac{M}{K}\right)^{\frac{3}{4}}\right)} \tag{403}$$

Units of mass are unspecified, so we can treat Equation #403 such that the units of mass (***M***) of our organism are in units of "the weight of a cell" (***N_w***). West's equation then becomes:

$$\boldsymbol{\frac{dN_w}{dt} = aN^{\frac{3}{4}}\left(1 - \left(\frac{N_w}{k}\right)^{\frac{3}{4}}\right)} \tag{404}$$

By re-arranging:

$$\boldsymbol{\frac{dN_w}{dt} = \left[\left(\frac{\log 2}{c}\right) N\right]\left[\left(\frac{ac}{\log 2}\right) N^{-\frac{3}{4}}\right]\left[1 - \left(\frac{N_w}{k}\right)^{\frac{3}{4}}\right]} \tag{405}$$

Thus, for any organism growing by the ***West equation***, the fraction of cells dividing, ***m***:

$$m = \left[\left(\frac{ac}{\log 2}\right) N_W^{-\frac{3}{4}}\right]\left[1 - \left(\frac{N_W}{k}\right)^{\frac{3}{4}}\right] \tag{406}$$

Equation #406 can be thought of as the ***West equation***, phrased in terms of the fraction of cells dividing, ***m***.

Fit of the relationship of the ***Mitotic Fraction*** (***m***) and size (***N***$_w$), to the ***Gompertz***, ***logistic***, and ***West equations***.

As can be seen in FIGURE A4, over the full range from conception to adult size, neither the Gompertz (Equation #402), nor the logistic (Equation #401) provides a close characterization of the behavior of the ***Mitotic Fraction***, ***m***, with respect to $\boldsymbol{N_w}$. As can be seen in FIGURE A6, while the ***West equation*** can be used to quite accurately capture growth over its full expanse, to do so requires $\boldsymbol{m > 1}$ early in development, which is mathematically valid, but biologically impossible, since individual cells would have to give rise to more than two daughter cells in a single ***Cell Cycle Time***. However, both our ***Universal Growth Equation*** (Equation #5), and the ***West equation*** (#406), work quite well (FIGURE A6) later in development. This means that our ***Universal Growth Equation*** is capable of achieving growth within the limits of the blood supply's capacity to support growth, as captured by the ***West equation***.[11]

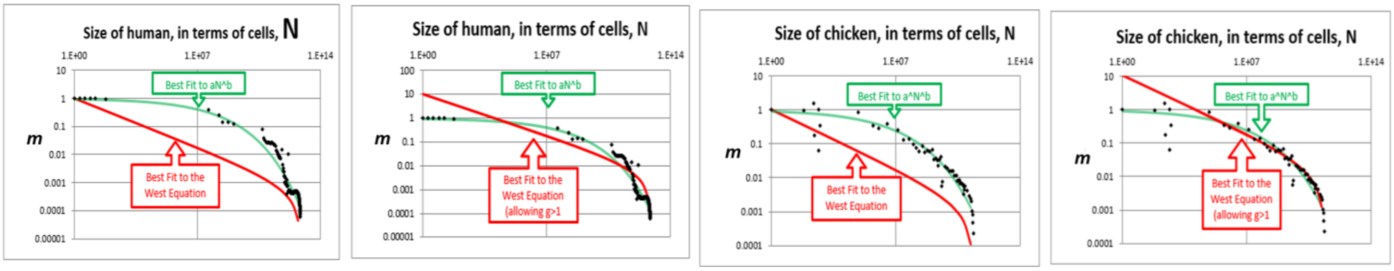


**<u>FIGURE A6</u>: Comparison of the fit of *Mitotic Fraction*, *m*, as a function of size, *N*$_w$, with human growth data, to the *Universal Mitotic Fraction Equation* (green) and *West equation* (red).**

Fit of the relationship between size ($N_w$) and Age ($t$) to the ***Gompertz***, ***logistic***, and ***West equations***.

We tested our data on body size, $N_w$, and age, $t$, against three of the most widely used density dependent growth equations (the ***Gompertz***, the ***logistic***, and the ***West*** [a variation of the ***von Bertalanffy***]) (FIGURE A2). None of these growth equations fit the actual growth data as well as the ***Universal Growth Equation*** (FIGURE A2).

Fine scale "***ripples***" identify a lower level of growth detail that occurs in the life of the organism.

Of course, no equation, however closely it may summarize actual data, can ever capture the world precisely. In fact, while the values for body size, $N_w$, and age, $t$, that we have assembled, fall remarkably close to the ***Universal Growth Equation***, we can also see hints of fine-scale minor "***ripples***" departing from the smooth edge of the curve (FIGURE A1 and A2). We suspect that these ***ripples*** identify specific events in the life of the organism, such as gastrulation, somite formation, weaning, puberty, and individual differences, that are associated with sex, malnutrition, genetic variation, and so on. To extract this second layer of growth information, we derived an expression, Equation #500, to measure the magnitude and timing of these growth residuals, $\boldsymbol{\delta_j}$:

$$m = A^{N^B} + \sum\delta_j \quad (500)$$

where each $\delta_j$ models a wave-like deviation away from the trend (the ***Universal Growth Equation*** #5b) that corresponds to an interpretable life-event in the growth of the organism. The appropriate functional forms for the $\delta_j$ may be determined through regression of the residuals of observed ***m*** against the ***Universal Growth Equation*** (#5b).

Indeed, a graph of these $\delta_j$ values, as a function of the size of the developing human male, $N_w$, which is shown in FIGURE A7 bears out our suspicion that these ***ripples*** identify biologically relevant processes, such as the right-most peak (at $\log N_w \approx 29.3$), which corresponds to puberty.

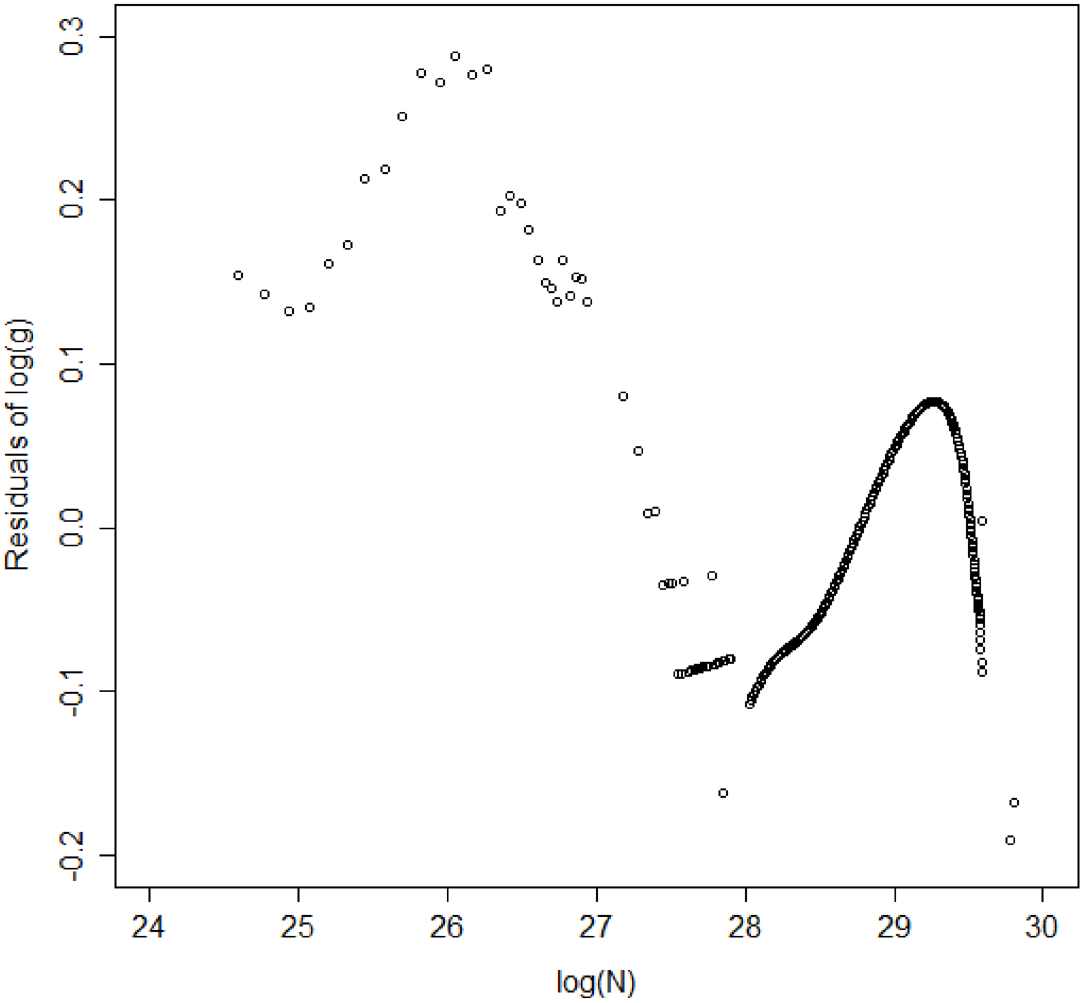


**FIGURE A7**: **Wave-like deviations ("*ripples*") from the *Universal Growth Equation curve*, for human growth.**

The Meaning of ***UNI-GROWTH***

The growth data shown in Tables 1 and 2, and FIGURE A1-A6 make clear that animals as diverse as mollusks, nematodes, fish, amphibians, birds, and mammals, including humans, all grow by the ***Universal Growth Equation*** (#5). The values of ***a***, ***b***, ***c***, and ***R*** make the ***Universal Growth Equation*** (#5) specific for each individual. Let us unpack the biological meanings of the form of the ***Universal Growth Equation*** and its parameters.

The ***Universal Growth Equation*** describes how growth occurs

Mathematically, each of the parameters of the ***Universal Growth Equation*** (#5) has a specific impact on the shape of the "***S-shaped***" growth curve. The ***a*** and ***b*** parameters determine the curviness of the "***S***". The ***c*** parameter, the ***Cell Cycle Time***, determines the speed of growth, expanding or contracting the "***S***" like an accordion.

The ***Universal Growth Equation*** is linked to the cell biology behind growth.

Biologically, each of the parameters of the ***Universal Growth Equation*** (#5) is linked to a specific aspect of cell division. The ***Mitotic Fraction***, ***m***, reflects how many cells are engaged in mitosis; ***m*** itself, which decreases as we become larger, is determined by the ***a*** and ***b*** parameters of ***Universal Mitotic Fraction Equation*** (#4), which lies within the ***Universal Growth Equation*** (#5). The ***c*** parameter, the ***Cell Cycle Time***, of the ***Universal Growth Equation*** (#5) describes how fast it takes cells to divide.

The ***Universal Growth Equation's*** description of how fast we grow is linked to the ***Cell Cycle Time***, ***c***, which is linked to genome size, which is linked to how much junk DNA we have.

What might be the mechanism behind the ***Cell Cycle Time***, the ***c*** parameter of the ***Universal Growth Equation*** (#5), which not only determines how long it takes a cell to divide, but also determines the speed of the growth of the body as a whole? Biochemically, the principal determinant of the ***Cell Cycle Time***, ***c***, is the amount of DNA that the cell contains. The more DNA a cell has to copy, the longer it takes to divide. This was first found by Van't Hof and Sparrow, who discovered a linear relationship between the total amount of amount of DNA and ***Cell Cycle Time*** for plant cells.[66] Their observation has subsequently been confirmed in many studies of many types of organisms.[67,68] For most animals, the largest part of the genome is non-coding DNA, sometimes called junk DNA,[69] no doubt imprecisely named.[70,71] The amount of this non-coding DNA has been found to be correlated with growth in salamanders,[72] anurans,[73] amphibians,[74] insects,[75] and copepods.[76]

The speed of growth has a profound impact on survival.[77,78] Human fetuses that grow to full-size in 10 months, or in 8 months, have a much lower chance of survival than fetuses that reach optimal size in 9 months.[79] Non-coding DNA, which determines genome size, is very susceptible to duplication or deletion, and thus provides populations of animals abundant genetic variation in genome size.[80-82] Perhaps this genetic variation in genome size leads to genetic variation in the ***Cell Cycle Time***, ***c***, which leads to genetic variation in the speed of body growth. Such genetic variation would appear to give species a powerful resource to draw upon, so that they can evolve, by the bitter reality of Darwinian selection, to growth rates that give them the greatest chance of survival. Indeed, this makes us wonder whether junk DNA may owe its very existence to its role in determining the speed of growth.

The ***Universal Growth Equation's*** description of the fraction of cells dividing, the ***Mitotic Fraction***, ***m***, is linked to the action of cell signaling molecules that induce mitotic quiescence.

What might be the mechanism behind the ***Mitotic Fraction***, ***m***, the fraction of cells that divide, which declines as we increase in size? What could cause a fraction of the body's cells to enter into a state of ***mitotic quiescence***, the internal block to cell division, which is set in motion by external signaling molecules?[60-62] Curiously, as we shall see below, this decline in the ***Mitotic Fraction***, by the ***Universal Mitotic Fraction Equation*** (#4), $\boldsymbol{m = a^{(N_w^b)}}$, might well trace its origin to the deceptively simple and unobvious discrete allocation of ligand molecule among cells[83]. To picture how this might occur, consider the case of an embryo growing in a constant volume, such as a bird's egg or mammal's uterus, whose cells produce an inhibitory growth factor and carry a receptor for that inhibitory molecule. Early in development, when our idealized embryo is just a few cells, those few cells would produce only a small number of inhibitory molecules, and thus the concentration of these inhibitory molecules would be low, and thus very few cells would have bound the number of inhibitory molecules needed to prevent cell division. Thus, at the beginning of development, most cells would divide, with the value of the ***Mitotic Fraction***, ***m***, being close to **1**. However, as the embryo grows, more and more cells are present to produce inhibitory molecules, the concentration of these inhibitory molecules would increase, more and more cells would have bound the number of inhibitory molecules needed to prevent cell division, fewer and fewer cells would be able to divide, and the value of the ***Mitotic Fraction***, ***m***, would decline. Remarkably, as we shall see next, when we rephrase this idealized case in mathematical terms, such a decline in the ***Mitotic Fraction***, ***m***, can be seen to occur in exactly the form of the ***Universal Mitotic Fraction Equation*** (#4), $\boldsymbol{m = a^{(N_w^b)}}$ .

Let us consider growth factor molecules in terms of integers.[83] Thus, when growth factor molecules bind to cells, they shall display a discrete allocation among cells, that is, as a Poisson process, which can be comprehended in terms of a Poisson probability (Equation #611 below).

Let **Ic** be the number of inhibitory growth factor molecule made by each cell which makes the molecule. (This number is a constant, being the result of the rates of synthesis and decay, most likely being first order reactions.)

Let $\boldsymbol{N}$ be the number of cells in the organism. We shall consider growth from conception, that is, from when $\boldsymbol{N = 1}$.

Let $\boldsymbol{N^b}$ be the number of cells producing the inhibitory growth factor molecules. If every cell produces the molecules, $\boldsymbol{b = 1}$. If only the cells on the surface produce the molecule, $\boldsymbol{b = \frac{2}{3}}$, since this captures the relationship between the surface of an isomorphic 3-dimensional shape, such as a sphere, and its volume.

Let $\boldsymbol{V}$ be the volume in which the embryo develops. (For a bird $\boldsymbol{V}$ is the volume of the egg; for a mammal it is the volume of the uterus.)

The total number of molecules of **I** in the body is $\boldsymbol{Ic \cdot N^b}$**.**

**[I]** is the concentration of inhibitory growth factor molecule in the body, assuming uniform diffusion.

Let us now put all this into mathematical form.

It follows that:

$$[\mathbf{I}] = \frac{Ic \cdot N^b}{V} \tag{601}$$

$$[\mathbf{I}] = \left(Ic \cdot N^b\right)\left(\frac{1}{V}\right) \tag{602}$$

Let $\mathbf{P_i}$ be the probability of a receptor molecule binding an inhibitory growth factor molecule, where **[R]** is the concentration of receptor molecules.

$$\mathbf{P_i} = \frac{[\mathbf{IR}]}{[\mathbf{R}]} \tag{603}$$

$$\frac{[\mathbf{I}][\mathbf{R}]}{[\mathbf{IR}]} = \mathbf{k} \tag{604}$$

according to the Law of Mass Action, thus

$$\frac{[\mathbf{IR}]}{[\mathbf{R}]} = \frac{[\mathbf{I}]}{\mathbf{k}} \tag{605}$$

$$\mathbf{P_i} = [\mathbf{I}] \cdot \frac{1}{\mathbf{k}} \tag{606}$$

$$\mathbf{P_i} = \left(\mathbf{Ic} \cdot \mathbf{N^b}\right)\left(\frac{1}{\mathbf{V}}\right)\left(\frac{1}{\mathbf{k}}\right) \tag{607}$$

$$\mathbf{P_i} = \mathbf{N^b}(\mathbf{Ic})\left(\frac{1}{\mathbf{V}}\right)\left(\frac{1}{\mathbf{k}}\right) \tag{608}$$

$$\mathbf{P_i} = \mathbf{N^b}\left(\frac{\mathbf{Ic}}{\mathbf{V \cdot k}}\right) \tag{609}$$

$$\mathbf{P_i} = \left(\frac{\mathbf{Ic}}{\mathbf{V \cdot k}}\right)\mathbf{N^b} \tag{610}$$

Let **Ro** be the number of receptors per cell.

We shall use the Poisson probability[84] to calculate the value of $\boldsymbol{m}$:

$$\mathbf{P(x;\mu)} = \frac{(\mathbf{e^{-\mu}})(\mathbf{\mu^x})}{\mathbf{x!}} \tag{611}$$

where **x** is the actual number of successes that result from the experiment.

**e**: A constant equal to approximately 2.71828

**μ**: The mean number of successes that occur in a specified region.

**x**: The actual number of successes that occur in a specified region.

$\mathbf{P(x;\mu)}$: The Poisson probability that exactly **x** successes occur in a Poisson experiment, when the mean number of successes is **μ**.

It follows that

$$\boldsymbol{m} = \mathbf{P(x;u)} = \frac{(\mathbf{e^{-\mu}})(\mathbf{\mu^x})}{\mathbf{x!}} \tag{612}$$

It follows that $\boldsymbol{x = 0}$ (since we want to calculate the chance of a cell binding **0** molecules of inhibitory growth factor).

And where $\mathbf{\mu = Ro \cdot P_i}$ (since the mean number of successes for each cell is the number of receptors, time the chance that a molecule will bind one of those receptors):

$$\boldsymbol{m} = \mathbf{P(x;\mu)} = \frac{(\mathbf{e^{-\mu}})(\mathbf{\mu^0})}{\mathbf{0!}} \tag{613}$$

$$\boldsymbol{m} = \mathbf{P(x;\mu)} = \frac{\mathbf{(e^{-\mu})(1)}}{\mathbf{1}} \tag{614}$$

$$\boldsymbol{m} = \mathbf{P(x;\mu)} = \mathbf{e^{-\mu}} \tag{615}$$

$$\boldsymbol{m} = \mathbf{P(x;\mu)} = \mathbf{e^{-Ro \cdot P_i}} \tag{616}$$

or

$$\boldsymbol{m} = \mathbf{e^{-RoP_i}} \tag{617}$$

It follows that

$$\boldsymbol{m} = \mathbf{e}^{\mathbf{-Ro}\left(\frac{\mathbf{Ic}}{\mathbf{V \cdot k}}\right)\mathbf{(N^b)}} \tag{618}$$

Let $\boldsymbol{Z} = \boldsymbol{Ro}\left(\frac{\boldsymbol{Ic}}{\boldsymbol{V}\cdot\mathbf{k}}\right)$

$$\boldsymbol{m} = \mathbf{e^{-Z(N^b)}} \tag{619}$$

Let $\boldsymbol{a} = \boldsymbol{e^{-Z}}$

$$\boldsymbol{m} = \boldsymbol{a}^{\mathbf{(N^b)}} \tag{4}$$

Which is the ***Universal Mitotic Fraction Equation***!

This leads to our ***Universal Growth Equation***, when combined with Equation #5a:

$$\frac{\boldsymbol{dN}}{\boldsymbol{dt}} = \frac{\mathbf{ln(2)}}{\boldsymbol{c}} \cdot \boldsymbol{N} \cdot \boldsymbol{a}^{(\boldsymbol{N^b})} \tag{5a}$$

where

$$\boldsymbol{a} = \mathbf{ln}\left(\boldsymbol{Ro}\left(\frac{\boldsymbol{Ic}}{\boldsymbol{V}\cdot\mathbf{k}}\right)\right) \tag{620}$$

Note that all of the biochemistry (the rate of growth factor synthesis, abundance of receptors, dissociation constants, etc.) is in the parameter ***a***, while all of the geometry (size of the tissues that making the growth factor in relationship to the size of the organism) is in the parameter ***b*** (see "Let $\boldsymbol{N^b}$ be …" above).

Having worked through Equation #601-620, let us step back and review the core implications of this math. This math is telling us that little more than the discrete allocation of mitotic signaling molecules among cells is all that is needed to explain how cell division becomes limited to a faction of cells, ***m***, the ***Mitotic Fraction***, as captured by the ***Universal Mitotic Fraction Equation*** (#2), and its two parameters ***a*** and ***b***, and from this, to cause our growth to occur by the ***Universal Growth Equation***. These mitotic signaling molecules, the receptors on the cells to which they bind, and the internal mitotic control molecules that respond to these signals, are all ruled by ordinary chemistry. Thus, little more than the conventional thermodynamics of all chemical reactions is sufficient to account for growth by the ***Universal Growth Equation***, and thus the biological process of ***UNI-GROWTH***. Let us also recall that since signaling molecules, their receptors, and the internal mitotic control molecules that determine whether cells divide, are the products of our genes, and thus genetic polymorphism in the value of ***a*** or ***b*** might well be expected to identify genetic polymorphisms of the proteins that do this mitotic signaling, a possibility that would certainly be worthy of experimental examination.

***UNI-GROWTH***: Summary Definition

Let us summarize: ***UNI-GROWTH*** is the process by which growth slows as we increase in size. ***UNI-GROWTH*** is captured by the ***Universal Mitotic Fraction*** and ***Growth Equations***, and their parameters, ***a***, ***b***, and ***c***, the average ***Cell Cycle Time***. ***UNI-GROWTH*** occurs by the decline in the fraction of cells dividing, the ***Mitotic Fraction***, occurring by the form of ***Universal Mitotic Fraction Equation***, ***m=a^(N^b)***. From this expression, we could then derive, and test, the ***Universal Growth Equation***, whose accuracy was confirmed, from fertilization until maturity, for 13 species, including nematodes, mollusks, amphibians, fish, birds, rodents, cows, and humans.

#### ***ALLO-GROWTH***

The ***Cellular Phylodynamic Analysis*** of the growth and development of the parts of the body

The ***Allometric Growth Equation***

For more than a century, biologists have been studying relative growth, that is, the growth of one part of the body, $\boldsymbol{p}$, against another part, $\boldsymbol{p_2}$, or against the body as a whole, $\boldsymbol{w}$.[15,18] These studies of relative growth, usually carried out in units of weight, volume, or length, had found that the relationships between $\boldsymbol{p}$ and $\boldsymbol{w}$ often form straight rows of dots on log-log graphs.[15-19] This log-linearity of relative growth, called ***allometric growth***[85], can be captured by:

$$\log(\boldsymbol{w}) = \frac{\boldsymbol{1}}{\boldsymbol{S}} \cdot \log(\boldsymbol{p}) + \log(\boldsymbol{B}) \tag{630}$$

which is equivalent to:

$$\boldsymbol{w} = \boldsymbol{B} \cdot \boldsymbol{p}^{\frac{\boldsymbol{1}}{\boldsymbol{S}}} \tag{631}$$

Here, we shall call ***B*** the "***allometric birth***" and ***S*** the "***allometric slope***". Below, it will become clear why we have named the parameters in this fashion, as their meanings emerge when relative growth is framed in terms of numbers of cells, ***N***.

While the empirical validity of the log-linearity of allometric growth has been evident from countless studies of the growth of many parts of the body, in many animals,[15-19] the meaning of this phenomenon has long been a mystery, as have been the biological meanings of the parameters ***B*** and ***S***.[19,86,87] However, as we shall outline next, when we frame the relationship of the growth of one part of the body, $\boldsymbol{p}$, against the body as a whole, $\boldsymbol{w}$, in terms of the number of cells in the part of the body, $\boldsymbol{N_p}$, against the number of cells in the body as a whole, $\boldsymbol{N_w}$, the biological basis and significance of the log-linear feature of allometric growth becomes evident, together with the biological meanings of the two parameters that define it, ***B*** and ***S***.

##### ***Cellular Allometric Growth Analysis***

The approach that we have taken to deciphering the nature of the relative growth of each part of the body is the same as the approach we described above for examining the growth of the body as a whole: ***Cellular Phylodynamic Analysis***, by which we again mean examining growth in terms of integer numbers of cells, ***N***. To do so, we examined the sizes of the various parts of the body, $\boldsymbol{p}$, in these integer units of numbers of cells, $\boldsymbol{N_p}$, in comparison to the numbers of cells in the body as a whole, $\boldsymbol{w}$, also in these integer units of numbers of cells, $\boldsymbol{N_w}$ , an approach we call "***Cellular Allometric Growth Analysis***" (FIGURES A8-A17). Graphically, such a ***Cellular Allometric Growth Analysis*** of relative growth is carried out by examining paired $\boldsymbol{N_p}$:$\boldsymbol{N_w}$ values on log-log graphs (FIGURES A8-A17).

When we have a full lineage chart for an animal, which characterizes each cell from the first fertilized egg onward, we call this "***Cellular Allometric Lineage Growth Analysis.***" When we are carrying this out without lineage data, but simply with cell numbers, whether these cell numbers have been counted, or estimated from weight values, based on the finding that there are about $10^8$ cells in every gram of tissue,[57] we call this approach "***Cellular Allometric Batch Growth Analysis.***"

###### Early development: ***Cell lineages***: ***Cellular Allometric Lineage Growth Analysis***:

To carry out a ***Cellular Phylodynamic Analysis*** of the relative growth of the various parts of the body early in development, we assembled values of $\boldsymbol{N_p}$ vs. $\boldsymbol{N_w}$ for each of the ***cell lineages*** (FIGURE A8) that have been characterized in the ***cell lineage charts*** of ***Caenorhabditis elegans***[20] and ***Meloidogyne incognita***[21] nematode worms, as well as for ***Oikopleura dioica*** chordate tunicates (FIGURES A9-A11).[88] To accomplish such ***Cellular Allometric Lineage Growth Analysis***, we simply counted the number of cells in each ***lineage***, $\boldsymbol{N_p}$, from the first ***Founder Cell*** onward ($\boldsymbol{N_p} = 1$), together with the corresponding values for the number of cells in the embryo as a whole, $\boldsymbol{N_w}$ (FIGURES A9-A11).

Such a ***Cellular Allometric Lineage Growth Analysis*** was easily carried out for the ***M incognita***, whose ***cell lineage chart*** fits on a single sheet of paper (FIGURES A9). However, ***Cellular Allometric Lineage Growth Analysis*** was considerably more difficult for ***C elegans*** and ***O dioica***, whose ***cell lineage charts*** have to be 6 feet long to see each cell, requiring many months of work to assemble the values of $\boldsymbol{N_p}$ and $\boldsymbol{N_w}$ shown in FIGURES A10 AND A11. This motivated the development of a new technique, which we shall describe below, the ***BinaryCellName Method***, for carrying out such ***Cellular Allometric Lineage Growth Analysis*** computationally.

###### Late development: ***Tissues***, ***Organs***, and ***Anatomical structures***: ***Cellular Allometric Batch Growth Analysis***:

To carry out a ***Cellular Phylodynamic Analysis*** of the relative growth of the various parts of the body later in development, that is, ***Cellular Allometric Batch Growth Analysis***, we assembled values of $\boldsymbol{N_p}$ vs. $\boldsymbol{N_w}$ for the ***tissues***, ***organs***, and ***anatomical structures*** evident in embryos and juveniles from published weight and volume values, again relying on the observation that there are about $10^8$ cells in every gram of tissue.[57] We assembled these values for the body parts of chicks (gizzards, livers, hearts, kidneys),[89] rats (livers, brains, kidneys, forelegs, ears, stomachs, spinal cords),[90] humans (brains, livers, kidneys, lungs, pancreases, adrenals, thymuses, spleens, lower extremities, upper extremities, stomach, heart, intestines),[91] zebrafish (eye lens),[92-95] mice (livers, brains, kidneys, forelegs),[90,96] clams and goldfish (FIGURES A12-A17).

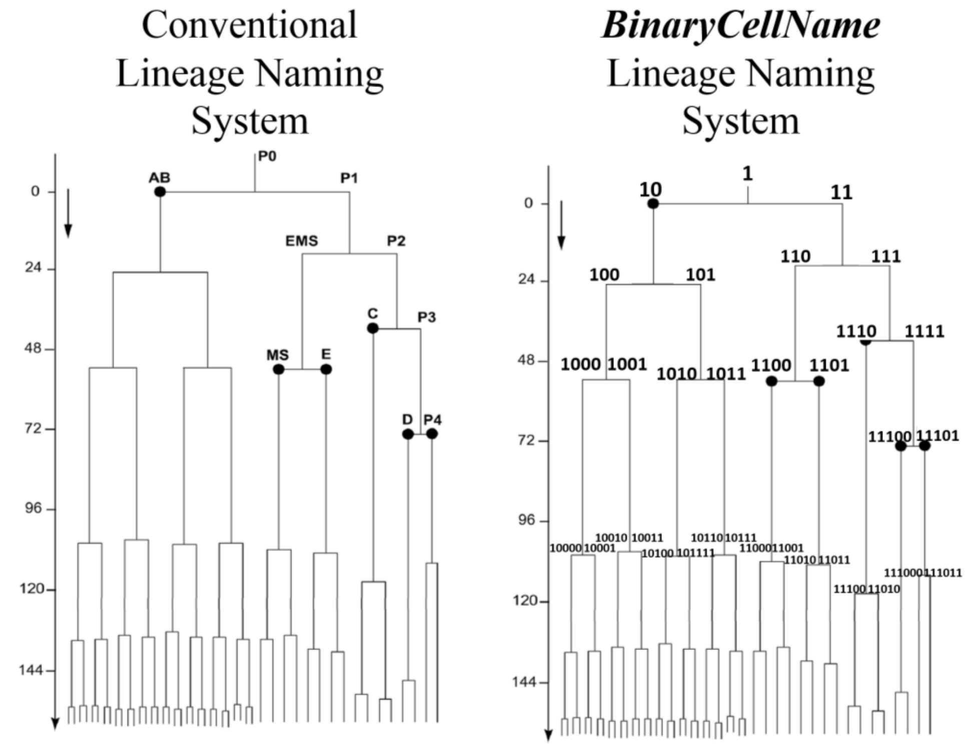


**FIGURE A8**: ***Cell lineage chart*** **of the** ***cell lineages*** **of the developing root knot nematode,** ***Meloidogyne incognita*****. Shown are the conventional and** ***BinaryCellName Lineage*** **naming methods. The AB** ***lineage*** **of nematodes forms the skin while the E** ***lineage*** **forms the intestine.** ***Cell lineage chart*** and data from Calderón-Urrea et al[21]

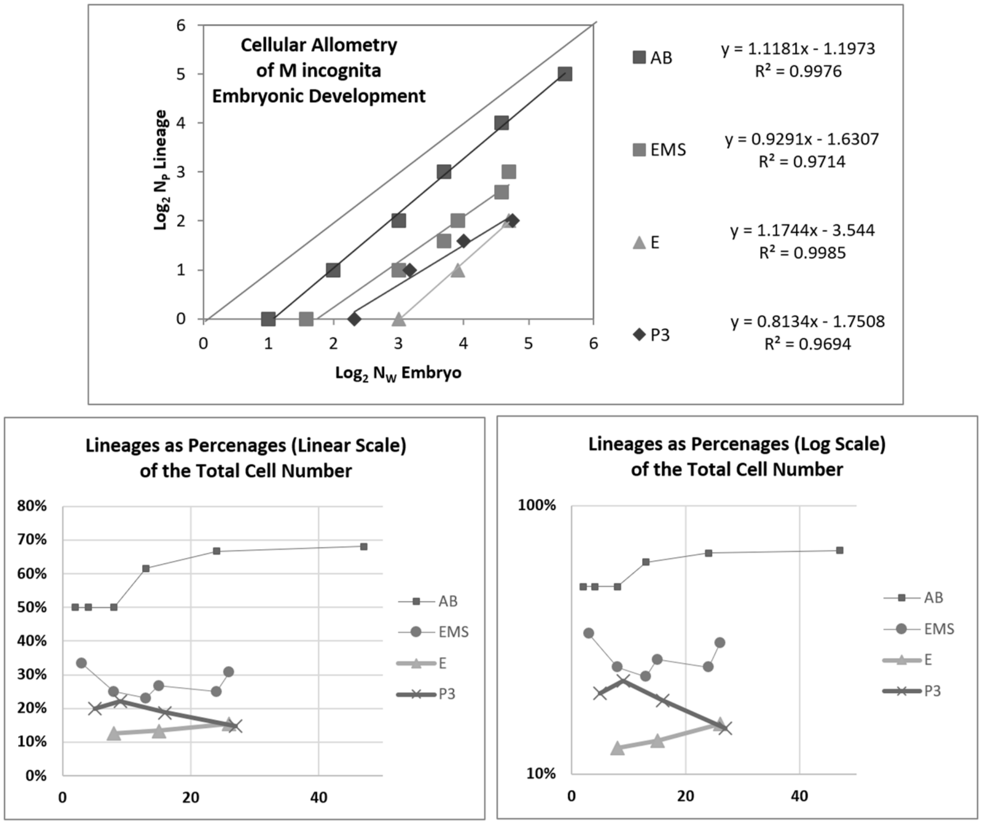


**FIGURE A9**: ***Cellular Allometric Lineage Growth Analysis*** **of the** ***cell lineages*** **of the developing root knot nematode,** ***Meloidogyne incognita*****.**[21]

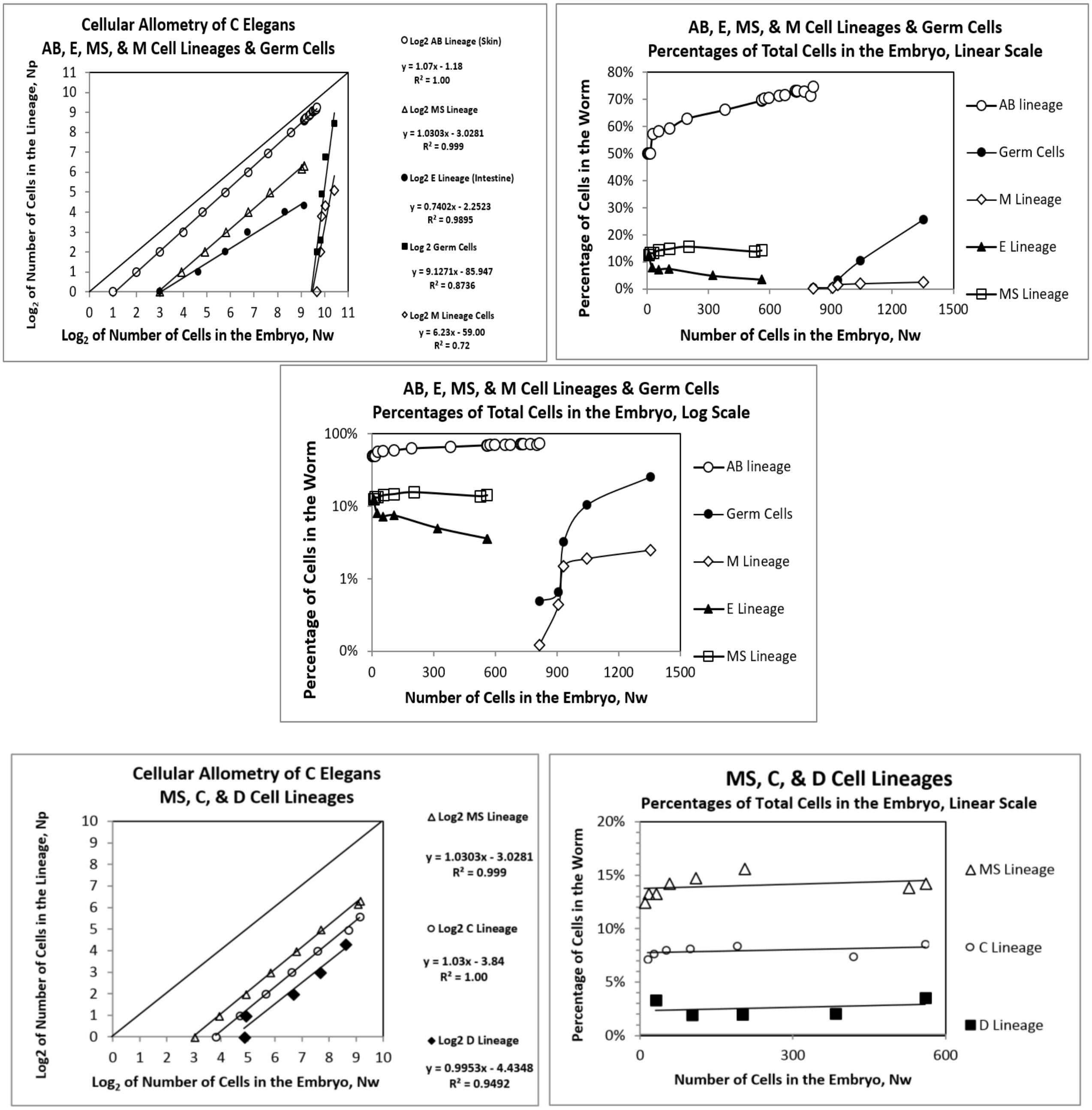


**<u>FIGURE A10</u>**: ***Cellular Allometric Lineage Growth Analysis*** **of the** ***cell lineages*** **of the developing** ***C elegans*** **nematode**

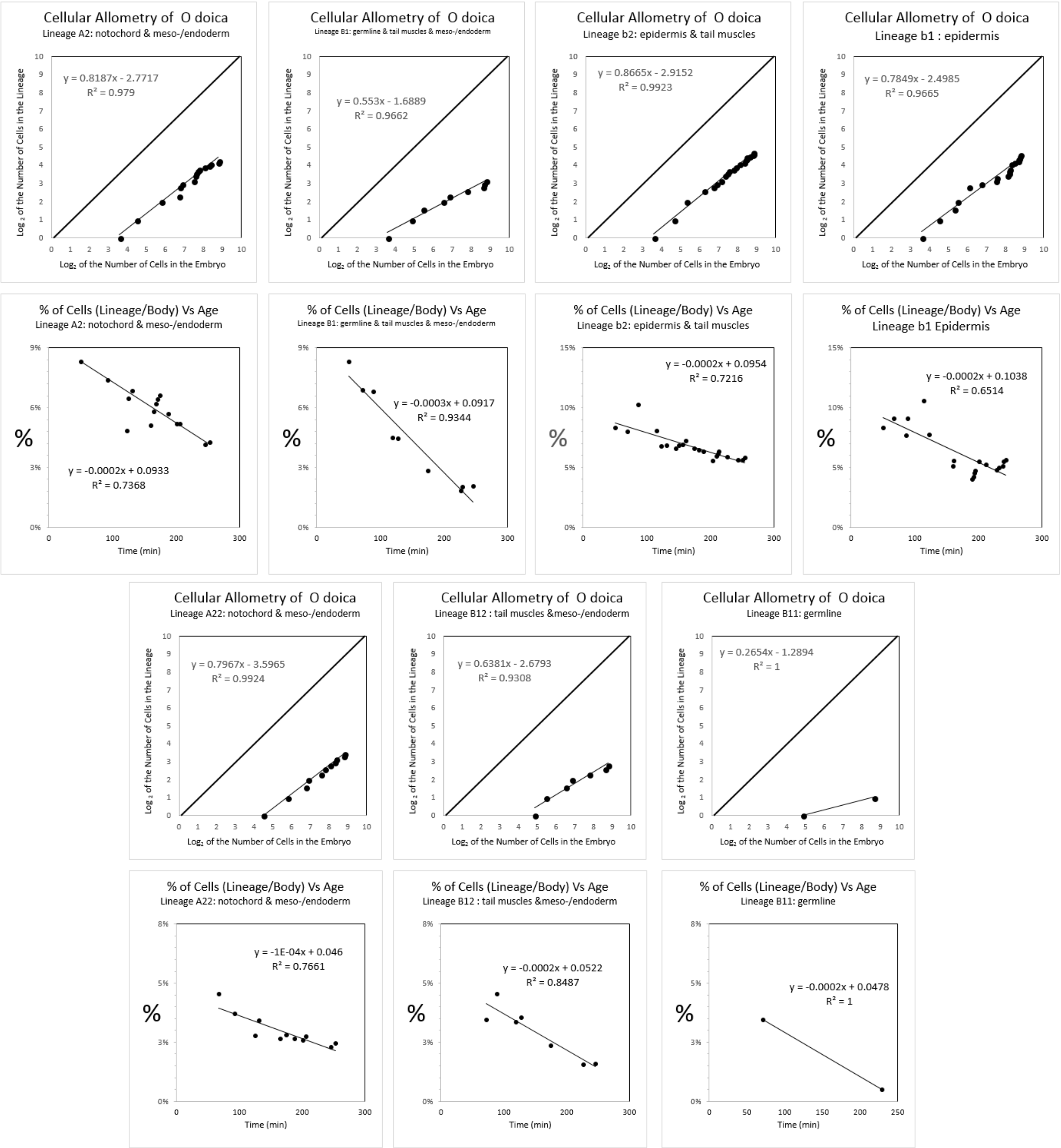


**<u>FIGURE A11</u>:** ***Cellular Allometric Lineage Growth Analysis* of the *cell lineages* of the chordate tunicate *Oikopleura dioica*.** Unlike ***C elegans*** and ***M incognita***, all of the cases of ***Cellular Selection*** for this species reflect reductions in cell number, although this could be due to the limited size of the dataset rather than a biological difference. ***Cell lineage chart*** data from Stach et al[88]

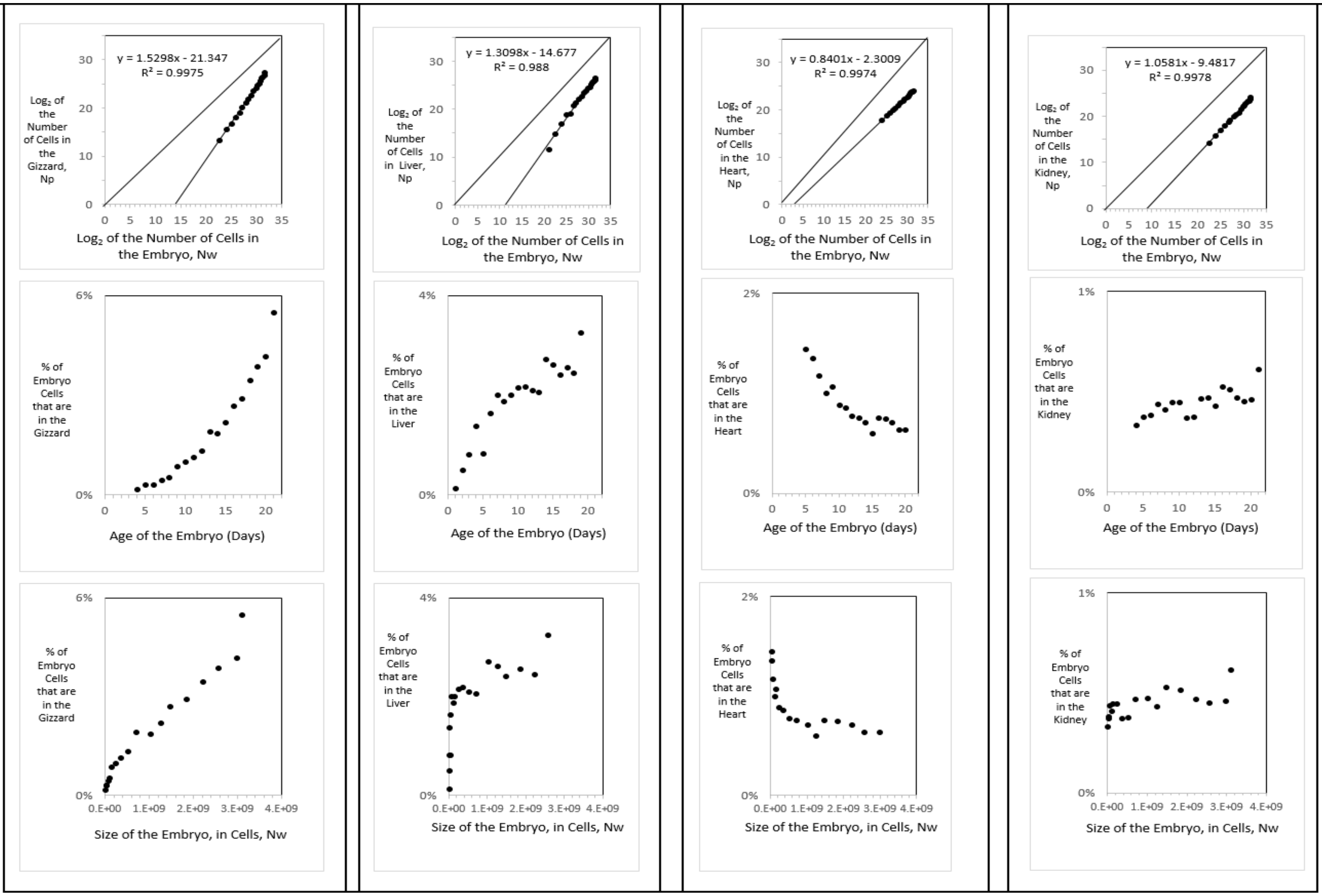


| | % | age | Size of Embryo in Cells | % | age | Size of Embryo in Cells, N | % | age | Size of Embryo in Cells, N | % | age | Size of Embryo in Cells, N |
|---|---|---|---|---|---|---|---|---|---|---|---|---|
| smallest value | 0.17% | 4 | 6.E+06 | 0.14% | 3 | 2.E+06 | 1.35% | 6 | 3.E+07 | 0.33% | 4 | 6.E+06 |
| largest value | 5.49% | 21 | 3.E+09 | 3.26% | 21 | 3.E+09 | 0.64% | 20 | 3.E+09 | 0.61% | 21 | 3.E+09 |
| % increase absolute | 5.32% | | 3.E+09 | 3.12% | | 3.E+09 | -0.72% | | 3.E+09 | 0.28% | | 3.E+09 |
| % increase relative | 33 | | 516 | 23 | | 1475 | 0.47 | | 87 | 2 | | 516 |
| absolute increase relative | 16,990 | | | 33,633 | | | 41 | | | 950 | | |

**FIGURE A12**: ***Cellular Allometric Batch Growth Analysis* of *tissues, organs,* and *anatomical structures* of the chick embryo.** Growth of the parts of the chick embryonic gizzard, liver, heart, and kidney, in units of numbers of cells, in comparison to the number of cells in the embryo as a whole shown on log-log graphs, showing how the fraction of cells comprising in the gizzard and liver increases as the number of cells in the embryo increase with development and how the fraction of cells comprising in the heart decreases as the number of cells in the embryo increase with development, and how the fraction of cells comprising in the kidney remains roughly constant as the number of cells in the embryo increases with development.

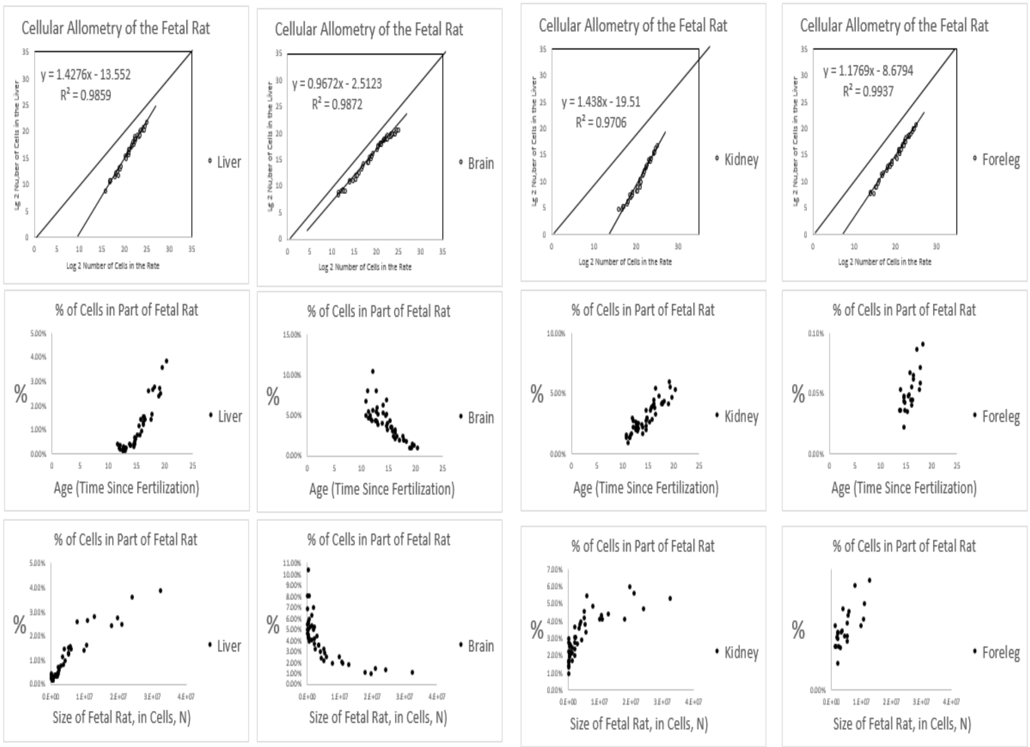


| | % | age | Size of Embryo in Cells, N |
|---|---|---|---|
| first value | 0.46% | 12 | 5.E+04 |
| last value | 3.89% | 20 | 3.E+07 |
| % increase absolute | 3.43% | | 3.E+07 |
| % increase relative | 8.51 | | 589 |
| te increase relative | 5,012 | | |

| | % | age | Size of Embryo in Cells, N |
|---|---|---|---|
| first value | 6.92% | 11 | 7.E+04 |
| last value | 1.10% | 20 | 3.E+07 |
| % decrease absolute | -5.82% | | 3.E+07 |
| % decrease relative | 6.31 | | 5.E+02 |

| | % | age | Size of Embryo in Cells, N |
|---|---|---|---|
| first value | 1.70% | 11 | 7.E+04 |
| last value | 5.37% | 20 | 3.E+07 |
| % increase absolute | 3.67% | | 3.E+07 |
| % increase relative | 3.16 | | 5.E+02 |
| absolute increase relative | 1,549 | | |

| | % | age | Size of Embryo in Cells, N |
|---|---|---|---|
| first value | 0.04% | 14 | 1.E+06 |
| last value | 0.21% | 20 | 3.E+07 |
| % increase absolute | 0.17% | | 3.E+07 |
| % increase relative | 5.62 | | 29 |
| absolute increase relative | 162 | | |

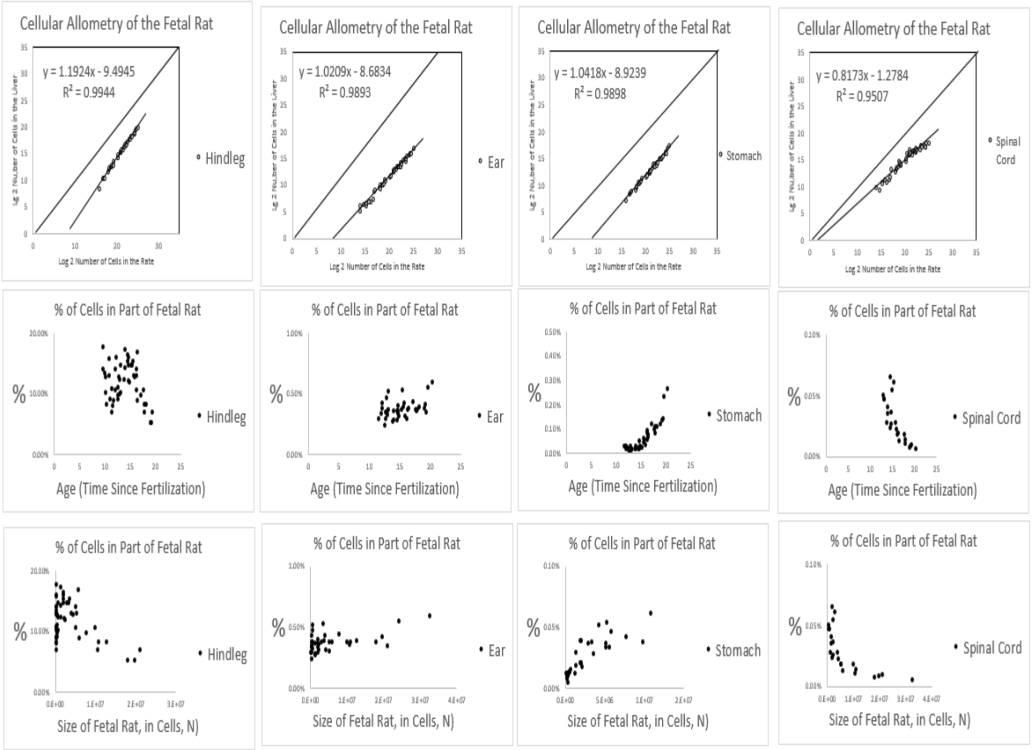


| | % | age | Size of Embryo in Cells, N |
|---|---|---|---|
| first value | 17.78% | 12 | 5.E+04 |
| last value | 5.25% | 20 | 3.E+07 |
| % decrease absolute | -12.53% | | 3.E+07 |
| % decrease relative | 3.39 | | 589 |

| | % | age | Size of Embryo in Cells, N |
|---|---|---|---|
| first value | 0.30% | 12 | 5.E+04 |
| last value | 0.60% | 20 | 3.E+07 |
| % increase absolute | 0.30% | | 3.E+07 |
| % increase relative | 2.00 | | 589 |
| absolute increase relative | 1.175 | | |

| | % | age | Size of Embryo in Cells, N |
|---|---|---|---|
| first value | 0.03% | 12 | 5.E+04 |
| last value | 0.27% | 20 | 3.E+07 |
| % increase absolute | 0.24% | | 3.E+07 |
| % increase relative | 8.13 | | 589 |
| absolute increase relative | 4.786 | | |

| | % | age | Size of Embryo in Cells, N |
|---|---|---|---|
| first value | 0.05% | 12 | 5.E+04 |
| last value | 0.01% | 20 | 3.E+07 |
| % decrease absolute | -0.04% | | 3.E+07 |
| % decrease relative | 0.13 | | 589 |

**FIGURE A13**: ***C̄ellular Allometric Batch Growth Analysis* of tissues, organs, and anatomical structures of the rat embryo**Growth of body parts of the rat embryonic liver, brain, kidney and foreleg, in units of numbers of cells, in comparison to the number of cells in the embryo as a whole, are shown on log-log graphs. Note how the fraction of cells comprising in the liver kidney and foreleg increases as the number of cells in the embryo increase with development, while the fraction of cells comprising the brain decreases.

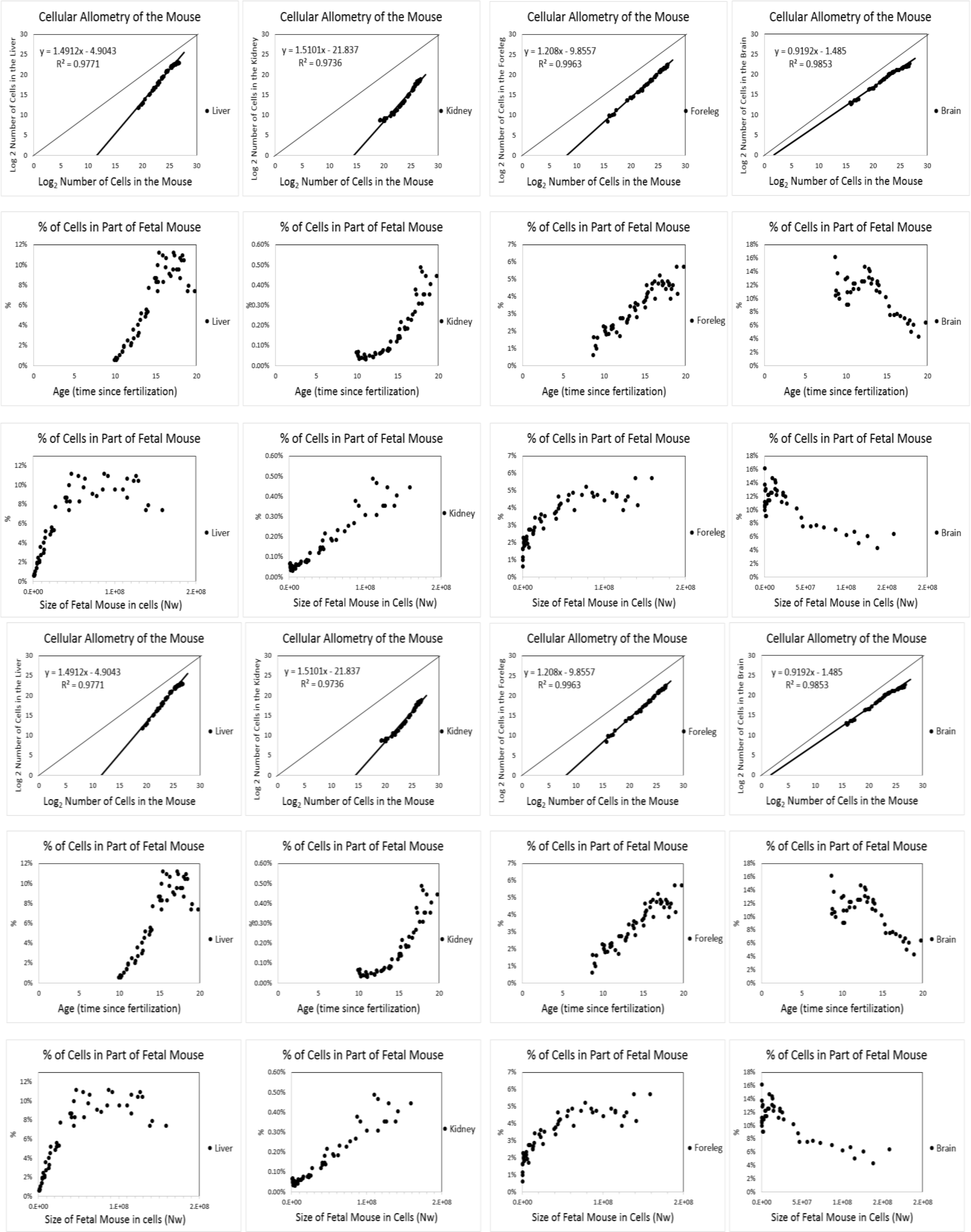


**FIGURE A14**: ***Cellular Allometric Batch Growth Analysis*** **of tissues, organs, and anatomical structures of the mouse embryo.**

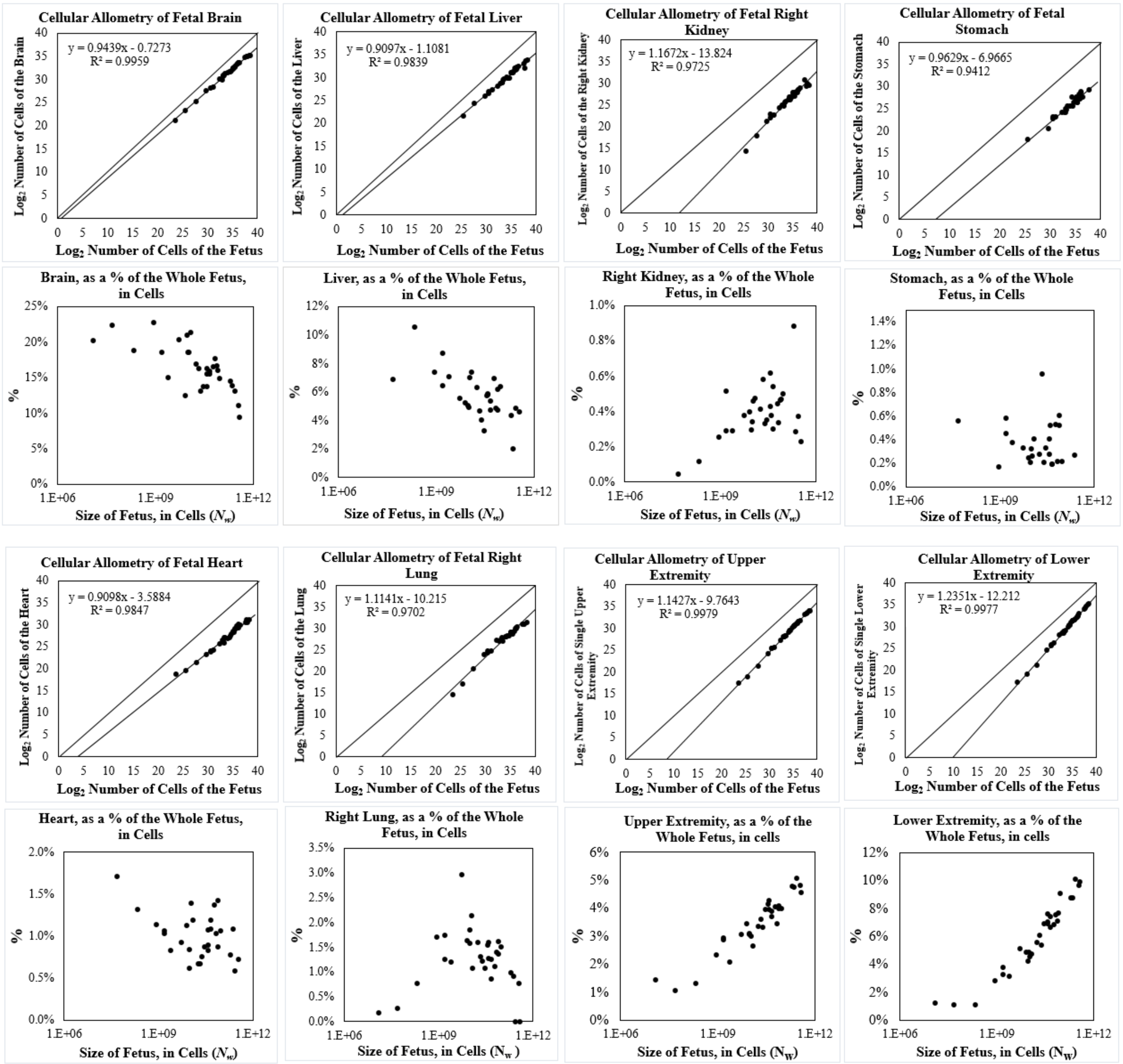


**FIGURE A15**: ***Cellular Allometric Batch Growth Analysis*** **of tissues, organs, and anatomical structures of the developing human fetuses, from autopsy data, of individual fetuses**.

For additional human relative growth data, see FIGURE A16

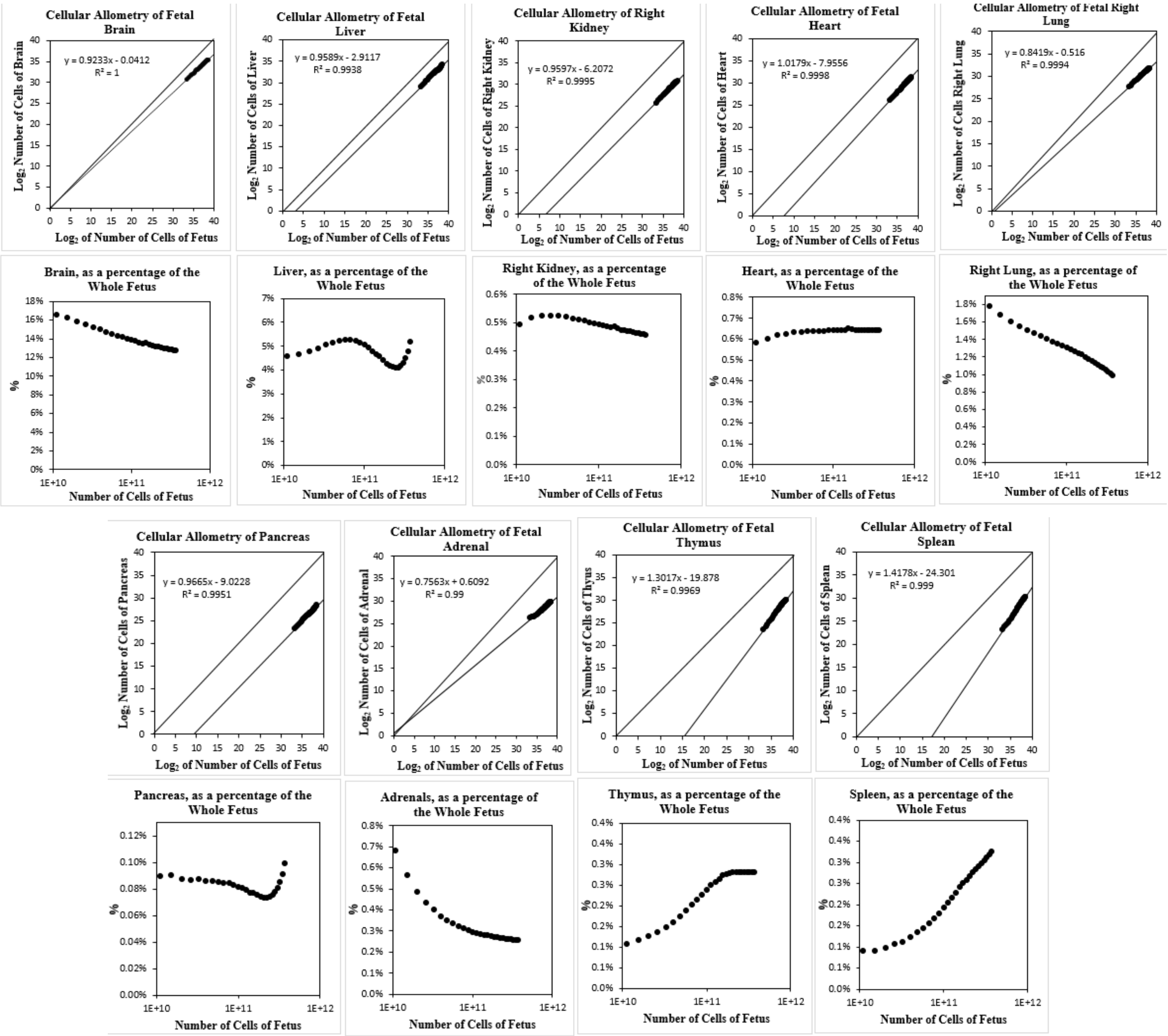


**FIGURE A16**: ***Cellular Allometric Batch Growth Analysis*** **of tissues, organs, and anatomical structures of the developing human fetuses, from autopsy data, averaged values**.[91,97]

For additional human relative growth data, see FIGURE A15

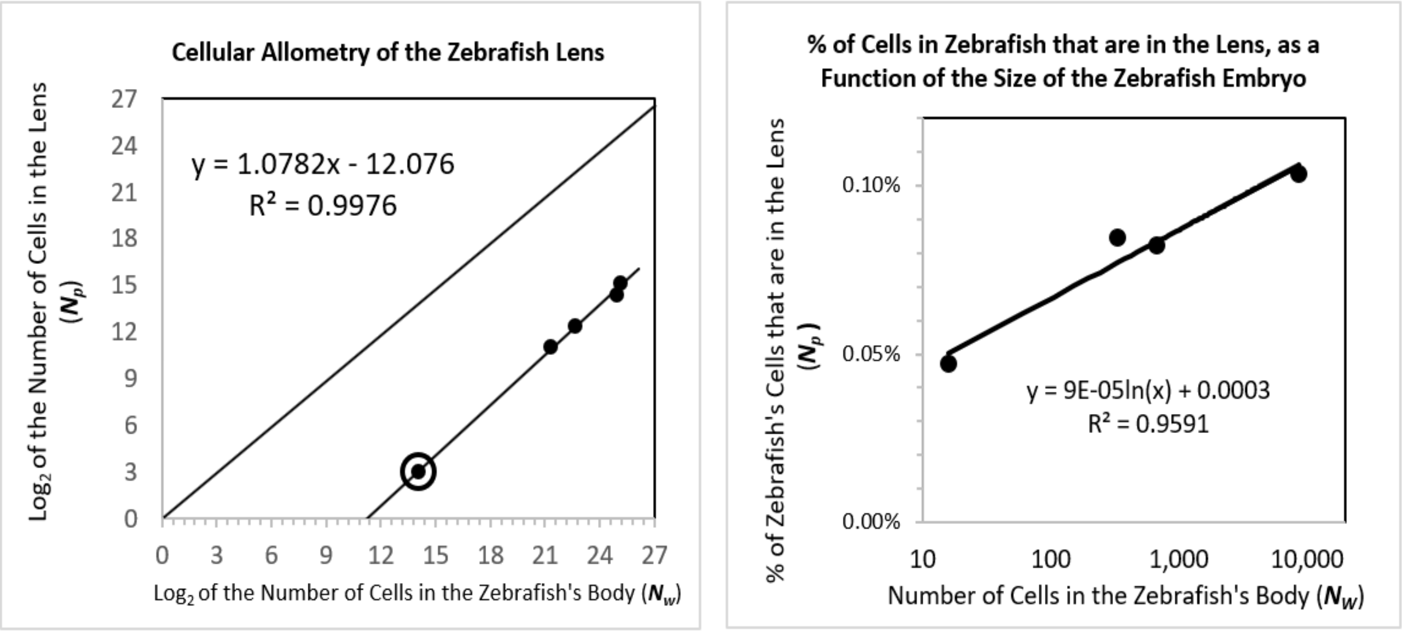


**<u>FIGURE A17</u>: *Cellular Allometric Batch Growth Analysis* of the Zebrafish Lens.**

The ***Cellular Allometric Growth Equation***

With the data outlined above, we were able to compare, on log-log graphs, the number of cells in various parts of the body (***cell lineages***, ***tissues***, ***organs***, and ***anatomical structures***), $N_p$, with the number of cells in the embryo as a whole, $N_w$. Remarkably, in the great majority of these cases, comparisons of $N_p$ vs. $N_w$ appeared as ramrod-straight rows of dots on log-log graphs (FIGURES A9-A17). The high $r^2$ values attest to the strength of these observations of cellular log-linearity. These empirically based observations make plain that ***relative body-part growth***, when examined in numbers of cells, $N_p$ vs $N_w$, often takes the form of the ***Allometric Equation***:

$$\log(N_w) = \frac{1}{S_N} \cdot \log(N_p) + \log(B_{N1}) \tag{4}$$

which is equivalent to:

$$N_w = B_N \cdot N_p^{\frac{1}{Sn}} \tag{4b}$$

We call these expressions "***Cellular Allometric Growth Equations***" (FIGURE A18), and the biological process which is captured "***ALLO-GROWTH***". We call the parameter $B_{N1}$ the "***Cellular Allometric Birth***"' and $S_N$, the "***Cellular Allometric Slope***".

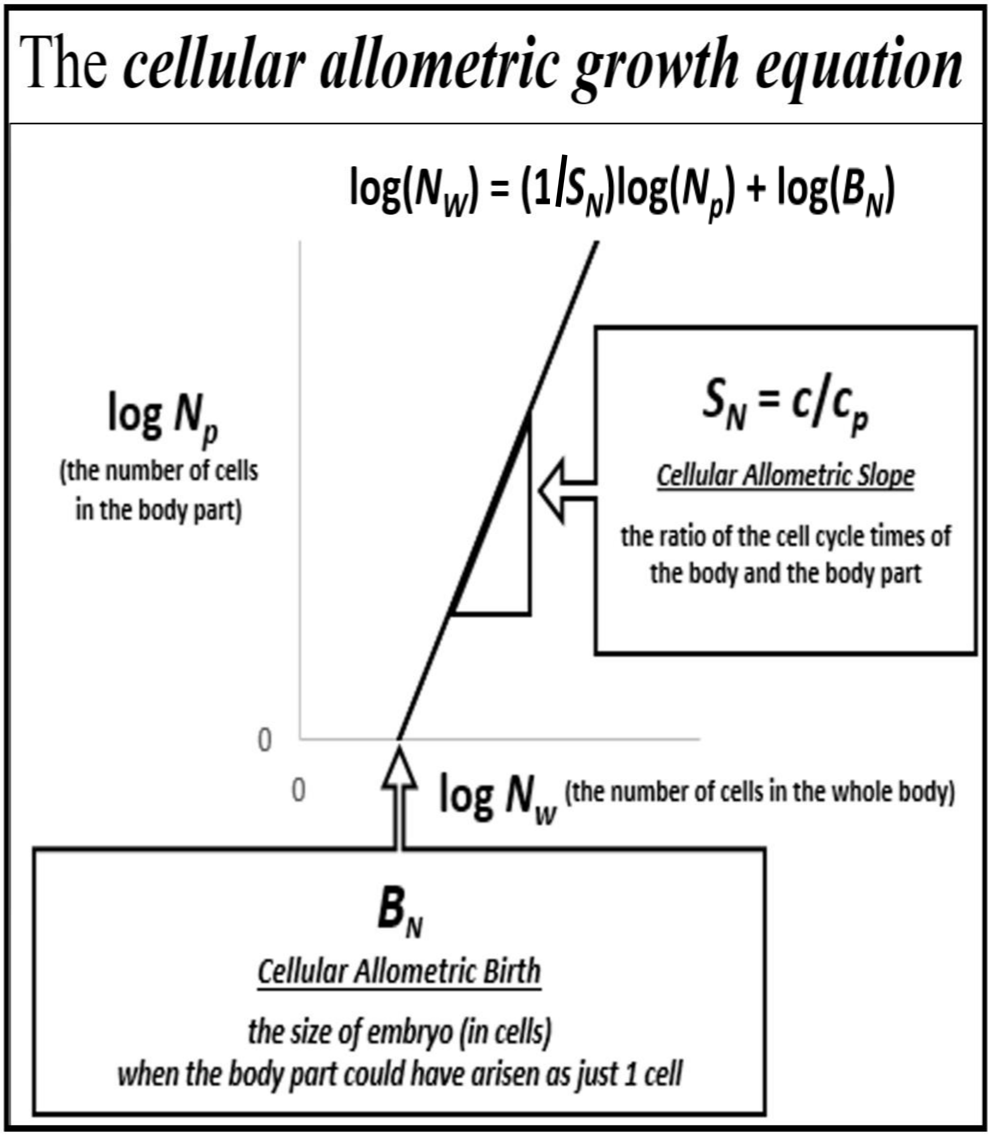


**FIGURE A18: The *Cellular Allometric Growth Equation*, log($N_w$) = (1/$S_N$) log($N_p$) + log($B_{N1}$).**

Body part ***ALLO-GROWTH*** results from whole body ***UNI-GROWTH***.

Recall that the ***Cellular Allometric Growth Equation*** captures the rate of growth of the whole embryo, while the rate of growth for each body-part is:

$$\frac{dN_p}{dt} = \frac{ln(2)}{c_p} \cdot N_p \cdot (m_p) \quad (650)$$

where $c_p$ is the ***Cell Cycle Time***, and $m_p$ the ***Mitotic Fraction*** of the cells of the body part. Thus, the ratio of the rate of growth of the body to the body-part is:

$$\frac{dN_w}{dN_p} = \left[\frac{ln(2)}{c} \cdot N_w \cdot (m)\right] \cdot \left[\frac{ln(2)}{c_p} \cdot N_p \cdot (m_p)\right]^{-1} \text{, where } m = a^{(N_w^b)} \text{ and } m_p = a_p^{(N_w^{bp})} \quad (651)$$

Consider the case where $m_p$=$m$ and $c_p$= $c$, that is, where a body part has the same ***Mitotic Fraction*** and ***Cell Cycle Time*** as the embryo, and starts sometime later than the first cell:

$$\frac{dN_w}{N_w} = \frac{dN_p}{N_p} \quad (652)$$

Integration[1] reveals:

$$log(N_w) = \log(N_p) + \log(B_N) \quad (652)$$

which is the allometric relationship, with $S_N$=**1.** Thus, growth of a part of the body by the ***Cellular Allometric Growth Equation*** is the consequence of ***Founder Cells*** giving rise to cell lineages in animals growing by the ***Universal Growth Equation***.

Parameter $S_N$, the ***Cellular Allometric Slope*** of the ***Cellular Allometric Growth Equation***, tells us how the embryo uses the ***Cell Cycle Time*** to drive ***Cellular Selection*** to adjust the size of each part of the body:

The ***Cellular Allometric Slope***, $S_N$, can be traced to the ***Cell Cycle Time***, $c_p$.

Now, let us consider the case where $m_p$=$m$ but $c_p \neq c$, that is, where a body part has the same ***Mitotic Fraction*** as the embryo but a different ***Cell Cycle Time***:

$$\frac{dN_w}{N_w} = \frac{c_p}{c} \cdot \frac{dN_p}{N_p} \quad (653)$$

Integration reveals:

$$log(N_w) = \frac{c_p}{c} \cdot \log(N_p) + \log(B_N) \quad (653)$$

where:

$$S_N = \frac{c}{c_p} \quad (655)$$

which is the allometric relationship, with $S_N$,≠**1**. Thus, $S_N$, the ***Cellular Allometric Slope*** of the ***Cellular Allometric Growth Equation***, can be traced to the ***Cell Cycle Time***, $c_p$ in each part of the body (FIGURE A18).

On the other hand, similar analysis revealed the ***Mitotic Fraction,*** $m_p$, that is, a change in the fraction of cell dividing, to be an unlikely source for allometric log-linearity. Indeed, changes in the value of either ***a*** or ***b*** of the ***Universal Mitotic Fraction Equation*** (#4), which sets the value of $m_p$, does not result in straight lines on log-log graphs but in curves (not shown).

The ***Cellular Allometric Slope***, $S_N$, and ***Cellular Selection***.

The value of $S_N$, the ***Cellular Allometric Slope*** of the ***Cellular Allometric Growth Equation***, appears on log-log graphs at a measure of whether a part of the body is growing faster ($S_N$>**1**), slower ($S_N$<**1**), or at the same speed ($S_N$=**1**) as the body as a whole. Thus, $S_N$, the ***Cellular Allometric Slope*** of the ***Cellular Allometric Growth Equation*** (#7b) (FIGURE A18) shows how embryos adjust the size of each body part by differential cellular proliferation. These changes in the value of $c_p$ are small, and thus have minor impact on the average ***Cell Cycle Time*** in the embryo as a whole, ***c***, but can have dramatic cumulative impacts on the relative sizes of body parts (FIGURES A8-A17). Thus ***ALLO-GROWTH***, and its mathematical abstraction, the ***Cellular Allometric Growth Equation*** (#7), distills ***Cellular Selection***, that is, differential cellular proliferation,[98,99]down to a single value, $S_N$. Embryos accomplish this by ***Cell-Heritable*** change the ***Cell Cycle Time***, $c_p$.

---

[1] For a superb YouTube discussion of why the integration of Equation #10 that leads to Equations #7c and #7d, which Salman Khan asserts is "one of the two coolest derivatives in all of calculus", see:
https://www.khanacademy.org/math/in-in-grade-12-ncert/in-in-advanced-differentiation-two/copy-of-proofs-for-derivatives-of-ex-and-lnx-ab/v/proof-d-dx-ln-x-1-x-old
and
https://www.khanacademy.org/math/ap-calculus-ab/ab-antiderivatives-ftc/ab-common-indefinite-int/v/antiderivative-of-x-1

Parameter $S_N$, the ***Cellular Allometric Slope***, at work in adjusting the size of each part of the body

What does the ***Cellular Allometric Slope***, $S_N$, of the ***Cellular Allometric Growth Equation*** (FIGURE A18) tell us about how embryos create our anatomy? The value of $S_N$, the ***Cellular Allometric Slope***, provides us with a measure of how the embryo adjusts the sizes of each of our parts by ***Cellular Selection***, that is, differential cellular proliferation, a remarkably subtle but powerful force in shaping the composition of the embryo.[98,99] We see in $S_N$, the ***Cellular Allometric Slope***, nothing less than the determination of the size of the embryo's ***cell lineages***, ***tissues***, ***organs***, and ***anatomical structures***, by ***Cellular Selection***, driven by the ***Cell-Heritable***, average ***Cell Cycle Time***, $c_p$, of the cells in each of these parts of the body. Thus, the ***Cellular Allometric Growth Equation*** allows us to distill the action of the fundamental morphogenetic force of ***Cellular Selection*** down to a single value, $S_N$.

Parameter $S_N$, the ***Cellular Allometric Slope***, at work early in development

Early in development, we can see ***Cellular Selection*** at work in molding the composition of the first structures of the embryo, the ***cell lineages***. For example, note in FIGURE A10 how the nematode ***C elegans* AB *lineage***, which forms the skin of the worm, has a ***Cellular Allometric Slope***, $S_N$, slightly greater than 1 ($S_N \approx 1.1$); this results in the **AB** ***lineage*** growing from 50% of the cells of the worm at its creation in the **AB** ***Founder Cell*** to almost 70% of the worm's body at hatching (FIGURE A10). In the other direction, the ***C elegans* E *lineage***, which forms the intestine, also grows as a straight line on a log-log graph, but with a ***Cellular Allometric Slope***, $S_N$, of less than 1 ($S_N \approx 0.7$); this results in the **E** intestinal ***lineage*** declining from about 12% of the cells of the ***C elegans*** when it first arises from its **E** ***Founder Cell*** to about 3% at hatching (FIGURE A10). Even more dramatically, the ***C elegans*** germ ***cell lineage*** rises by log-linear ***Cellular Selection*** from a single cell, comprising about 0.02% of the embryo. to about 20% of the adult, roughly a 100-fold increase, through the action of a ***Cellular Allometric Slope***, $S_N$, of ~9 (FIGURE A10). A similar dramatic log-linear post-embryonic increase can be seen for the ***C elegans* M *lineage***, which form muscle cells (FIGURE A10). Many other ***cell lineages*** display such ***Cellular Selection***, as can be seen in FIGURES A9 and A11.

Parameter $S_N$, the ***Cellular Allometric Slope***, at work late in development

Later in development, we again see ***Cellular Selection*** at work molding body part size, this time for ***tissues***, ***organs***, and ***anatomical structures*** (FIGURES A12 - A17). For example, note in FIGURE A12 how the chick's liver starts out as just 0.17% of the cells of the embryo 2 days after the egg is laid, but has grown, with a ***Cellular Allometric Slope***, $S_N$, of ~1.3, to about 5.5% of the embryo by day-21, a 33-fold increase. In the other direction, the chick's heart contains just 1.35% of the embryo's cells at day 2, declining with a ***Cellular Allometric Slope***, $S_N$, of ~0.84, to about 0.64% of the embryo by day-21, a ~2-fold decrease (FIGURE A12). Such ***Cellular Selection*** can be seen to be at work in molding the sizes of many of the ***tissues***, ***organs***, and ***anatomical structures*** of animals, whose log-log graphs can be seen in FIGURES A12 - A17. We found this log-linearity of ***Cellular Selection*** in the relative growth of these embryonic structures of developing mice, rats, clams, and goldfish (data not shown), and human embryos as well (FIGURES A15 and A16).

Parameter, $S_N$, the ***Cellular Allometric Slope***, $S_N$, and the ***Cell Cycle Time***, $c_p$,

Where does the ***Cellular Selection*** captured by the ***Cellular Allometric Slope***, $S_N$, come from? By expanding ***Cellular Allometric Growth Equation*** (#8c), we have been able to see above that the underlying cause of ***Cellular Selection*** can be traced to the average ***Cell Cycle Time*** of the cells in a part of the body, $c_p$, in comparison to the average ***Cell Cycle Time*** of the cells in the body as a whole, $c$, with this expression, which we repeat here for convenience:

$$S_N = \frac{c}{c_p} \qquad (655)$$

In short, the ***Cellular Allometric Growth Equation***, and the relative growth data that it summarizes, tell us that our bodies adjust the size of our body-parts by adjusting the average ***Cell Cycle Time***, $c_p$, of the cells in each of our parts, in a ***Cell-Heritable*** fashion. The ***Cellular Allometric Growth Equation*** is also pointing to possible times when this specification of body part size occurs: the ***Cellular Allometric Birth***. These graphs shown in FIGURES A9-A17, empirical summaries of actual data, of many body parts, in many animals, are also pointing to ***Cellular Allometric Births*** for virtually all body parts, whether large or small, as mapping to very early in development. This suggests that many parts of the body could arise from small numbers of cells, experiencing cell heritable changes in the ***Cell Cycle Time***, $c_p$, very early in development.

The changes in the ***Cell Cycle Time***, $c_p$, that occur within us, and which drive the changes in relative growth that we have seen in the ***Cellular Allometric Slope***, $S_N$, from all of the $N_p$ vs. $N_w$ comparisons we have carried out (FIGURES A9-A17), are generally very small, just a few percentage points. Indeed, these change in the ***Cell Cycle Time***, $c_p$, that occur within the body are tiny in comparison to the hundred-fold differences in the ***Cell Cycle Times***, $c$, which we have seen between different species of animals (Table 2). Nonetheless, these small internal changes in the ***Cell Cycle Time*** can accumulate, often leading to dramatic changes in the body's composition, such as the 33-fold increase noted above in the size of the liver in chick embryos caused by an $S_N$ value of ~1.3.

The role of DNA methylation in the change in the ***Cell Cycle Time***, $c_p$ in body parts

What could cause such a ***Cell-Heritable*** change in ***Cell Cycle Time*** that lies behind the ***Cellular Allometric Slope***, $S_N$, and drives the ***Cellular Selection***, and thus determine the sizes of our ***cell lineages***, ***tissues***, ***organs***, and ***anatomical structures***? There are a number of ***Cell-Heritable*** biological processes that influence the time it takes for a cell to divide, of which DNA methylation has been the most widely studied; methylated DNA takes longer to copy, and thus causes cells to take more time to divide.[100,101]

When might DNA methylation, or some similar ***Cell-Heritable*** process, set the value of the ***Cell Cycle Time***, $c_p$, and thus the ***Cellular Allometric Slope***, $S_N$, of the cells of a part of the body, and thus, ultimately, the size of that part? An appealing possibility for consideration is the moment when that part could have been created, is it arose form a single cell, that is, at its ***Cellular Allometric Birth*** $B_{N1}$, when the part could have been but a single cell. Should this be found to be the case, then the ***Cellular Allometric Birth*** $B_{N1}$, would appear to be the moment when both the part, and its size, are determined by the embryo.

The Parameter, $B_{Na}$, the ***Cellular Allometric Birth*** of the ***Cellular Allometric Growth Equation***, tells us how embryos create body parts from single cells

The value of $B_{Na}$, the ***Cellular Allometric Birth*** of the ***Cellular Allometric Growth Equation***, can be seen easily on log-log graphs (FIGURE A18), as the place where the ***Cellular Allometric Growth Equation*** crosses the ***x***-axis, and thus where $\log(N_p) = 0$, and, therefore, where $N_p = 1$.

Should a part of the body come into existence as a single ***Founder Cell***, the ***Cellular Allometric Birth***, $B_{N1}$, corresponds to the number of cells in the body of the embryo, $N_w$), when that single ***Founder Cell*** arose by mitosis (i.e. $B_N = N_w$ when $N_p = 1$). In fact, this is precisely what has been seen for those animals for which we have ***cell lineage charts***, which have been collected to describe every cell in the embryo from the zygote onward (FIGURE A8), that is, for nematode worms ***Caenorhabditis elegans***[20] and ***Meloidogyne incognita***[21], and for tunicates, ***Oikopleura dioica***,[88] which are chordates quite closely related to ourselves.

For the ***tissues***, ***organs***, and ***anatomical structures*** for which we have growth data later in development, and for ***drosophila*** body parts which we shall discuss below, we seldom have data on their cellular origins. There has been much scholarship on this point, but, fortunately, modern light sheet 4D microscopy should allow us to answer these questions,[22,23,102] and the ***Cellular Allometric Growth Equation*** points us to where we should look. Furthermore, whether a body part arises from a single ***Founder Cell***, or multiple ***Founder Cells***, ***Cellular Phylodynamic Analysis*** allows us to characterize the features of the process. For example, should a part of the body come existence from ***z Founder Cells***, each of which is born at the same time, and has the same ***Cell-Heritable Cell Cycle Time***, $c_p$, the part made of these ***z Founder Cells*** will also grow by the ***Cellular Allometric Growth Equation***. Even more complex such examples are amenable to this ***Cellular Phylodynamic Analysis*** approach, a topic we shall address below.

When the parts of the body are born: The ***Cellular Allometric Birth in Time***, $B_T$

The ***Cellular Allometric Growth Equation*** tells us that embryos create our body parts from individual ***Founder Cells***. When does this occur? In terms of the size of the embryo, each part is born when the size of the embryo is $B_{N1}$ ($B_N = N_w$ when $N_p = 1$). In terms of the time when each part is born (relative to the time of fertilization, when $t = 0$), which we call the "***Cellular Allometric Birth in Time***", $B_T$, we can reach back to the ***Universal Growth Equation*** (#5):

$$B_T = \left[\mathrm{Ei}\left(-B_N^b \, log(a)\right) - \mathrm{Ei}(-\,log(a))\right] \frac{c}{b \; log(2)} \tag{660}$$

where $B_T$ is the time of a ***Cellular Allometric Birth***, when a ***Founder Cell*** becomes a body part.

Furthermore, early in development, when most ***Founder Cells*** appear, growth is quite close to exponential, as most cells are dividing. Thus, building from Equation #2:

$$B_T \approx c \cdot (ln(B_N)) \cdot [ln2]^{-1} \tag{661}$$

The time of the ***Cellular Allometric Birth*** of a body part affects its size.

An interesting functional correlate of the single cell origin of ***cell lineages*** is that the structures they form can be big or small by having earlier or later ***Cellular Allometric Births***, $B_{N1}$. For example, while the **MS**, **C**, and **D** ***lineages*** of ***C elegans*** all grow on log-log graphs right along with the embryo as a whole, thus with ***Cellular Allometric Slopes***, $S_N$, close to **1**, the cells of the **MS** ***lineage*** make up about 12%-14% of the worm because it is born at the 8-cell stage ($B_N = 8$), while the cells of the **C** ***lineage*** make up about 8% of the worm by being born at about the 16-cell stage ($B_N = 16$), and the cells of the **D** ***lineage*** makes up about 3% of the worm by being born at about the 32-cell stage ($B_N = 32$) (FIGURE A11). Thus, the option of generating structures from single cells at various points in the growth of the body as a whole gives the embryo another trick for adjusting the relative sizes of its many parts.

Data on the number of ***Founder Cells*** that make various parts of the body.

Early in development, we can see the creation of body parts from single ***Founder Cells*** for those animals for which ***cell lineage charts*** have been made. For example, note in FIGURES A9 and A10, how the nematode ***C elegans* AB *Founder Cell*** arises when the embryo is just 2 cells in size, whose progeny go on to form the **AB *lineage***, which forms the skin from the ***Cellular Allometric Birth*** of $\boldsymbol{B_N = 2}$ (i.e., $\boldsymbol{N_w = 2}$ when $\boldsymbol{N_p = 1}$). Similarly, note in FIGURES A9 and A10, how the ***C elegans* E *Founder Cell*** arises when the embryo is 8 cells in size, whose progeny go on to form the **E *lineage***, which forms the intestine from the ***Cellular Allometric Birth*** of $\boldsymbol{B_N = 8}$ (i.e., $\boldsymbol{N_w = 8}$ when $\boldsymbol{N_p = 1}$). Many other such examples of the creation of body parts from single ***Founder Cells*** can be seen for ***C elegans*** and ***M incognita*** nematode worms and ***O dioica*** chordate tunicates in FIGURES A9 and A10.

Later in development, we get hints of the creation of body parts from single ***Founder Cells***, as the ***Cellular Phylodynamic Analysis*** of the ***tissues***, ***organs***, and ***anatomical structures*** of the body shows that the line of the ***Cellular Allometric Growth Equation*** points down to the spot on the $\boldsymbol{x}$-axis where the ***Cellular Allometric Birth***, $\boldsymbol{B_{N1}}$, lies (FIGURES A12-A17). Remarkably, in almost every case, the line of the ***Cellular Allometric Growth Equation*** points to a ***Cellular Allometric Birth***, $\boldsymbol{B_{N1}}$, that corresponds to the number of cells in the body of the embryo, $\boldsymbol{N_w}$ when the embryo was quite early in its development, and never to numbers of cells less than **2**. For example, as can be seen in FIGURE A12, $\boldsymbol{B_N \approx 8}$ for the chicken heart, suggesting that the heart could have arisen from just 1 cell when the embryo was ~8 cells in size. For the chicken liver, $\boldsymbol{B_N \approx 2,000}$, suggesting that the liver could have arisen from just 1 cell when the embryo was ~2,000 cells in size (FIGURE A12). Many other such examples of the creation of body parts from single ***Founder Cells*** can be seen in the relative growth of these embryonic structures of developing mice, rats, clams, and human embryos as well (FIGURES A15 and A16). While few of these large structures have been measured down to the $\boldsymbol{B_{N1}}$ intersection point, the zebrafish eye lens comes close, as Greiling and Clark documented its growth from just 8 cells, when the embryo is 16 hours post fertilization, with the lens's ***Cellular Allometric Growth Equation*** pointing down to a ***Cellular Allometric Birth*** when the embryo was about 2000 cell in size, and thus $\boldsymbol{B_N \approx 2,000}$ when $\boldsymbol{N_p = 1}$ (FIGURE A17).[93-95] These observations complement other studies that have also suggested that large anatomical structures may arise from small numbers of cells,[103,104] including recent CRISPR/Cas9 ***cell lineage*** labeling studies.[105,106]

Body parts can be made of one ***Founder Cell*** or more than one ***Founder Cell***

Curiously, whether a part of the body arises from **1 *Founder Cell***, or more than **1 *Founder Cell***, ***ALLO-GROWTH*** will result in body parts growing by the ***Cellular Allometric*** and ***Allometric Growth Equations***, and, early in development, by the ***Exponential Growth Equation***, if all ***Founder Cells*** of a body part arise at the same time and with the same ***Cell Cycle Time***, $\boldsymbol{c_p}$. This could be seen in a simple calculation of an idealized part of the body that is made from **3 *Founder Cells***, arising early in development, when growth is close to exponential (FIGURE A19). As can be seen in the top two graphs of FIGURE A19, if all three ***Founder Cells***, arise at the same time, and have the same ***Cell Cycle Time***, $\boldsymbol{c_p}$, their growth together will be indistinguishable from a part made of **1 *Founder Cell*** (once it gets up to 3+ cells!). Furthermore, as can be seen in the other graphs of FIGURE A19, no matter when the three ***Founder Cells*** are born, or what their ***Cell Cycle Times***, $\boldsymbol{c_p}$, are, the aggregate growth of the part will still appear as a quite straight row of datapoints on a log graph. Thus, the fact that the growth of a body part points back to a single ***Cellular Allometric Birth***, doesn't mean that it arose from a single cell; we still have to do the microscopy work to see what actually happened.[22,23,93,153,107,153]

This simple excise also shows, however, that body parts made of more than a single ***Founder Cell*** do have some challenges. Note for example, that ***Founder Cells*** born after the first ***Founder Cell***, or having a lower ***Cell Cycle Time***, $\boldsymbol{c_p}$, than the fastest growing ***Founder Cell***, will soon become irrelevant, and the body part will become functionally monoclonal. A ***Founder Cell*** born three ***Cell Cycle Times***, $\boldsymbol{c}$, after the first ***Founder Cells***, which in a human or a mouse is about three days, will make up only about 10% of the part, while a ***Founder Cell*** born eight ***Cell Cycle Times***, $\boldsymbol{c}$, after the first ***Founder Cell*** will make up only about 1% of the part. ***Founder Cells*** that have ***Cell Cycle Times***, $\boldsymbol{c_p}$, slower than the fastest ***Founder Cell*** will soon loose in the race to make up a significant percentage of the part.

While these this simple exercise examined the idealized case of a part of the body arising early in development, when growth is close to exponential, the results shown here are likely to be similar later in development as well, when ***UNI-GROWTH*** begins to be felt, although exploring precise details for this would be worthwhile.

Of course, we can imagine all sorts of complicated multi-clonal anatomical structures. Perhaps their ***Cell Cycle Times*** are elegantly intertwined, or they adjusted their ***Mitotic Fractions*** to stay in league. Fortunately, however, whatever the cellular behavior, if one collects the cell numbers, ***Cellular Phylodynamic Analysis*** provides us with a way to isolate the underlying processes.

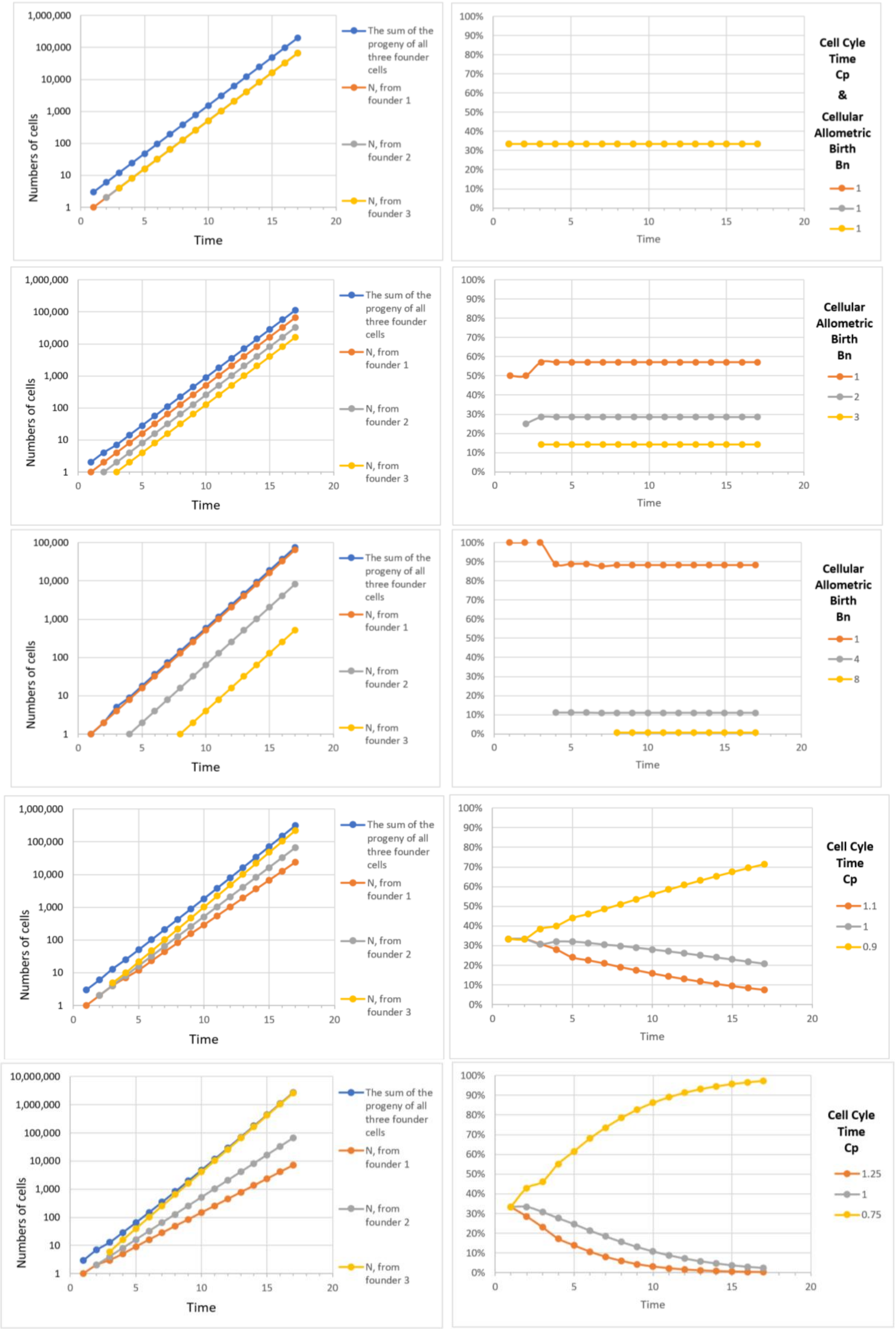


**<u>FIGURE A19</u>: Calculations for an idealized part of the body that is made from 3 *Founder Cells*, arising early in development**

#### The ***Cellular Phylodynamic Analysis*** of ***Drosophila Growth*** and ***Development***

To carry the ***Cellular Phylodynamic Analysis*** of drosophila growth and development, we assembled values for the numbers of cells in the fly as a whole, $N_w$, from fertilization onward, and in the fly's various parts, $N_p$ (FIGURE A20). Many of these parts grow from small structures, which contain few cells, including discs, histoblast nests, and pole cells (which will give rise to the fly's germ cells), as we have found in studies scattered throughout the literature over the past 80 years.[108-133]

##### Numbers of cells in the fly as a whole, $N_w$.

Measuring the size of the body in units of numbers of cells, $N_w$, and gauging growth from these values, is complicated by the quirky nature of drosophila growth, which is marked by periods of massive cell suicide, followed by regrowth from small structures, as well as by the mass polyploidization that goes on as flies develop. Nonetheless, the relationship between cell number, $N_w$, and age, $t$, has been quite accurately measured for the first 6000 cells by Zalokar and Erk (1975).[108] As we shall see, for our first look, we don't require precise values for $N_w$ beyond this point, but rough estimates can be made from measured of body mass by Sawala and Gould[109] (who kindly provided raw data) and from DNA content.[110]

##### Numbers of cells in the body parts of the fly, $N_p$.

There has been considerable uncertainty as to whether parts of the fly's body arise from single ***Founder Cells*** (clones) or multiple ***Founder Cells*** (polyclones)[111] addressed in many studies of challenging effort that used ***Cell-Heritable*** making techniques, such as gynandromorphs, or mitotic recombination, to track these parts to their origins. We shall return below to this area of scholarship, and its relationship to the work described here, once we have presented the findings based on the raw numbers of cells in the fly as a whole, $N_w$, and in its various parts, $N_p$, and age, $t$, from fertilization onward, based on cell counting studies.

Many of the body parts of flies have been observed from small numbers of cells (FIGURE A20). Both the spiracular anlage[114] and the pole cells (which form the germ cells)[108,120] have been seen from 2 cells onward (FIGURE 20). Genital imaginal discs have been observed from 2 cells,[112,113] while other discs have been observed from as few as 12 cells.

Superb data on the numbers of cell is imaginal discs and histoblast nests over periods of time have been reported in studies by Madhaven and Madhaven (1980),[114] Madhavan and Schneiderman (1977),[115] Graves and Schubiger (1982),[116] Bate and Arias (1991),[117] and Bryant and Levinson (1985).[118] These researchers reported that the number of cells, $N_p$, in antenna discs (33 cells to 48 cells), eye discs (40 cells to 250 cells), wing discs (24 cells to 4500 cells), haltere discs (12 cells to 63 cells), prothoracic leg discs (36 cells to 54 cells), mesothoracic leg discs (42 cells to 85 cells), metathoracic leg discs (45 cells to 89 cells), over various points in time from as early as 13 hours until 120 hours after fertilization. For histoblast nests, cell numbers, $N_p$, have been counted from as small as 4 cells.[114,119] The spiracular anlage has been tracked from 2 cells.[115] From 118 to 138 hours of life, we have data on the three anterior dorsal histoblast nests from 15 to 466 cells, on the three posterior dorsal histoblast nests from 5 to 110 cells, and on the three ventral nests, with numbers ranging from 11 to 284 cells.[115] Verma and Cohen[119] have provided videos that show the growth of these histoblast nests back to as few as 4 cells at the beginning of their videos; one wonders what would have been seen if one turned on the camera before then, and such a simple data collection exercise cries out for experimental examination. Furthermore, these videos make clear that nest cells increase in number by cell division rather than by recruiting unrelated cells. The population of pole cells, which gives rise to the germ cells of the fly, have been counted back to 2 cells by Rabinowitz (1941)[120] while Zalokar and Erk (1975)[108] have counted the number pole cells from 12 cells, together with the number of cells in embryo as a whole from the first cell until ~6000 cells. For discs below 20 cells in size, the literature contains some contradictory values,[114,115,117] which calls out for modern high resolution 4D microscopy[22,23,93,153,153] not available to these researchers 40+ year ago, but, for the projections we shall describe below, this does not present an impediment.

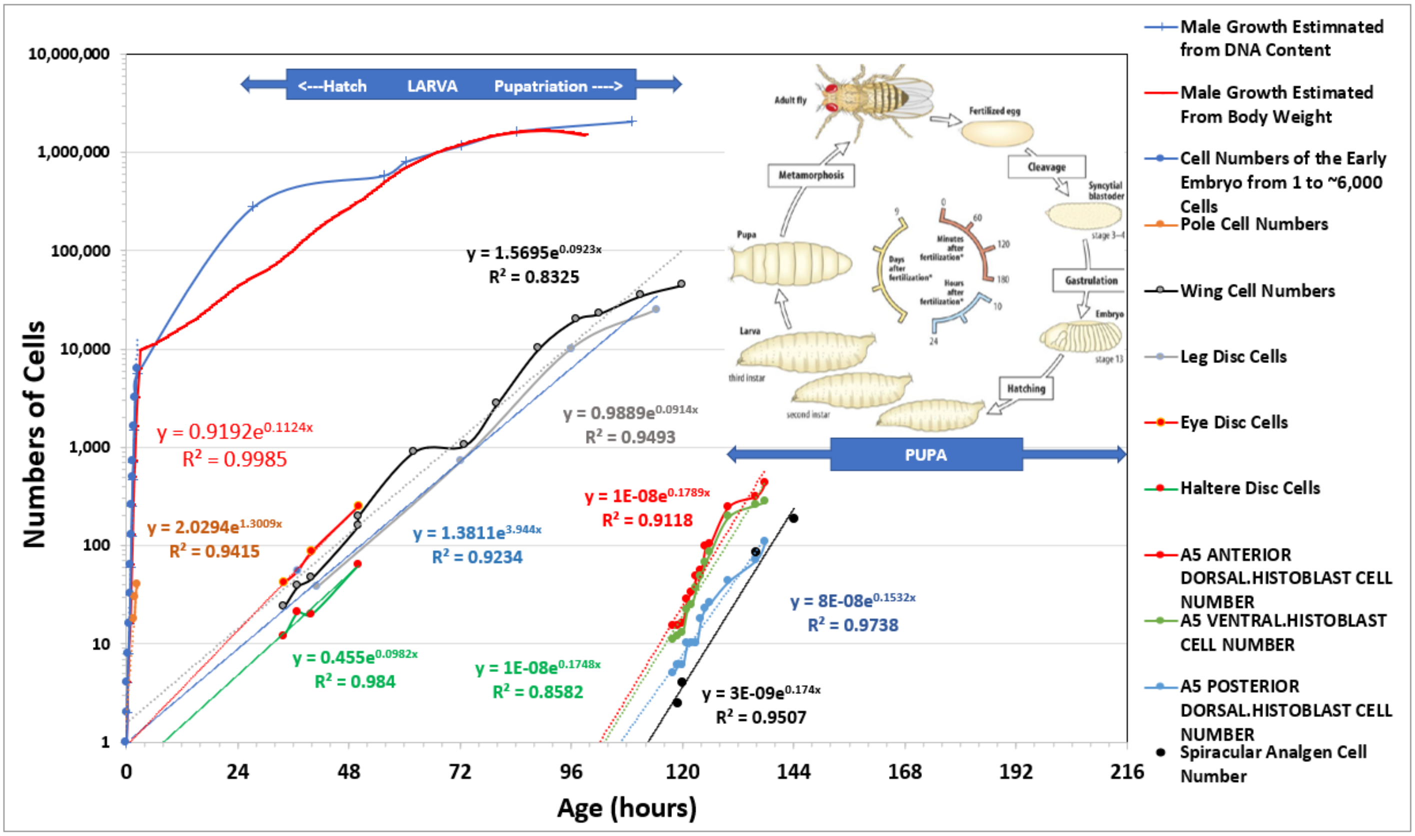


**<u>FIGURE A20:</u> The *Cellular Phylodynamic Analysis* of *Drosophila Growth* and *Development***

<u>Log-Linearity of the increase in the numbers of cells in various body parts of the fly, $N_p$.</u>

When examined on log graphs of cell number, $N_p$, versus time, $t$ (where $t$=**0** when $N_w$=**1**, FIGURE A20), most of the parts of the fly's body display striking exponential growth, which appears as straight-rows of datapoints on log-graphs, below 10,000 cells, indicating that they aren't much modified by a ***mitotic fraction*** until this size point. The exponential lines of pole cells, and disc cells, point back to ***Cellular Allometric Births in Time***, $B_T$, close to the first moments of fly life (FIGURE A21), about which we shall have more to say below.

The exponential lines of growth for the histoblast nests and the spiracular anlage point back to ***Cellular Allometric Births in Time***, $B_T$, to between 100 and 125 hours of development (FIGURE A20). Zalokar's and Erk's data[108] and Rabinowitz's,[120] on pole cells numbers show a remarkable log-log linearity back to 2 cells (**$r^2$=0.91,** FIGURE A21).

There are dramatic differences in the slopes of these exponential lines, indicating that these body parts have a variety of cell-heritable ***cell cycle times***, $c_p$, which account for changes in the relative size of these parts that occur as the fly matures (FIGURES A20 and A21). These differences in slope lead to dramatic changes in the relative sizes of each of these body parts.

Does the data on this graph tell us that discs arise from single ***Founder Cells***? We regard the best way to treat these graphs is to consider them as intriguing findings and useful hints for where we should look, by high resolution 4D microscopy, for how these parts might arise,[22,23,93,153,153] and we shall return to this question again below.

The remarkable log-linearity of growth that is seen below 10,000 cells is followed by a decline in this log-linearity, as can be seen more carefully when we display just a single imaginal disc (FIGURE A21). The reduction in the proliferative potential of discs and histoblast nests above 10,000 cells in size would allow their growth at these larger sizes to be amenable to the ***Mitotic Fraction Method***, another possibility worthy of experimental and computational examination.

Several structures of the developing drosophila have been traced from as few as 2 cells.

Note in FIGURE A21 how the Spiracular Analgen and Pole cells have been tracked to just 2 cells, and that the exponential growth of these populations points directly to those two cells, which project back beyond in time to a point where these populations could possibly have arisen from just a single cell. Of course, the numbers shown in FIGURES A20 and A21 were collected at separate moments in time, but, as we noted above, Verma and Cohen have carried out continuous microscopic observations of the development of histoblast nests, in which they observed each nest to arise as the progeny of the first cells in their videos, which they captured starting from as few as 4 cells[119]. Clearly, a useful goal would be to use such a method, beginning slightly earlier in time, to determine if these, and other structures, might, indeed, have arisen from single cells. We shall return again to this question below.

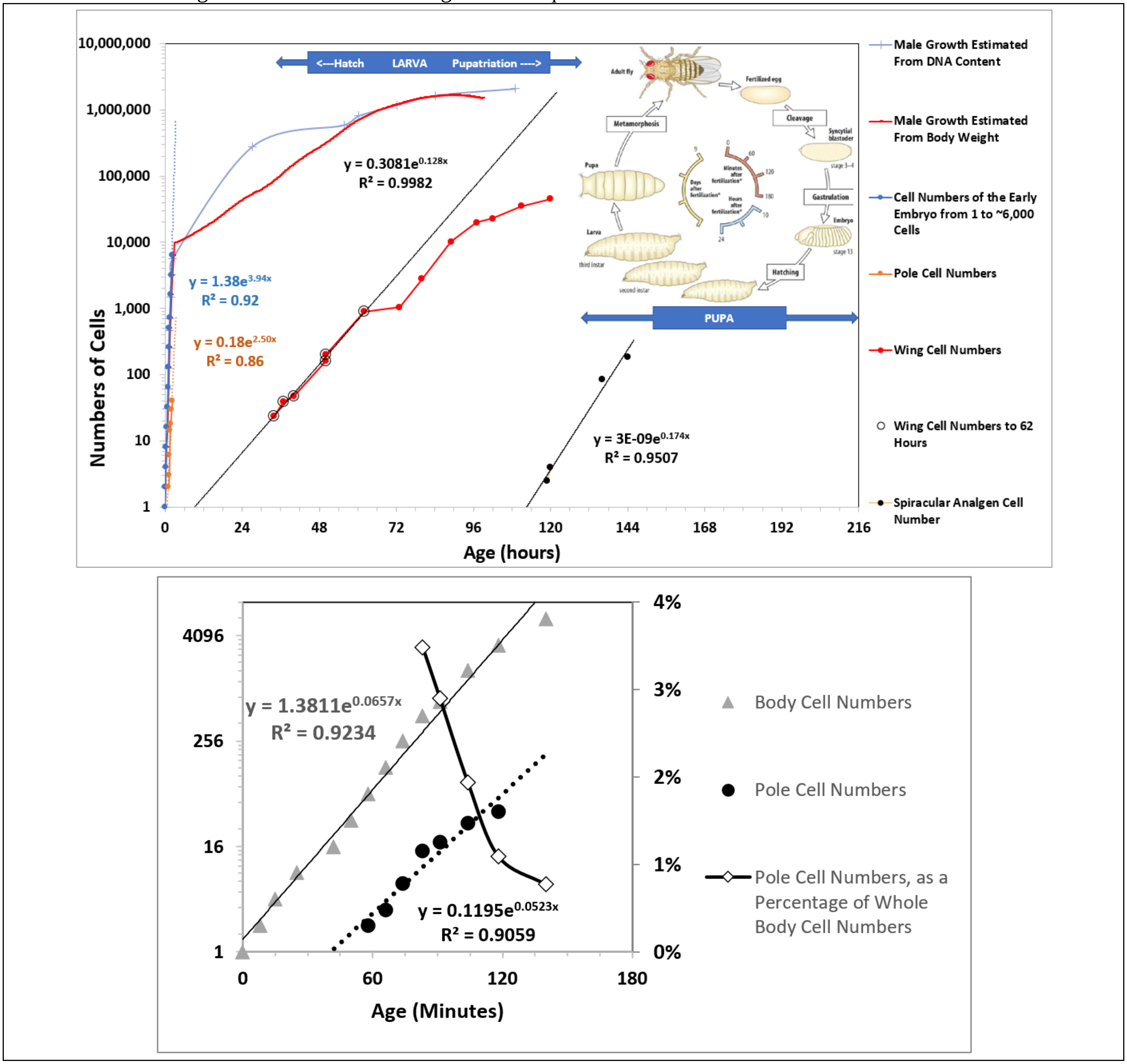


**FIGURE A21:**

**The *Cellular Phylodynamic Analysis* of *Drosophila Growth* and *Development* showing just one imaginal disc and the Spiracular Analgen (top) and Pole cells (bottom): note how the Spiracular Analgen and Pole cells have been tracked from just 2 cells.**

Mitotic Recombination

By mitotic recombination induced by irradiation at ~3 hours (embryos irradiated before that time point die at a very high rate), Weishchaus and Gehring (1971)[121] have identified that 6 ***Ancestor Cells*** were marked for the leg disks (3 hours), while Postlethwait and Schneiderman (1971)[122] have identified that 8 ***Ancestor Cells*** were marked for the leg disks (3 hours), and Bryant (1971) [123] has identified 10 ***Ancestor Cells*** were marked for the wing disks. (Bryant also saw 11 ***Ancestor Cells*** were marked for the wing disks when marked at 2 hours, a datapoint that requires confirmation, since the standard errors were quite large, and Weishchaus and Gehring (1971)[121] and others have found high levels of embryonic and cell death by irradiation at this time).

Taken together, these findings reflect discs arising from small numbers of ***Ancestor Cells***. Are these small numbers of ***Ancestor Cells*** the sizes at which these structures are created, or could these body parts arise from even smaller numbers of precursor cells? Several studies have carried out mitotic recombination induced by irradiation at various points in time, and these data for legs, wings, and antenna discs are shown in FIGURE A22 below. As Larsen-Rapport (1986)[124] points out "if the recombination event occurs at the time of X irradiation, the genetically marked cell itself will have to divide (some 15 hr after irradiation) before it can be used for certain purposes like testing for cell lineage restriction to one or another structure" (and our ***Cellular Phylodynamic Analysis*** shows imaginal disc doubling time in this range: FIGURE A20), so we have added the 15 hours. Remarkably, when displayed on log graphs, these numbers of ***Ancestor Cells*** marked by X-radiation point back to 1 ***Founder Cell*** early in the development of the drosophila embryo, precisely where the log graphs from cell counts point (FIGURE A20).

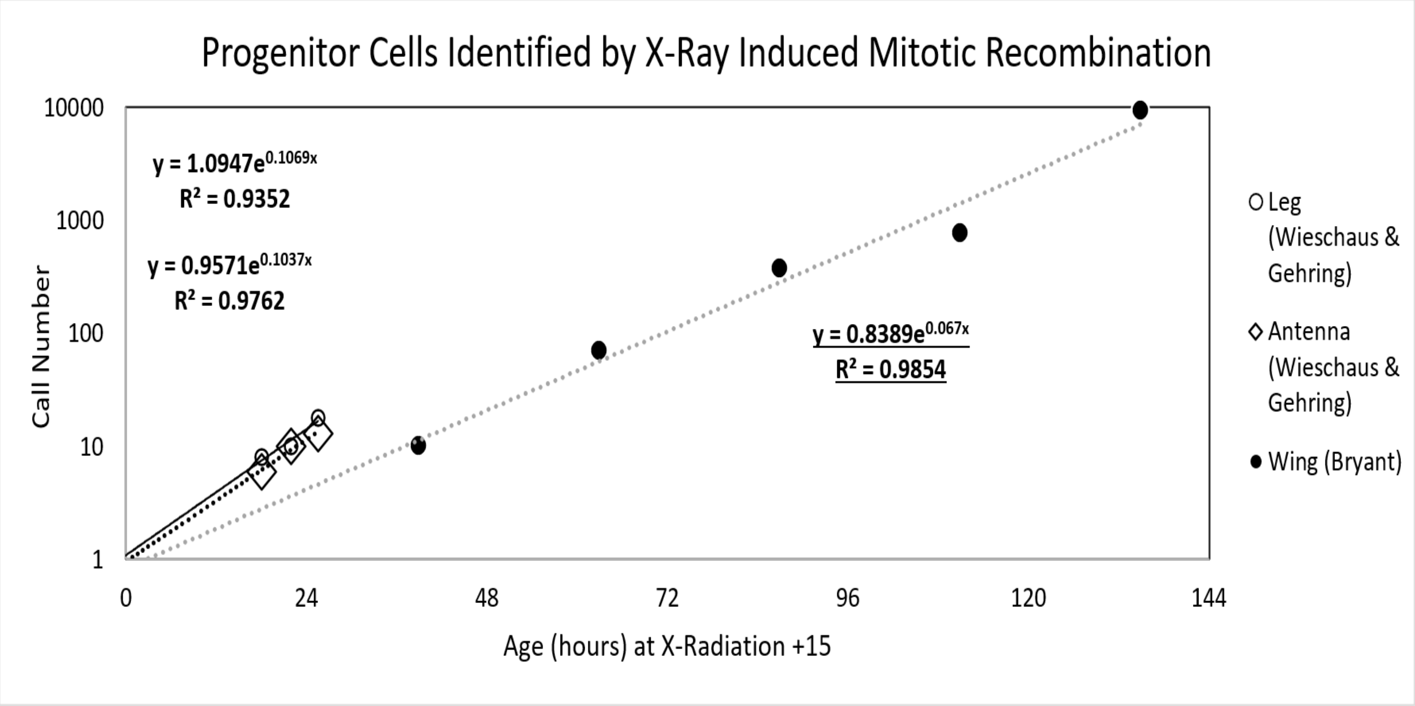


**<u>FIGURE A22</u>: The *Cellular Phylodynamic Analysis* of *Drosophila* cells marked by mitotic recombination**

Does the data on this graph again tell us that discs arise from single ***Founder Cells***? We again regard the best way to treat these graphs is to consider them as intriguing findings and useful hints for where we should look, by high resolution 4D microscopy, for how these parts might arise.[22,23,93,153,153]

Gynandromorph Marking

Using cell marking by gynandromorphs, Madhavan and Schneiderman (1977)[115] have identified that 8-9 ***Ancestor Cells*** were marked for the antenna disks, Bryant and Schneiderman (1969)[125] have identified that 20 ***Ancestor Cells*** were marked for the leg disks, Ripo1l (1972)[126] has identified that 47 ***Ancestor Cells*** were marked for wing disks, Morata and Garcia-Bellido (1976)[127] have identified that 6 ***Ancestor Cells*** were marked for haltere disks, Schupbach et al (1976) have identified that 9 ***Ancestor Cells*** that were marked for female genital discs disks and 7 ***Ancestor Cells*** for male genital discs.

These findings again reflect discs arising from small numbers of ***Ancestor Cells***. Unfortunately, these numbers are not precise, ***first*** because their calculation assumes that all parts of the body grow at the same rate, while the ***Cellular Phylodynamic Analysis*** of the growth and development of the drosophila body and its parts shown in FIGURES A20-A22 has revealed that they have dramatically different ***Cell Cycle Times*** and rates of growth; and, ***second***, because gynandromorph ***Ancestor Cell*** number calculations assume that the genetic change always takes place at the same time in the same cells. Indeed, details in the literature make clear that these changes occur over a range of time during early development. Clearly, authors making these calculations have assumed change takes place at a single point in time. For example, Garcia-Bellido and Meriman (1969)[128] suggest that "Gynandromorphs are generally thought to result from nondisjunction or chromosome elimination at the first zygotic division." and Hotta and Benzer (1972)[129] note that "The method of generating Drosophila mosaics used in this paper depends on loss of an unstable, ring-X chromosome … during the first division of the zygote nucleus of a female embryo". However, a number of studies have found that such change does not take place at a single point in time. For example, Zalokar et al (1980)[130] reports that such losses are common in the first division, but also occur in subsequent divisions. As he notes "The proportion of male nuclei ('by loss of the ring-X chromosome') varied from 80.8 to 0.4%, indicating that there must have been more than one loss of the ring-X in most of the eggs and that losses occurred as late as the ninth division". Portin's analysis (1978)[131] of the phenotype of gynandromorphs seems to support this, as he states "two-thirds of the gynandromorphs arise as a consequence of first division loss, whereas one third arise as a consequence of X chromosome loss in a later division." Of course, these concerns are relevant for an earlier era, and we now have high resolution 4D microscopy,[22,23,93,153, 153] which can define the originating events with precision. The results of the ***Cellular Phylodynamic Analysis*** we have presented here again point us to where we should look.

Body Part Size to Body Part Size Relationships; ***Body Part Proportion Equations***

Let us now consider the growth of two parts of the body, $p_1$ and $p_2$, both of which display allometric growth with reference to the growth of the body as a whole, $w$.

For part 1:

$$\log(N_w) = \left(\frac{S_{N1}}{c}\right)\log(N_{p1}) + \log(B_{N1}) \tag{719}$$

For part 2:

$$\log(N_w) = \left(\frac{S_{N2}}{c}\right)\log(N_{p2}) + \log(B_{N2}) \tag{720}$$

It follows that:

$$\log(N_w) = (S_{N1})\log(N_{p1}) + \log(B_{N1}) = (S_{N2})\log(N_{p2}) + \log(B_{N2}) \tag{721}$$

which is equivalent to:

$$\log(N_{p1}) = \left(\frac{S_{N2}}{S_{N1}}\right)\log(N_{p2}) + \log\left(\frac{B_{N2}}{B_{N1}}\right) \tag{721b}$$

In other words, if two parts of the body show log-linear allometric growth with respect to the size of the body as a whole, they will also show log-linear allometric growth with respect to each other, with each part's ***Cellular Allometric Birth***, $B_N$ and ***Cellular Allometric Slope***, $S_N$ repackaged into the slope and intercept of this new variation on the ***Cellular Allometric Growth Equation***.

It follows that if the body parts in question are in units of mass or volume or length, we can repeat the treatment outlined in the previous section. Thus:

$$\log(p_1) = \left(\frac{1}{S_{1,2}}\right)\log(p_2) + \log(B_{1,2}) \tag{722}$$

which is equivalent to:

$$p_1 = B_{1,2} \cdot p_2^{\frac{1}{S_{1,2}}} \tag{722b}$$

We call these expressions ***Body Part Proportion Equations***.

#### *Clones Within Clones*

As can be seen in Equation #722 above, if each of two parts of the body are seen by ***Cellular Allometric Growth Analysis*** to conform to the ***Cellular Allometric Growth Equation*** (#7), when each is compared with the body as a whole, they will also conform to the ***Cellular Allometric Growth Equation*** when compared with each other. An actual example of this can be seen in FIGURE A23, in which we display the number of cells in the anterior part of the drosophila wing, $N_{p\text{-}a}$, in comparison to the number of cells in the wing as a whole, $N_{p\text{-}w}$. These values were generated from measurements of the area of the whole wing disc, and of its anterior part, that have been collected by Parker and Shingleton,[132] translated into likely cell numbers, $N_{p\text{-}a}$, and $N_{p\text{-}w}$, with cell number per area data derived by Ulrike.[133]

These log-log comparisons of $N_{p\text{-}a}$ (number of cells in the anterior part of the drosophila wing), and $N_{p\text{-}w}$ (number of cells in the wing as a whole), are shown in FIGURE A23. This **log**($N_{p\text{-}a}$) vs **log**($N_{p\text{-}w}$) comparison reveals that the ***Cellular Allometric Growth Equation*** captures the growth of the anterior part of the wing, pointing to a ***Cellular Allometric Birth***, $B_{N1}$, growing from when the wing as a whole was 8 cells in size ($B_{N1}=2^3=8$), should the anterior part have arisen from a single ***Founder Cell***. Furthermore, the ***Cellular Allometric Slope***, $S_{Np}$, is greater than **1**, indicating an increase in speed of the ***cell cycle time***, $c_p$, of the cells in the anterior part of the wing, thus accounting for the progressively increasing relative size of the anterior part of the wing in relationship to the wing as a whole (FIGURE A24). We call this process "***Clones Within Clones***", a mechanism which provides animals a way to create functional modularity.

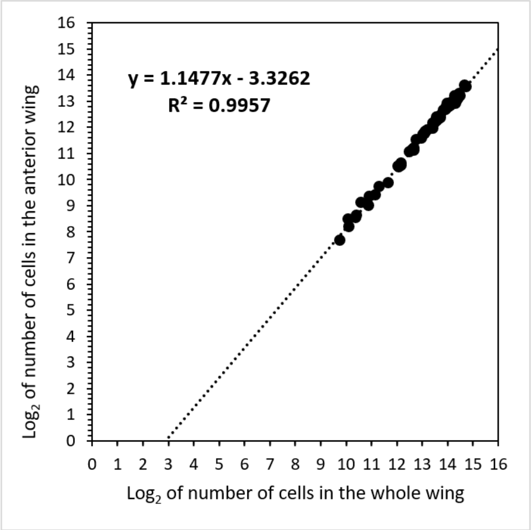


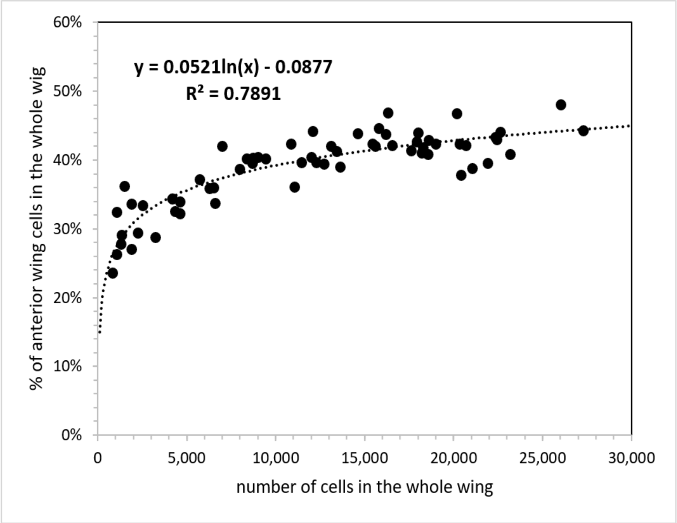


**FIGURE A23: Left. *Cellular Allometric Growth Analysis* of the anteriors wing vs whole wing.**

**FIGURE A24: Right. Percentage of anteriors wing cells in the whole wing vs number of cells in the whole wing.**

These observations are also strikingly complementary to the long appreciated finding the anterior and posterior parts of the fly's wing are clonally separate entities, called ***compartments***.[111]

Whether the anterior part of the wing grew from a single ***Founder Cell***, at the ***Cellular Allometric Birth***, $B_{N1}$, or from more than **1** ***Founder Cell***, after the ***Cellular Allometric Birth***, $B_{N1}$, cannot be known from the data we have, since the values for $N_{p\text{-}a}$, and $N_{p\text{-}w}$ only point down to that origin at when the wing as a whole was 8 cells in size ($B_{N1}=2^3=8$). Nonetheless, this poses an eminently answerable question, which could be settled by light sheet microscopy, again providing motivation for experiment.[22,23,93,153,153]

The formation of ***Clones Within Clones*** occurs in many instances of embryonic development of many animals. For example, the melanocytes of the skin of mice appear to be derived from 34 ***Founder-Cells*** (17 on each side of the body) created around the 12th day of life.[134] We also see ***Clones Within Clones*** at work in the immune system, where ***clonal selection*** selects individual cells and expand them up to large populations of cells through the action of antigens stimulating cell division by attaching to antibodies or T-cell receptors at the cell surface.[98,99,135] Similar processes may well occur in other systems, such as in the production of plasma proteins by the liver, in which each of the hundred-or-so blood proteins produced by the liver appears to be made by separate cells, in separate clones, that arise as patches continuously throughout life.[83,99] Anatomically, the creation of such functional diversity by the creation of ***Clones Within Clones*** has been seen time after time, such as in the clones that form the crypts of the intestine,[136] and in the sequential clones of spermatocytes that produce the spermatozoa,[137] as well as the many anatomical structures that seem to be made of countless subunits, such as in the lungs, the liver, the kidney, and so on. ***Cellular Phylodynamic Analysis*** seems poised to make sense of how cells create such modular structures.

***ALLO-GROWTH***: Summary Definition

Let us summarize: ***ALLO-GROWTH*** is the process by which body parts are created from ***Founder Cells***. ***ALLO-GROWTH*** is captured by the ***Cellular Allometric*** and ***Allometric Growth Equations***, and their parameters, ***S***, ***B***, $S_N$, the ***Cellular Allometric Slope***, and $B_{N1}$, the ***Cellular Allometric Birth***. ***ALLO-GROWTH*** occurs by a ***Founder Cell*** acquiring a ***Cell-Heritable***, ***Cell Cycle Time***, $c_p$ at the ***Founder Cell's Cellular Allometric Birth***, $B_{N1}$, and then undergoing mitotic expansion, as captured by the ***Cellular Allometric Slope***, $S_N$, of the ***Cellular Allometric Growth Equation***. Whether a body part arises from a single ***Founder Cell***, or multiple ***Founder Cells*** arising at the same time with the same ***Cell Cycle Time***, growth by the ***Cellular Allometric*** and ***Allometric Growth Equations*** will occur. For body parts arising from a single ***Founder Cell***, the ***Cellular Allometric Birth***, $B_{N1}$, corresponds to the number of cells in the body when the ***Founder Cell*** arose.

The ***E Unim Pluribus*** Hypothesis; Perhaps Embryos Make Their Body Parts from Single ***Founder Cells***

In 1973, Garcia-Bellido, Ripoll, and Morata[138] reported that there were boundaries within the fly wing, across which the progeny of individual cells do not travel, thus defining the structures which they called "compartments".[139] Building on these findings, Crick and Lawrence[111] "propose[d] to call the cells in a compartment a 'polyclone'" and went on to suggest "the other side of the idea must also be mentioned: a compartment is never a clone, except perhaps accidentally in rare cases". Others, such as Held,[140] proceeded to assert that drosophila imaginal "Discs are not clones". This polyclone hypothesis has slowly hardened into common knowledge, curiously without experimental verification. Indeed, in my years-long search of the literature, I have yet to find the example of any body part, in drosophila, or in another animal, in which more than **1** unrelated ***Founder Cell***, goes on to make a body part. Furthermore, as we have seen here, in every case for which we have full ***cell lineage charts***, which describe every cell in the embryo from the zygote onward (FIGURE A8), that is, for nematode worms ***Caenorhabditis elegans***,[20] and ***Meloidogyne incognita***,[21] and for tunicates, ***Oikopleura dioica***,[88] which are chordates quite closely related to ourselves, body parts trace back to single ***Founder Cells***. We have also seen here how cell marking experiments have not actually yielded the unambiguous results one might have hoped for in support of the polyclone concept. In addition, as we have outlined in the many cases described above, the ***Cellular Phylodynamic Analysis*** of body part growth and development has repeatedly provided tempting hints that the way embryos make body parts is to make them out of individual ***Founder Cells***.

To address this dilemma, I (jsm) would suggest the "***E Unim Pluribus***" hypothesis (***From One***, ***Many***). This ***E Unim Pluribus*** hypothesis proposes that developing animals create ***cell lineages***, ***tissues***, ***organs***, and ***anatomical structures*** out of undifferentiated populations of cells, by the ***Cellular Selection***[98,99] of single ***Founder Cells***, which have undergone ***Cell Heritable*** changes in phenotype, and which are then expanded, by mitotic ***ALLO-GROWTH***, into to the functional body parts that make multicellular animal life possible. The ***E Unim Pluribus*** hypothesis embraces the possibility that it is from single founder cells that embryos make parts. The ***Clones Within Clones*** phenomenon embraces the possibility that ***E Unim Pluribus*** events may occur at the initiation of body parts, and again, and again, within body parts, thus providing for a way to make modular body parts, such as the intestine, with its multiple crypts,[141] or the liver, with its multiple clones making various plasma proteins.[99] This ***E Unim Pluribus*** hypothesis is now testable, as it was not 40 years ago, with high resolution 4D microscopy.[22,23,93,153,153] All we have to do is go and take a look and count.

***CELLULAR POPULATION TREE VISUALIZATION SIMULATION***

The reader will note that we have extracted information on the relationship the number of cells in the body as a whole, $N_w$, with age, $t$, from the cell lineage charts of nematodes (FIGURES A8 and A10). However, even without full lineage chart information, we have been able to reconstruct a plausible approximation of the cell lineage of an animal by simulation with the $a$, $b$, $c$, $S_N$, $B_{N1}$, parameters of the ***Universal Growth*** and ***Cellular Allometric Growth Equations*** extracted by ***Cellular Allometric Batch Growth Analysis*** of aggregate growth that has been measured in animals for which cell lineages have not been characterized. Stadler, Pybus, and Stumpf[1] have defined four types of cell lineage charts, the first of which is, "a population tree represents the whole process by which an ancestral cell yields a population of descendent cells through division and death", and in the spirit of their approach, we call the results of such as method a "***Cellular Population Tree Visualization Simulation***". We created this ***Cellular Population Tree Visualization Simulation*** method for generating likely cell lineages of the body and its parts in JAVA (code at the end of this APPENDIX), examples of which can be seen in FIGURE 25.

In FIGURE A25 (left) we display a ***Cellular Population Tree Visualization Simulation*** of the idealized case of the cell lineage chart of an animal growing by the ***Universal Growth Equation***. Note how, as time goes on, an increasing fraction of cells, the ***Mitotic Fraction, m,*** don't divide, and some of the branches of the tree become extended in the absence of mitosis.

The ***Universal Growth*** and ***Cellular Allometric Growth Equations*** can also be taken to together, with their various parameters, to reconstruct an even more likely chart of an animal's estimated cell lineage. In FIGURE A25 (right) we display the idealized case of the ***Cell Heritable*** shortening of the ***Cell Cycle Time*** that occurs in one ***Founder Cell*** when the embryo is 8 cells in size (***Cellular Allometric Birth***, $B_{N1}$=**8**), leading to the progeny of that ***Founder Cell*** increasing in number in comparison to the rest of the embryo.

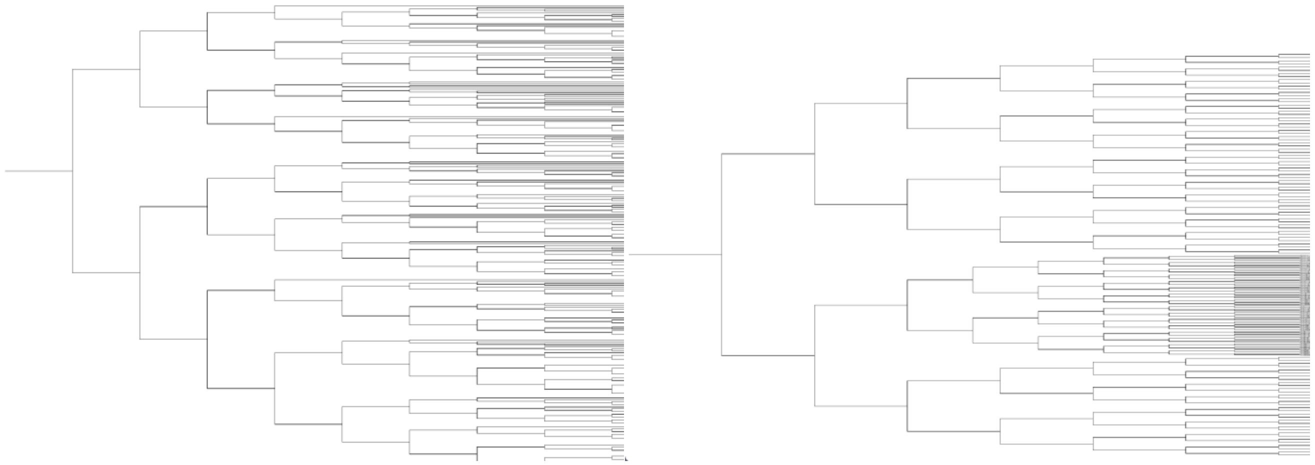

**FIGURE A25**

Cellular ***Population Tree Visualization Simulations***.

Left, the idealized case of the cell lineage chart of an animal growing by the ***Universal Growth Equation***. Note how, as time goes on, an increasing fraction of cells, the ***Mitotic Fraction, m,*** don't divide, and some of the branches of the tree become extended in the absence of mitosis. We have treated the ***Mitotic Fraction, m,*** as a random variable, a plausible assumption simply for not knowing which cells are dividing at each moment in time, and a possible biological process, if the ***Mitotic Fraction*** is actually found to be the result of the discrete allocation of ligand molecule among cells, as indicated by the mathematical analysis of such a process (Equations #700-#720).

Right, the idealized case of the cell heritable shortening of the ***Cell Cycle Time*** that occurs in one ***Founder Cell*** when the embryo is 8 cells in size (***Cellular Allometric Birth***, $B_{N1}$=**8**); note how the progeny of the ***Founder Cell*** increased in number in comparison to the rest of the embryo.

Code is executed in JAVA (see below). Visualizations made with IcyTree[142]. Future generations of such simulations may well benefit for the stochastics simulator[143] located within BEAST.[144]

***Cellular Population Tree Visualization Simulations*** brings the powerful mathematics of lineage formation, coalescent theory,[145-148] to developmental biology. For example, there has been much interest in the past few years in extracting information on gene expression, differentiation, and lineage formation, from single cell RNA expression, and cellular barcoding. However, as Telford[149] and Klein[150,151] have pointed out, without knowledge of the mitotic pattern of an animal's cell lineage, extraction of useful information from such single cell data is problematic. One might hope, indeed expect, that the potential of ***Cellular Population Tree Visualization Simulation***, driven by the parameters measured by ***Cellular Phylodynamic Analysis***, could unburden single cell analysis from the requirement of deciphering ancestry, and unlock the much hoped-for treasure of how gene expression is linked to cell lineage formation.

These ***Cellular Population Tree Visualization Simulations*** offer some intriguing possibilities for gaining a greater insight into how cells form themselves into animals and the operational components of animals. For example, the analysis of dendritic cell lineages has revealed that repeated experiments identify lineages that always seem to be different,[152,153] and this could well be ascribed to the randomness of cell division which the ***Mitotic Fraction*** imposes on cellular proliferation. Similarly, Klein and colleagues have noted[137] that the clones that form male germ cells show a curious distribution of sizes, and this could also reflect the randomness of cell division created by the ***Mitotic Fraction***, played out in lineage formation.

Because of the random quality of the ***Mitotic Fraction***, running ***Cellular Population Tree Visualization Simulations*** again and again will likely yield similar aggregate growth curves, but always yielding different cell lineage charts. That is to say, we are always the same, but we are never the same! Clearly, the law of large numbers can smooth out such stochastic variability, but one does wonder whether such lineage variability may sometime lead to phenotypic variability, or even disease. This is certainly a quality of cell lineage formation that is worthy of further investigation.

The reader will also note that we have placed much emphasis on the potential of high resolution 4D light sheet microscopy to extract information on the actual process of lineage formation.[22,23,93,153,153] We see the enormous potential of this technology to create cell lineage charts and carry out ***Cellular Lineage Growth Analysis*** to characterize the parameters of the ***Universal Growth*** and ***Cellular Allometric Growth Equations***. ***Cellular Population Tree Visualization Simulation*** should make it possible to run this mathematics in reverse, to see if it actually generates the cell lineages derived by the initial high resolution 4D light sheet microscopy. In this way, the analysis should allow a powerful back-and-forth process of discovery, to see if our comprehension of the mitotic formation of biological entities is on target, and thus to reach a progressively more accurate picture of development.

#### AGING

##### *Mitotic Fraction* and Age

The relationship between ***age***, ***t***, and ***Mitotic Fraction***, ***m***, can be calculated by combining Equations #1 and #3:

$$t(m) = \frac{c}{b \cdot log(2)} \cdot \quad [\mathrm{Ei}(-log(m)) \; - \; \mathrm{Ei}\,(-log(a))] \qquad (5)$$

We call this expression, the ***Universal Mitotic Fraction in Time Equation***. With it, we could generate curves for the 10 species listed above, and in TABLE II, which, when ended at the age of the longest known individual in each species, reveals lifespan to be correlated with an age when fewer than 1-in-1,000 cells are dividing (FIGURE 2 in text, FIGURES A 26 and A27). We call this region the ***Death Zone***, marking the thousand+fold decline in ***Mitotic Fraction*** from conception to death.

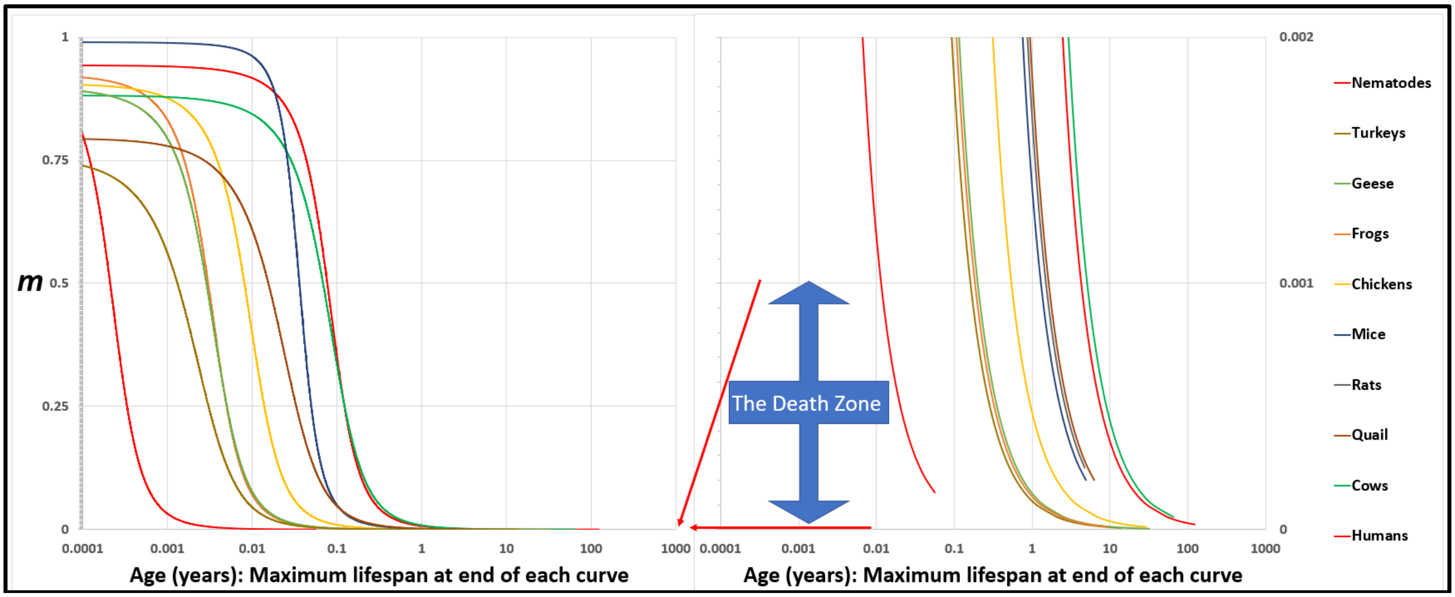


**<u>FIGURE A26</u>: The *Universal Mitotic Fraction in Time Equation* (#5) captures change in *Mitotic Fraction*, *m*, with *Age*, *t*.**

$$t(m) = \frac{c}{b \cdot log(2)} \cdot \quad [\mathrm{Ei}(-log(m)) \; - \; \mathrm{Ei}\,(-log(a))] \qquad (4)$$

The graph on the right side shows the bottom 0.2% of the Y-axis of the graph on the left side: the ***Death Zone*** occurs when fewer the 0.03% of the cells of the body are dividing. Each curve ends at the age of the oldest known member of its species.

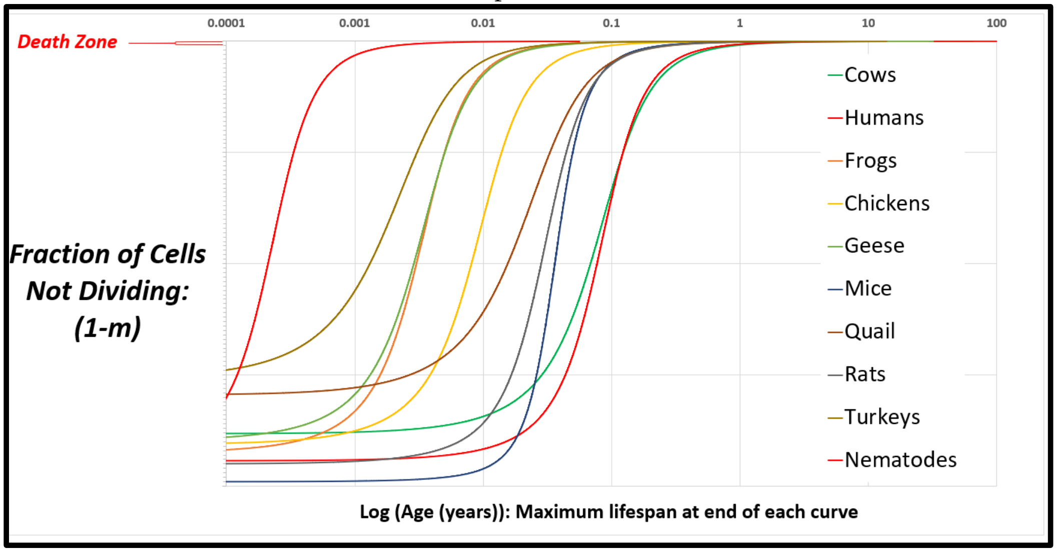


**<u>FIGURE A27</u>: The inverse of the *Universal Mitotic Fraction in Time Equation* (#5) visualizes the change in the fraction of cell not dividing with age.**

**Lifespan**

All three parameters of the ***Universal Growth Equation***, $\boldsymbol{a}$, $\boldsymbol{b}$ and $\boldsymbol{c}$, considered genetically, allow species to evolve to patterns of growth that give the greatest chances of survival, and to lifespans calculable with this expression:

$$\boldsymbol{Life\ Span} = \frac{\boldsymbol{c}}{\boldsymbol{b}\cdot \boldsymbol{log}(\boldsymbol{2})} \cdot \left[\mathrm{Ei}\big(-\boldsymbol{log}(\boldsymbol{Z})\big) - \mathrm{Ei}\big(-\boldsymbol{log}(\boldsymbol{a})\big)\right] \quad (6)$$

***Z*** is the ***Mitotic Fraction*** at the end of life. We call Equation#5 the ***Universal Lifespan Equation***, whose calculations, based on $\boldsymbol{a}$, $\boldsymbol{b}$, and $\boldsymbol{c}$ values from growth data, distinguished between the longest lifespans (humans and cows), and the shortest (nematodes), lumping in between animals with 1-to-20-year lifespans (FIGURE 3 in text, FIGURE A28). For the ten species we have examined, the average ***Z*** value of the average lifespan for each species, $\boldsymbol{Z_A}\approx 0.000037$. The average ***Z*** value of the longest known lifespans, $\boldsymbol{Z_L}\approx 0.000024$.

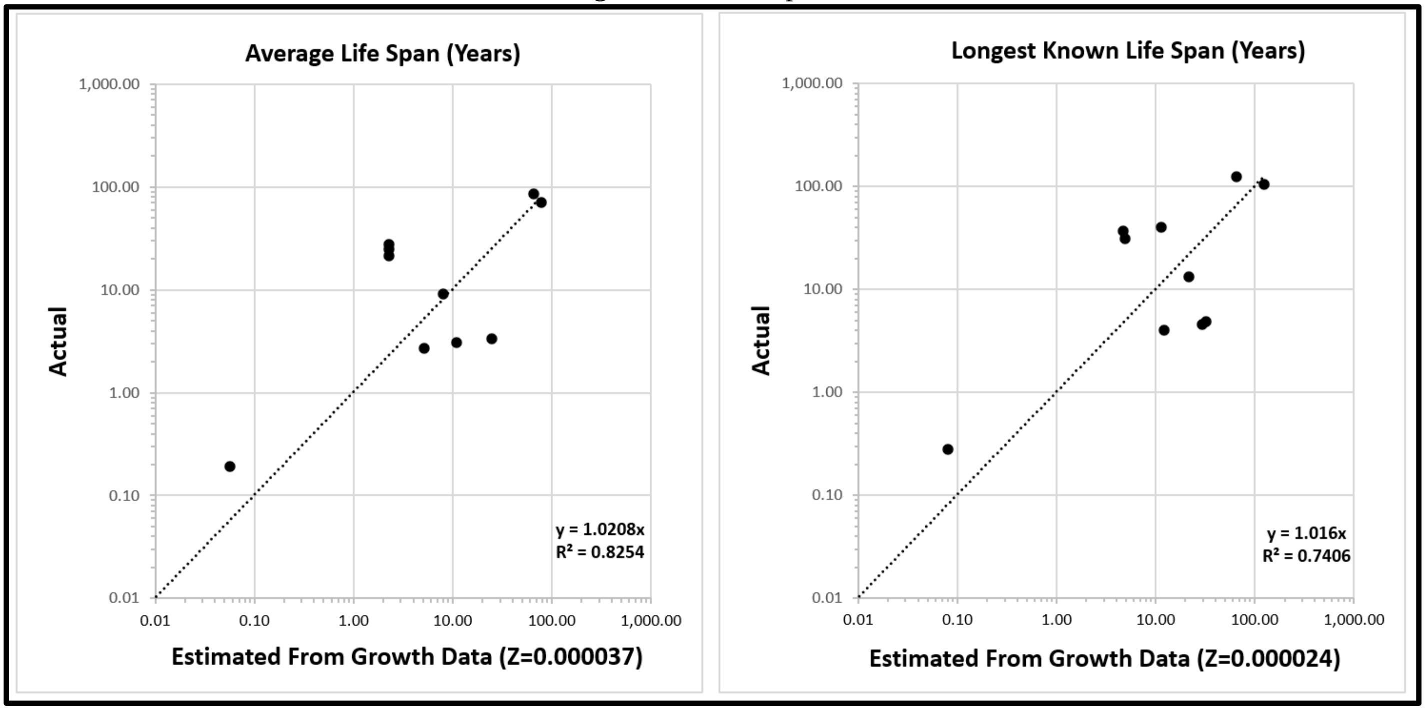


**<u>FIGURE A28</u>: Life Span - Actual vs Predicted from Growth Data** with the ***Universal Lifespan Equation*** (#6). Datapoints are for Humans, Frogs, Nematodes, Chickens, Cows, Geese, Mice, Quail, Rats, Turkeys.

#### PRESENT AND FUTURE OPPORTUNITIES FOR *CELLULAR PHYLODYNAMIC ANALYSIS*

To this point, we have seen that by examining growth and development in discrete units of integer numbers of cells, ***N***, we can gain considerable insight into multicellular organization in 2 dimensions: size and time. Let us now consider computational methods that could extend this ***Cellular Phylodynamic*** approach to examining growth and development in all 4 dimensions of space and time.

##### 4D Microscopy

There have been remarkable improvements in light sheet microscopy over the past decade, which now make possible the precise tracking body parts to their origins.[22,23,93,153,154] As might be expected, much of the use of this technology has focused on examination of the dramatic results later in the development of the parts of the body, but there is no reason why similar attention could not now be turned to the origin of these parts. As the reader has seen in this report, the ***Cellular Phylodynamic Analysis*** of animals and their parts has, over and over, pointed us to intriguing possibilities for where and when to look to see how body parts are created.

##### Cell Death

We have not addressed here the impact of cell death on morphogenesis. However, it seems clear that assembling data on the numbers of cells undergoing such death, $N_D$, offers an enticing approach to use ***Cellular Phylodynamic Analysis*** to decipher the features of this developmental process as well. For example, perhaps quantifying the numbers of cells undergoing such death, $N_D$, occurring as an ***Apoptotic Fraction***, ***D***, of the body as a whole, should provide us with an avenue for searching for the regularities of apoptosis. The occurrence of specific numbers of dying cells, $N_D$, in specific parts of the embryo, as well as in specific locations in the embryo, and at specific times in embryonic development, should also give us insight into the nature of apoptosis and its role in molding development. Such an approach to apoptosis is clearly analogues to how we carried out a ***Cellular Phylodynamic Analysis*** of the ***Mitotic Fraction***, ***m***, which led us to the ***Universal Mitotic Fraction Equation,*** and hence to the ***Universal Growth Equation,*** thus telling us how the embryo employed ***mitotic quiescence*** to regulate its growth. When combined with ***Cellular Population Tree Visualization Simulations***, such an analysis should also give us insight into the impact of cell death on the cell lineage structure of animals.

##### Manual Execution of *Cellular Lineage Growth Analysis*

The features of growth and development that were revealed by carrying out manual ***Cellular Lineage Growth Analysis*** of early nematode and tunicate embryos shown in FIGURE A9- and A10 were arrived at by sitting down with paper versions of ***cell lineage charts***, measuring the locations of each of the cells with a ruler, and counting them, both the number of cells in each ***cell lineage***, $N_p$, and the number of cells in the animal as a whole, $N_w$. This manual execution of ***Cellular Lineage Growth Analysis*** was easily done for the ***M incognita lineage chart***, which fits on a single sheet of paper (FIGURE A9) but was somewhat acrobatic for the ***C elegans*** and ***O dioica cell lineage charts***, which must be 6 feet long to discern each line (FIGURE A10 and A11). These manual executions were also exceedingly time consuming; the assembly of ***lineage*** cell number data for ***O dioica*** took four months of constant work. Furthermore, we were only able to carry out these analyses because others had already done the enormously time-consuming process of creating ***cell lineage charts*** from 4D microscope data.[22,23,93,153,154] The approach to assembling ***cell lineage charts*** for ***M incognita***, ***C elegans*** and ***O dioica***, while providing valuable information by naming ***Founder Cells*** based on the ***cell lineages*** that they produce, is also challenged as a method to assemble such data in an unbiased fashion.

##### Computational Execution of *Cellular Lineage Growth Analysis*: The *BinaryCellName* Method

Fortunately, our manual executions of the ***Cellular Phylodynamic Analysis*** of the ***cell lineages*** of suggest a way to accomplish this task computationally, in an unbiased fashion, and with data taken directly from 4D microscope images of developing embryos[22,23,93,153,154] The key to this approach is to assign each cell a binary number tag, the ***BinaryCellName***, beginning with the zygote having ***BinaryCellName*** "1", and then giving each cell a ***BinaryCellName*** that is "**1 +** ***BinaryCellName***" of the cell's parent (FIGURE A9). Thus, ***BinaryCellName*** "111" would be the child of "11", the sibling of "110", and the parent of "1111" and "1110". With this method, the forbears of each cell, as well as its place in the ***lineage*** hierarchy of the embryo, are obvious from its ***BinaryCellName***. From this information, ***Cellular Lineage Growth Analysis*** could be carried out computationally.

Comprehending Development in 4 Dimensions at the Microscopic Scale with the ***BinaryCellName*** Method

The 3-dimensional shape of each part of the embryo is affected, and possibly determined, by specifying the planes of division at each mitosis. We can see this in a number of instances, such as in the formation of the epidermis by the **AB *cell lineage*** of the ***C elegans*** nematode, as each cell division occurs perpendicular to the surface, causing the epidermis to remain one cell thick. At about the same time, the **E *lineage*** forms the intestine from the **E *Founder Cell*** by a neat series of cell divisions that occur front/back, then left/right, then front/back, then (mostly) front/back, thus creating a structure made of two, side-by-side, front-to-back, rows of cells, with each row being 7 cells long, with a couple of other cells cleverly shunted off to the bottom.[155] The ***BinaryCellName Method*** provides a way to characterize such shaping of the embryo computationally with data taken directly from 4D microscope images of developing embryos.[22,23,93,153,154]

Once assigned, each ***BinaryCellName*** can keep track of a lot of information: the cell's traditional name, the time it arose by the division of its parent, the time it disappeared by cell division or death, its phenotypic markers such as expressed genes, ***Cell-Heritable*** markers, its location in space, characterized by its centroid and boundary and other objects such as its nucleus, in units of ***x***, ***y***, ***z***, and ***t***, as well as the location of the cell's plane of division, the features of that mitotic plane of division in units of ***yaw***, ***pitch***, ***roll***, and ***t***, and the ratio of the volumes on each side of that plane. The ***Cellular Phylodynamic Analysis*** of the ***cell lineages*** could then be carried out computationally, including the assessments of log-linearity and slope for every ***lineage***, the extraction of values of the growth of each ***lineage*** in terms of the parameters of the ***Cellular Allometric Growth Equation***, especially the value of each ***lineage***'s ***Cellular Allometric Birth***, $B_N$, and ***Cellular Allometric Slope***, $S_N$, with the three-dimensional visualization of each ***lineage*** in the point-cloud that makes up the cells of the embryo as a whole. This potential of the ***BinaryCellName Method*** to examine the events of cell division and cell geometry, testing for the presence of ***allometric log linearity*** in each ***lineage***, together with the changes in gene expression that occur at each cell division, with software that could be linked directly to the microscope that collects such data, offers a way to decipher how cellular proliferation at the beginning of development drives the creation of the 3D structure of anatomy in the 4$^{th}$ dimension of time.[22,23,93,153,154]

Comprehending Development in 4 Dimensions at the Macroscopic Scale with the $S_{N3}$ ***Method***

Our ***Cellular Phylodynamic*** examination of the growth of the parts of the body has focused on embryonic structures as single dimensional masses, comprised of specific numbers of cells, $N_p$, whose relative growth we measured by the ***Cellular Allometric Slope***, $S_N$, of the ***Cellular Allometric Growth Equation***. However, there is no reason why $S_N$ couldn't also be treated as a three-dimensional array, so as to be able to infer how the changes in the locations of cells underlie the formation of the shape of each body part, an approach we call the "$S_{N3}$ ***method***".

JAVA Code and Flow Charts for ***Cellular Population Tree Visualization Simulation***

Flow chart and code for ***Cellular Phylodynamic Lineage Simulation***

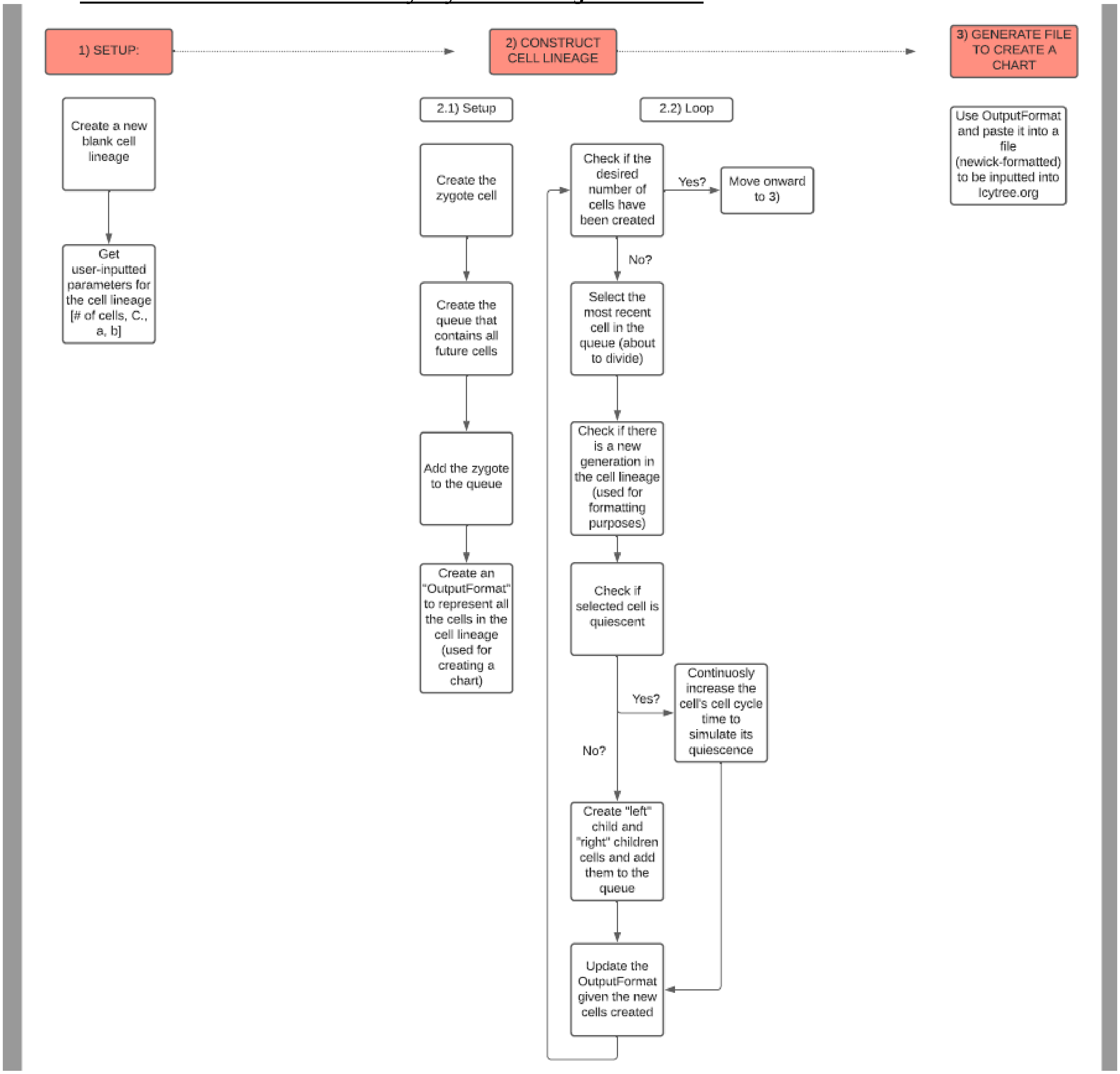


```
// Daniel Lee
// 8-4-21
// Cellular Phylodynamic Lineage Simulation

// Packages used for this simulation
import java.util.*;
import java.io.*;

/*
The "Cell" class defines the characteristics of each cell
*/
class Cell {
   // Characteristics:
```

```
    int BinaryCellName; // The binary-cell-name; to understand what this is, see the paper.
    int generation; /* The generation of the cell
                    THE ZYGOTE EXISTS IN GENERATION=1,
                    THE NEXT TWO CELLS EXISTS IN GENERATION=2,
                    ETC */
    double CellCycleTime; // Cell cycle time, C., for this cell
    Cell left, right; // This defines the two cells (children) that arise from a cell division.
    boolean IsQuiescent; // State of being quiescent or not

    // CREATION OF A NEW CELL ON THE CELL LINEAGE
    public Cell (int BinaryCellName, int generation, double CellCycleTime) {
        this.BinaryCellName = BinaryCellName;
        this.generation = generation;
        this.CellCycleTime = CellCycleTime;
        left = right = null;
    }
}

/*
The "WholeAnimalCellLineage" class used to store away information on the cell lineage
*/
class WholeAnimalCellLineage {
    Cell zygote; // The zygote cell
    int TotalNumberOfCells; // Total number of cells to be simulated
    int NumberOfAliveCells = 1; // Number of alive cells at any given moment
    int generation = 1; // Latest generation in the cell lineage
    double PresetCellCycleTime; // The uniform cell cycle time for all cells
    /* Used to calculate the mitotic fraction */
    double a;
    double b;
    StringBuffer OutputFormat; // Contains the newick-formatted representation of the cell lineage
                        // This contains all the necessary information about each cell in the cell lineage
                        // to create a nice figure using Icytree (https://icytree.org/)
                        // It consists of one long string with numbers, parenthesis, commas, and colons.

    // CREATION OF A NEW CELL LINEAGE
    public WholeAnimalCellLineage() throws IOException {
        getInfo(); // Retrieve user-inputted parameters
    }

    /*
    Generate the cell lineage based on given parameters
    */
    public void generate() {
        zygote = new Cell(0b1, 1, PresetCellCycleTime); // Creation of the zygote

        Queue<Cell> queue = new LinkedList<>(); // Contains a list (in order) of all cells that will divide
                                    // starts with the zygote
        queue.add(zygote);

        // Create the output format (with just the zygote) and update it as more cells are created
        OutputFormat = new StringBuffer(zygote.BinaryCellName + ":" + String.format("%.4f", zygote.CellCycleTime));
        int NewickIndex = -1; // Index to add future cells

        // Run the simulation until the desired number of cells is reached
        while (NumberOfAliveCells < TotalNumberOfCells) {
            Cell cell = queue.remove(); // Select the cell that may divide

            // Calculate (using the binary-cell-name) if there is a new generation in the cell lineage
```

```
        if (cell.BinaryCellName == 0b1 || (int) (Math.ceil((Math.log(cell.BinaryCellName) / Math.log(2)))) == (int)
(Math.floor(((Math.log(cell.BinaryCellName) / Math.log(2)))))) {
            // Increment the latest generation in the cell lineage
            generation++;

            // Reset the newick index
            NewickIndex = 0;
            while (OutputFormat.charAt(NewickIndex) == '(') {
                NewickIndex++;
            }
        }

        /*
        ------ CELL DIVISION + CREATION OF THE NEW CELLS ------
        */
        // CHECK THE SELECTED CELL'S QUIESCENCE
        if (cell.IsQuiescent) {

            /*
            Continuously increase the cell cycle time of the quiescent cell each iteration
            to simulate its inability to divide.
            */
            queue.add(cell);
            cell.CellCycleTime += PresetCellCycleTime;

            // Update OutputFormat to increase the cell cycle time
            int temporaryNewickIndex = NewickIndex; // temporaryNewickIndex is the left index of the cell cycle time
                                        // in OutputFormat.
            while (OutputFormat.charAt(temporaryNewickIndex) != ':') {
                temporaryNewickIndex++;
            }
            temporaryNewickIndex++;

            int temporaryNewickIndex2 = temporaryNewickIndex; // temporaryNewickIndex is the right index of the cell cycle time
                                                // in OutputFormat.

            while (OutputFormat.charAt(temporaryNewickIndex2) != ',' && OutputFormat.charAt(temporaryNewickIndex2) != ')') {
                temporaryNewickIndex2++;
            }

            OutputFormat.replace(temporaryNewickIndex, temporaryNewickIndex2, String.format("%.4f", cell.CellCycleTime)); //
Update the OutputFormat
            NewickIndex = temporaryNewickIndex2; // Update the NewickIndex
        } else {
            // CELL IS NOT QUIESCENT AND WILL DIVIDE

            // CREATION OF "LEFT" CHILD
            cell.left = new Cell(cell.BinaryCellName * 0b10, generation, cell.CellCycleTime);
            double mitoticFraction = Math.pow(a, Math.pow(NumberOfAliveCells, b)); // Calculate mitotic fraction;
                                                            // Probability the cell can divide
            if (Math.random() > mitoticFraction) {
                cell.left.IsQuiescent = true;
            }
            queue.add(cell.left);

            // CREATION OF "RIGHT" CHILD
            cell.right = new Cell(cell.BinaryCellName * 0b10 + 0b1, generation, cell.CellCycleTime);
            mitoticFraction = Math.pow(a, Math.pow(NumberOfAliveCells, b)); // Calculate mitotic fraction;
                                                        // Probability the cell can divide
            if (Math.random() > mitoticFraction) {
```

```
                cell.right.IsQuiescent = true;
              }
              queue.add(cell.right);

              NumberOfAliveCells++; // Calculate net increase of alive cells (1 splits into 2)

              /*
              ------ UPDATE THE OUTPUT FORMAT ------
              */
              OutputFormat.insert(NewickIndex, "(" + Integer.toBinaryString(cell.left.BinaryCellName) + ":" + String.format("%.4f",
cell.left.CellCycleTime) + "," + Integer.toBinaryString(cell.right.BinaryCellName) + ":" + String.format("%.4f",
cell.right.CellCycleTime) + ")");
            }

            // Update NewickIndex for future cells
            while (OutputFormat.charAt(NewickIndex) != ')' && !cell.IsQuiescent) {
              NewickIndex++;
            }
            while (NewickIndex < OutputFormat.length() && OutputFormat.charAt(NewickIndex) != ',' && generation != 1) {
              NewickIndex++;
            }
            NewickIndex++;
            while (NewickIndex < OutputFormat.length() && OutputFormat.charAt(NewickIndex) == '(') {
              NewickIndex++;
            }
        }
    }

    /*
    Used to collect user input for the cell lineage
    */
    public void getInfo() throws IOException {
        BufferedReader reader = new BufferedReader(new InputStreamReader(System.in));
        System.out.println("Enter the desired total number of cells: ");
        TotalNumberOfCells = Integer.parseInt(reader.readLine());
        System.out.println("Enter the uniform cell cycle time: ");
        PresetCellCycleTime = Double.parseDouble(reader.readLine());
        System.out.println("Enter a for the mitotic fraction: ");
        a = Double.parseDouble(reader.readLine());
        System.out.println("Enter b for the mitotic fraction: ");
        b = Double.parseDouble(reader.readLine());
        System.out.println("Great! Now submit the file named: 'WholeAnimalCellLineage.tree' to icytree.org");
    }

    /*
    Generate newick formatted file:
    https://en.wikipedia.org/wiki/Newick_format#Grammar
    */
    public void createNewickFile() throws IOException {
        try (PrintWriter fout = new PrintWriter(new BufferedWriter(new FileWriter("WholeAnimalCellLineage.tree")))) { // Create new
file to store OutputFormat
            fout.println(OutputFormat); // Store OutputFormat in the created file
        } catch (IOException e) {
            System.out.println(e);
        }
    }
}
public class CellularPhylodynamicLineageSimulation {
    public static void main(String[] args) throws IOException {
        WholeAnimalCellLineage tree = new WholeAnimalCellLineage(); // Create the cell lineage
```

```
        tree.generate(); // Generate the cell lineage
        tree.createNewickFile(); // Create the newick file named "WholeAnimalCellLineage.tree"
    }
}
```

Flow chart and code for ***Cellular Phylodynamic Lineage Simulation*** of BODY PARTS

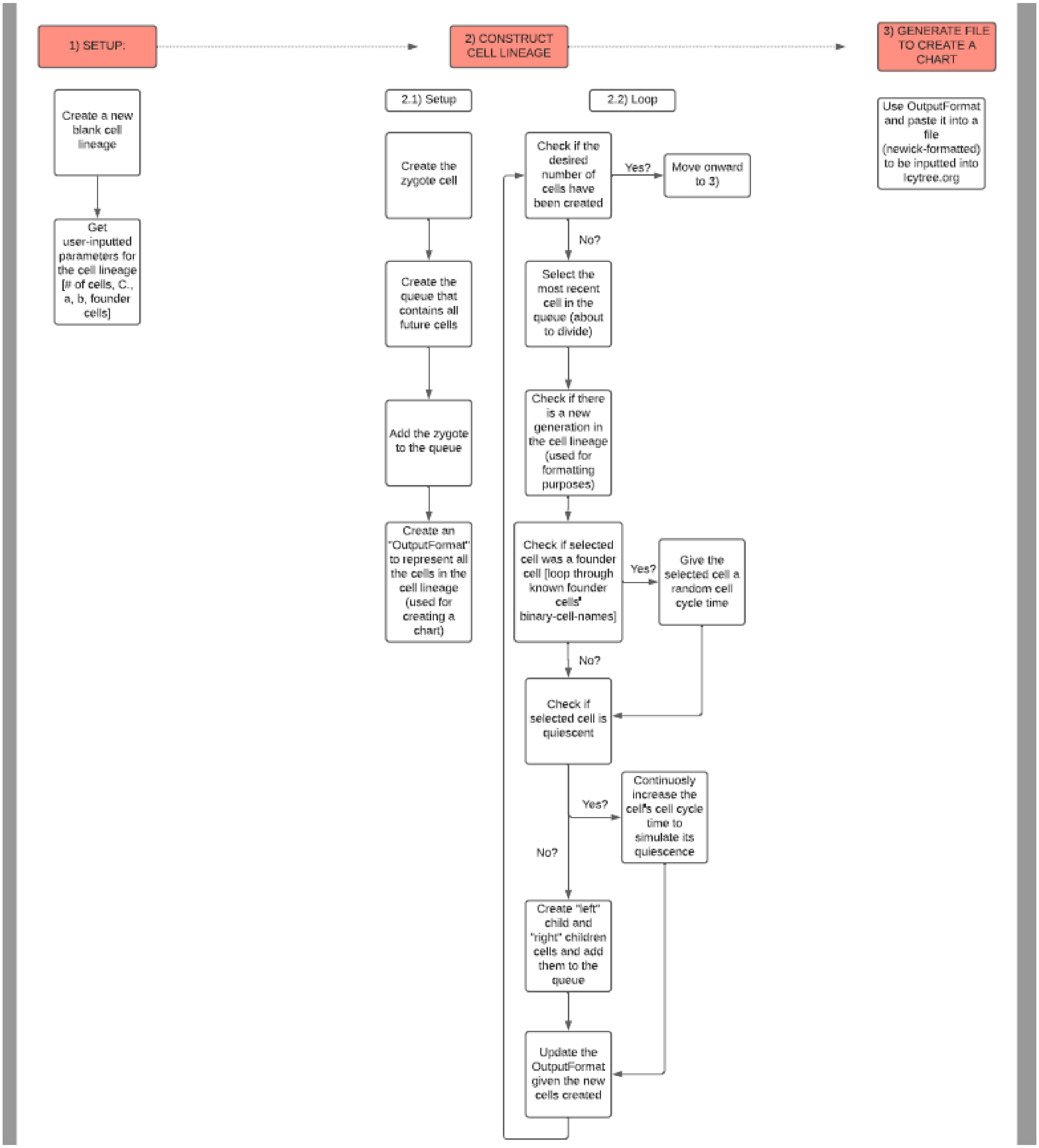

```
// Daniel Lee
// 8-4-21
// Cellular Phylodynamic Lineage Simulation of Body Parts

// Packages used for this simulation
import java.util.*;
import java.io.*;

/*
The "Cell" class defines the characteristics of each cell
*/
class Cell {
    // Characteristics:
    int BinaryCellName; // The binary-cell-name; to understand what this is, see the paper.
    int generation; /* The generation of the cell
                    THE ZYGOTE EXISTS IN GENERATION=1,
                    THE NEXT TWO CELLS EXISTS IN GENERATION=2,
                    ETC */
    double CellCycleTime; // Cell cycle time, C., for this cell
    Cell left, right; // This defines the two cells (children) that arise from a cell division.
    boolean IsQuiescent; // State of being quiescent or not

    // CREATION OF A NEW CELL ON THE CELL LINEAGE
    public Cell (int BinaryCellName, int generation, double CellCycleTime) {
        this.BinaryCellName = BinaryCellName;
        this.generation = generation;
        this.CellCycleTime = CellCycleTime;
        left = right = null;
    }
}

/*
The "WholeAnimalCellLineage" class used to store away information on the cell lineage
*/
class WholeAnimalCellLineage {
    Cell zygote; // The zygote cell
    int TotalNumberOfCells; // Total number of cells to be simulated
    int NumberOfAliveCells = 1; // Number of alive cells at any given moment
    int generation = 1; // Latest generation in the cell lineage
    double PresetCellCycleTime; // The uniform cell cycle time for all cells before the first founder cell
    ArrayList<String> FounderCells; // Contains a list of all founder cells' binary-cell-names in the cell lineage
    /* Used to calculate the mitotic fraction */
    double a;
    double b;
    StringBuffer OutputFormat; // Contains the newick-formatted representation of the cell lineage
                          // This contains all the necessary information about each cell in the cell lineage
                          // to create a nice figure using Icytree (https://icytree.org/)
                          // It consists of one long string with numbers, parenthesis, commas, and colons.

    // CREATION OF A NEW CELL LINEAGE
    public WholeAnimalCellLineage() throws IOException {
        getInfo(); // Retrieve user-inputted parameters
    }

    /*
    Generate the cell lineage based on given parameters
    */
    public void generate() {
        zygote = new Cell(0b1, 1, PresetCellCycleTime); // Creation of the zygote
```

```
    Queue<Cell> queue = new LinkedList<>(); // Contains a list (in order) of all cells that will divide
                                    // starts with the zygote
    queue.add(zygote);

    // Create the output format (with just the zygote) and update it as more cells are created
    OutputFormat = new StringBuffer(zygote.BinaryCellName + ":" + String.format("%.4f", zygote.CellCycleTime));
    int NewickIndex = -1; // Index to add future cells

    // Run the simulation until the desired number of cells is reached
    while (NumberOfAliveCells < TotalNumberOfCells) {
      Cell cell = queue.remove(); // Select the cell that may divide

      // Calculate (using the binary-cell-name) if there is a new generation in the cell lineage
      if (cell.BinaryCellName == 0b1 || (int) (Math.ceil((Math.log(cell.BinaryCellName) / Math.log(2)))) == (int)
(Math.floor(((Math.log(cell.BinaryCellName) / Math.log(2)))))) {
        // Increment the latest generation in the cell lineage
        generation++;

        // Reset the newick index
        NewickIndex = 0;
        while (OutputFormat.charAt(NewickIndex) == '(') {
          NewickIndex++;
        }
      }

      /*
      ------ CHECK IF SELECTED CELL IS A FOUNDER CELL ------
      */
      // Iterate through list of founder cells' binary-cell-names and compare to selected cell's binary-cell-name
      for (int i = FounderCells.size() - 1; i >= 0; i--) {

        // Candidate founder cell binary-cell-name
        String PotentialFounderCell = FounderCells.get(i);
        // Cell binary-cell-name
        String CellBinaryCellName = Integer.toBinaryString(cell.BinaryCellName);

        /*
        Iterate through, digit by digit, the candidate founder cell's binary-cell-name and compare it to the selected cell's
        */
        boolean isFounder = true;
        for (int j = 0; j < PotentialFounderCell.length() && isFounder; j++) {

          // Check if any digit is different or if the binary-cell-names have different lengths
          // --> Cell is not this founder cell
          if (PotentialFounderCell.length() != CellBinaryCellName.length() || PotentialFounderCell.charAt(j) !=
CellBinaryCellName.charAt(j)) {
            isFounder = false;
          }
        }

        // Found matching binary-cell-name
        if (isFounder) {
          // Set the new founder cell's cell cycle time, C., to a random number
          // (and all of its children)
          cell.CellCycleTime = PresetCellCycleTime * 5/6 + Math.random() * PresetCellCycleTime * 1/3;
          break;
        }
      }

      /*
```

```
        ------ CELL DIVISION + CREATION OF THE NEW CELLS ------
        */
        // CHECK THE SELECTED CELL'S QUIESCENCE
        if (cell.IsQuiescent) {

          /*
          Continuously increase the cell cycle time of the quiescent cell each iteration
          to simulate its inability to divide.
          */
          queue.add(cell);
          cell.CellCycleTime += cell.CellCycleTime;

          // Update OutputFormat to increase the cell cycle time
          int temporaryNewickIndex = NewickIndex; // temporaryNewickIndex is the left index of the cell cycle time
                                          // in OutputFormat.
          while (OutputFormat.charAt(temporaryNewickIndex) != ':') {
            temporaryNewickIndex++;
          }
          temporaryNewickIndex++;

          int temporaryNewickIndex2 = temporaryNewickIndex; // temporaryNewickIndex is the right index of the cell cycle time
                                                // in OutputFormat.

          while (OutputFormat.charAt(temporaryNewickIndex2) != ',' && OutputFormat.charAt(temporaryNewickIndex2) != ')') {
            temporaryNewickIndex2++;
          }

          OutputFormat.replace(temporaryNewickIndex, temporaryNewickIndex2, String.format("%.4f", cell.CellCycleTime)); //
Update the OutputFormat
          NewickIndex = temporaryNewickIndex2; // Update the NewickIndex
        } else {
          // CELL IS NOT QUIESCENT AND WILL DIVIDE

          // CREATION OF "LEFT" CHILD
          cell.left = new Cell(cell.BinaryCellName * 0b10, generation, cell.CellCycleTime);
          double mitoticFraction = Math.pow(a, Math.pow(NumberOfAliveCells, b)); // Calculate mitotic fraction;
                                                        // Probability the cell can divide
          if (Math.random() > mitoticFraction) {
            cell.left.IsQuiescent = true;
          }
          queue.add(cell.left);

          // CREATION OF "RIGHT" CHILD
          cell.right = new Cell(cell.BinaryCellName * 0b10 + 0b1, generation, cell.CellCycleTime);
          mitoticFraction = Math.pow(a, Math.pow(NumberOfAliveCells, b)); // Calculate mitotic fraction;
                                                    // Probability the cell can divide
          if (Math.random() > mitoticFraction) {
            cell.right.IsQuiescent = true;
          }
          queue.add(cell.right);

          NumberOfAliveCells++; // Calculate net increase of alive cells (1 splits into 2)

          /*
          ------ UPDATE THE OUTPUT FORMAT ------
          */
          OutputFormat.insert(NewickIndex, "(" + Integer.toBinaryString(cell.left.BinaryCellName) + ":" + String.format("%.4f",
cell.left.CellCycleTime) + "," + Integer.toBinaryString(cell.right.BinaryCellName) + ":" + String.format("%.4f",
cell.right.CellCycleTime) + ")");
        }
```

```
            // Update NewickIndex for future cells
            while (OutputFormat.charAt(NewickIndex) != ')' && !cell.IsQuiescent) {
                NewickIndex++;
            }
            while (NewickIndex < OutputFormat.length() && OutputFormat.charAt(NewickIndex) != ',' && generation != 1) {
                NewickIndex++;
            }
            NewickIndex++;
            while (NewickIndex < OutputFormat.length() && OutputFormat.charAt(NewickIndex) == '(') {
                NewickIndex++;
            }
        }
    }

    /*
    Used to collect user input for the cell lineage
    */
    public void getInfo() throws IOException {
        BufferedReader reader = new BufferedReader(new InputStreamReader(System.in));
        System.out.println("Enter the desired total number of cells: ");
        TotalNumberOfCells = Integer.parseInt(reader.readLine());
        System.out.println("Enter the uniform cell cycle time: ");
        PresetCellCycleTime = Double.parseDouble(reader.readLine());
        System.out.println("Enter a for the mitotic fraction: ");
        a = Double.parseDouble(reader.readLine());
        System.out.println("Enter b for the mitotic fraction: ");
        b = Double.parseDouble(reader.readLine());
        System.out.println("Which cells will correspond to a new body part? (Enter the binary-cell-names separated by spaces): ");
        String bodyParts = reader.readLine();
        FounderCells = new ArrayList<String>(Arrays.asList(bodyParts.split(" ")));
        System.out.println("Great! Now submit the file named: 'WholeAnimalCellLineage.tree' to icytree.org");
    }

    /*
    Generate newick formatted file:
    https://en.wikipedia.org/wiki/Newick_format#Grammar
    */
    public void createNewickFile() throws IOException {
        try (PrintWriter fout = new PrintWriter(new BufferedWriter(new FileWriter("WholeAnimalCellLineage.tree")))) { // Create new
file to store OutputFormat
            fout.println(OutputFormat); // Store OutputFormat in the created file
        } catch (IOException e) {
            System.out.println(e);
        }
    }
}
public class CellularPhylodynamicLineageSimulationBODYPARTS {
    public static void main(String[] args) throws IOException {
        WholeAnimalCellLineage tree = new WholeAnimalCellLineage(); // Create the cell lineage
        tree.generate(); // Generate the cell lineage
        tree.createNewickFile(); // Create the newick file named "WholeAnimalCellLineage.tree"
    }
}
```

**BOX 1**
**TERMS, EXPRESSIONS, AND FINDINGS FOR**
**THE *CELLULAR PHYLODYNAMICS* OF ANIMAL GROWTH**

***Cellular Phylodynamics*:**
The analysis of growth and development in units of integer numbers of cells, $N$. $N_w$ is the number of cells in an animal as a whole while $N_p$ is the number of cells in a part of the body, such as a ***cell lineage***, ***tissue***, ***organ***, ***or anatomical structure***.

The ***Mitotic Fraction, m***:
A measure of the fraction of cells dividing, derived from growth data.

***Mitotic Fraction Method***
The ***Cellular Phylodynamic Analysis*** method, Equation #3b, which gives a measure how many cells would have to divideto account for the amount of growth that occurs over each period of time, the ***mitotic fraction***.

The ***Universal Mitotic Fraction Equation***:
An empirically derived expression, $\boldsymbol{m = a^{(N_w^b)}}$, which captures a close approximation of the decline in the ***mitotic fraction***, ***m***, that occurs as animals increase in size, $N_w$, potentially traceable to the discrete allocation of growth factor molecules among cells.

The ***Universal Growth Equation***
An empirically derived expression for the rate of animal growth, $\frac{dN_w}{dt} = \left(\frac{\log 2}{c}\right) N_w a^{(N_w^b)}$. ***c*** is the ***cell cycle time***, the amount of time it takes cells to divide, traceable to how much DNA the genome contains.

The Integrated Form of the ***Universal Growth Equation***
An empirically derived expression that captures a close approximation of how animals increase in size, $N_w$, with time, $t$, such that $N_w = \int \left(\frac{\log 2}{c}\right) N_w a^{(N_w^b)}$ [Closed form: $t = c \cdot \left(\mathrm{Ei}\left[-N_w^b \cdot \log(a)\right] - \mathrm{Ei}[-\log(a)]\right) \frac{1}{b} \log(2)$].

***UNI-GROWTH*.**
The biological process captured by the ***Universal Mitotic Fraction Equation*** and the ***Universal Growth Equation***.

***Cellular Allometric Growth Analysis*.**
The ***Cellular Phylodynamic Analysis*** of relative growth carried out by examining paired $N_p$ / $N_w$ values on log-log graphs. When carried out a full lineage chart for an animal, which characterizes each cell from the first fertilized egg onward, this is called ***Cellular Allometric Lineage Growth Analysis***. When carried out without lineage data, but simply with cell numbers, this is called ***Cellular Allometric Batch Growth Analysis***.

***Cellular Allometric Lineage Growth Analysis*.**
The ***Cellular Phylodynamic Analysis*** of cell lineages, carried out by counting the number of cells in each ***lineage***, $N_p$, from the first ***Founder Cell*** onward ($N_p = 1$),

***Cellular Allometric Batch Growth Analysis*.**
The ***Cellular Phylodynamic Analysis*** of ***tissues***, ***organs***, and ***anatomical structures*** based on aggregate size, which is then converted into cell number, relying on the observation that there are about $10^8$ cells in every gram of tissue[57], or by other such batch methods, such as by making cell counts microscopically, or measuring DNA content.

The ***Cellular Allometric Growth Equation***:
An empirically derived expression that captures the relationship between the number of cells in an individual part of the body, $N_p$, with the number with the number of cells in the embryo as a whole, $N_w$, such that $\log(N_w) = \left(\frac{1}{S_N}\right)\log(N_p) + \log(B_N)$.

The ***Cellular Allometric Birth***, $B_{N1}$, of the ***Cellular Allometric Growth Equation***:
An empirically derived measure of the number of cells in the body as a whole, $N_w$, at the point where the ***cellular allometric growth equation*** indicates that a body part could have been a single cell ($N_p \approx 1$). For a cell lineage, $B_{N1}$ is the number of cells in the body as a whole, $N_w$, when the cell lineage is exactly 1 single cell ($N_p = 1$).

***Cellular Allometric Slope***, $S_N$, of the ***Cellular Allometric Growth Equation***:
An empirically derived measure of the relative growth of each part of the body, whose value can be traced to subtle ***cell-heritable*** changes in the ***cell cycle time***, potentially caused by processes such as DNA methylation.

***ALLO-GROWTH***
The biological process captured by the ***Cellular Allometric Growth Equation***.

***Cellular Population Tree Visualization Simulation***
The creation of likely approximations of cell lineage charts with the parameters of the ***Universal Growth*** and ***Cellular Allometric Growth Equations***, derived from macroscopic growth data.

#### FIGURE LEGENDS

**FIGURE A1: Growth, in units of numbers of cells, $N_w$, from fertilization until maturity, in units of days, *t*.** Datapoints for animal size's ($N_x$) versus time (***t***), for ***C elegans***, chickens, mice, turkeys, quail, geese, frogs, and humans, and their fit to the numerically integrated form of the ***Universal Growth Equation*** (#5b**).**

**FIGURE A2: Fit of human growth to the *Universal Growth Equation* (red), the *logistic equation* (blue), and the *Gompertz equation* (green).** Data values as black circles.

**FIGURE A3: Analysis of the closed form integration of the *Universal Growth Equation* (#3).**

**FIGURE A4: Decline in fraction of cells dividing, the *Mitotic Fraction, m*, calculated by the *Mitotic Fraction Method*, that occurs as animals increase in size, as seen by the *Universal Mitotic Fraction Equation*.** Data points for size, $N_w$, in integer units of numbers of cells, vs. the ***Mitotic Fraction***, ***m***, from fertilization, until maturity, for humans.

**FIGURE A5: Values of the *Mitotic Fraction*, *m*, calculated by the *Mitotic Fraction Method,* as a function of animal size, $N_w$, as shown at various scales.** Data points for animal size, $N_w$, in integer units of numbers of cells, vs. the ***Mitotic Fraction***, ***m***, from fertilization, until maturity, for humans, chickens, ***C elegans*** nematode worms, and mice.

**FIGURE A6: Comparison of the fit of *Mitotic Fraction, m,* as a function of size, $N_w$, with human growth data, to the *Universal Mitotic Fraction Equation* (green) and *West equation* (red).**

**FIGURE A7: Wave-like deviations ("*ripples*") from the *Universal Growth Equation curve,* for human growth.**

**FIGURE A8: *Cell lineage chart* of the *cell lineages* of the developing root knot nematode, *Meloidogyne incognita*.** Shown are the conventional and ***BinaryCellName Lineage*** naming methods. The **AB *lineage*** of nematodes forms the skin while the **E *lineage*** forms the intestine. ***Cell lineage chart*** and data from Calderón-Urrea et al[21]

**FIGURE A9: *Cellular Lineage Growth Analysis* of the *cell lineages* of the developing root knot nematode, *Meloidogyne incognita.***[21]

**FIGURE A10: *Cellular Lineage Growth Analysis* of the *cell lineages* of the developing *C elegans* nematode**

**FIGURE A11: *Cellular Lineage Growth Analysis* of the *cell lineages* of the chordate tunicate *Oikopleura dioica*.** Unlike ***C elegans*** and ***M incognita***, all of the cases of ***Cellular Selection*** for this species reflect reductions in cell number, although this could be due to the limited size of the dataset rather than a biological difference. ***Cell lineage chart*** data from Stach et al[88]

**FIGURE A12: *Cellular Allometric Batch Growth Analysis* of *tissues*, *organs*, and *anatomical structures* of the chick embryo.**
Growth of the parts of the chick embryonic gizzard, liver, heart, and kidney, in units of numbers of cells, in comparison to the number of cells in the embryo as a whole shown on log-log graphs, showing how the fraction of cells comprising in the gizzard and liver increases as the number of cells in the embryo increase with development and how the fraction of cells comprising in the heart decreases as the number of cells in the embryo increase with development, and how the fraction of cells comprising in the kidney remains roughly constant as the number of cells in the embryo increases with development,

**FIGURE A13**: ***Cellular Allometric Batch Growth Analysis* of tissues, organs, and anatomical structures of the rat embryo**.
Growth of body parts of the rat embryonic liver, brain, kidney and foreleg, in units of numbers of cells, in comparison to the number of cells in the embryo as a whole, are shown on log-log graphs. Note how the fraction of cells comprising in the liver kidney and forleag increases as the number of cells in the embryo increase with development, while the fraction of cells comprising the brain decreases.

**FIGURE A14**: ***Cellular Allometric Batch Growth Analysis* of tissues, organs, and anatomical structures of the mouse embryo**.
Growth of body parts of the mouse embryonic liver, brain, kidney and foreleg, in units of numbers of cells, $N_p$, in comparison to the number of cells in the embryo as a whole, $N_w$, shown on log-log graphs. Note how the fraction of cells comprising in the liver kidney and foreleg increases as the number of cells in the embryo increase with development, while the fraction of cells comprising the brain decreases.

**FIGURE A15**: ***Cellular Allometric Batch Growth Analysis* of tissues, organs, and anatomical structures of the developing human fetuses, from autopsy data, of individual fetuses**.
For additional human relative growth data, see FIGURE A16

**FIGURE A16**: ***Cellular Allometric Batch Growth Analysis* of tissues, organs, and anatomical structures of the developing human fetuses, from autopsy data, averaged values.**[156]
For additional human relative growth data, see FIGURE A15

**FIGURE A17**: ***Cellular Allometric Batch Growth Analysis* of the Zebrafish Lens.**

**FIGURE A18**: **The *Cellular Allometric Growth Equation*, $\log(N_w) = (1/S_N) \log(N_p) + \log(B_{N1})$.**

**FIGURE A19**: **Calculations for an idealized part of the body that is made from 3 *Founder Cells*, arising early in development**

**FIGURE A20**: **The *Cellular Phylodynamic Analysis* of *Drosophila Growth* and *Development***

**FIGURES A21**: **The *Cellular Phylodynamic Analysis* of *Drosophila Growth* and *Development* showing just one imaginal disc**

**FIGURE A22**: **The *Cellular Phylodynamic Analysis* of *Drosophila* cells marked by mitotic recombination**

**FIGURE A23**: ***Cellular Allometric Batch Growth Analysis* of the anteriors wing vs whole wing**

**FIGURE A24**: **Percentage of cells in the of the anteriors wing vs whole wing**

**FIGURE A25**: **Cellular *Population Tree Visualization Simulations***.
Left, the idealized case of the cell lineage chart of an animal growing by the ***Universal Growth Equation***. Note how, as time goes on, an increasing fraction of cells, the ***Mitotic Fraction, m,*** don't divide, and some of the branches of the tree become extended in the absence of mitosis. We have treated the ***Mitotic Fraction, m,*** as a random variable, a plausible assumption simply for not knowing which cells are dividing at each moment in time, and a possible biological process, if the ***Mitotic Fraction*** is actually found to be the result of the discrete allocation of ligand molecule among cells, as indicated by the mathematical analysis of such a process (Equations #700-#720).
Right, the idealized case of the cell heritable shortening of the ***Cell Cycle Time*** that occurs in one ***Founder Cell*** when the embryo is 8 cells in size (***Cellular Allometric Birth***, $B_{N1}$**=8**); note how the progeny of the ***Founder Cell*** increased in number in comparison to the rest of the embryo.
Code is executed in JAVA (see below). Visualizations made with IcyTree[142]. Future generations of such simulations may well benefit for the stochastics simulator[143] located within BEAST.[144]

**FIGURE A26: The *Universal Mitotic Fraction in Time Equation* (#5) captures change in *Mitotic Fraction*, *m*, with *Age*, *t*.**

$$\boldsymbol{t}(\boldsymbol{m}) = \frac{\boldsymbol{c}}{\boldsymbol{b} \cdot \boldsymbol{log}(\mathbf{2})} \cdot \left[\mathrm{Ei}\left(-\boldsymbol{log}(\boldsymbol{m})\right) - \mathrm{Ei}\left(-\boldsymbol{log}(\boldsymbol{a})\right)\right] \qquad (4)$$

The graph on the right side shows the bottom 0.2% of the Y-axis of the graph on the left side: the ***Death Zone*** occurs when fewer the 0.03% of the cells of the body are dividing. Each curve ends at the age of the oldest known member of its species.

**FIGURE A27: The inverse of the *Universal Mitotic Fraction in Time Equation*** (#5) **visualizes the change in the fraction of cell not dividing with age.**

**FIGURE A28: Life Span - Actual vs Predicted from Growth Data** with the ***Universal Lifespan Equation*** (#6). Datapoints are for Humans, Frogs, Nematodes, Chickens, Cows, Geese, Mice, Quail, Rats, Turkeys.

#### TABLE A1
#### FILES WITH GROWTH DATA, FOR DISTRIBUTION TO INTERESTED READERS

| Species | Genus Species | Basic Data Files | Calculations Files |
|---|---|---|---|
| Human Males | *Homo sapiens* | Boys Basic Growth Data jsm 2 12 2013.xlsx | 140330 Human males.xlsx |
| Frogs | *Rana pipiens* | Rana Basic Growth Data jsm 2 15b 2013.xlsx | 140330 Rana Pipiens.xlsx |
| Nematodes | *C. elegans* | BASIC FORM Celegans SingleSheet jsm 2 13 13 LogMethod.xlsx | 140426 C elegans.xlsx |
| Chickens | *Gallus gallus* | CHikcens Aggrey and Byerley and cleavage Data 2 14 2013.xlsx | 140426 Chickens.xlsx |
| Cows | *Bos taurus* | COW Basic Growth Data jsm 1 27 2013.xlsx | 140426 Cows.xlsx |
| Geese | *Anser anser* | BASIC FORM Geese SingleSheet jsm 2 14 2013 Log Method.xlsx | 140426 Geese.xlsx |
| Mice | *Mus musculus* | BASIC FORM MicePoiley M AKR jsm 1 28 2013 Log&LinearMethods.xlsx | 140426 Mice.xlsx |
| Quail | *Colinus virgianus* | BASIC FORM Quail SingleSheet jsm 2 16 2013.xlsx | 140426 Quail.xlsx |
| Rats | *Rattus norvegicus* | RAT Basic Growth Data jsm 2 14 2013.xlsx | 140426 Rats.xlsx |
| Turkey | *Meleagris gallopavo* | BASIC FORM TURKEYwithOvaposition SingleSheet jsm 2 8 2013.xlsx | 140426 Turkey.xlsx |
| Clams | *Merceneria mercenaria* | BASIC FORM LOG METHOD mercenaria 1 11 2013 v2 from Boys SingleSheet jsm 2 12 2013 | 140705 Mercenaria.xlsx |
| LOCATION: | | C:_0\My Papers\Growth_DRAFTS__final Excell Files_Basic Data 7 7 14 | C:_0\My Papers\Growth_DRAFTS__final Excell Files_Phils Analysis 7 7 14 |

#### TABLE A2
***Universal Growth Equation*.**
Parameters and goodness-of-fit metrics

| Organism | Genus species | ***Fit of the data to the rate of growth g=N*a^N^b (c=Cell Cycle Time**, in days)* | | | | | ***Fit of the data to growth N=∫[log(2)/c]*N*a^N^b*** |
|---|---|---|---|---|---|---|---|
| | | c | a | b | R^2 (linear fit) | R^2 (log fit) | R^2 (log fit) |
| Humans (♂) | *Homo sapiens* | 1.016 | 0.943 | 0.169 | 89.9% | 95.0% | 99.5% |
| Frogs | *Rana pipiens* | 0.070 | 0.925 | 0.269 | 57.0% | 82.3% | 97.3% |
| Nematodes | *C. elegans* | 0.011 | 0.946 | 0.680 | 66.7% | 91.6% | 99.8% |
| Chickens | *Gallus gallus* | 0.127 | 0.906 | 0.166 | 80.8% | 97.8% | 99.8% |
| Cows | *Bos taurus* | 0.921 | 0.882 | 0.128 | 76.8% | 95.1% | 100.0% |
| Geese | *Anser anser* | 0.048 | 0.898 | 0.170 | 77.0% | 92.9% | 97.1% |
| Mice | *Mus musculus* | 0.508 | 0.990 | 0.281 | 65.0% | 96.2% | 99.7% |
| Quail | *Colinus virgianus* | 0.292 | 0.795 | 0.126 | 76.5% | 95.3% | 94.2% |
| Rats | *Rattus norvegicus* | 0.417 | 0.950 | 0.197 | 66.5% | 90.6% | 99.5% |
| Clams | *M. mercenaria* | 0.033 | 0.962 | 0.275 | 79% | 91% | 99.5% |
| Turkeys | *Meleagris gallopavo* | 0.028 | 0.757 | 0.122 | 65.3% | 92.2% | 99.7% |

1.

Zebrafish (*Danio rerio*) and European sea bass (*Dicentrarchus labrax*) also show these features of growth (data not shown)
